# Single-Islet Proteomics Maps Pseudo-Temporal Islet Immune Responses and Dysfunction in Stage 1 Type 1 Diabetes

**DOI:** 10.1101/2025.11.10.687674

**Authors:** Shane Kelly, Soumyadeep Sarkar, Sarai M. Williams, An D. Fu, Elizabeth A. Butterworth, Tyler J. Sagendorf, Lorenz A. Nierves, Yumi Kwon, Xiaolu Li, Vladislav A. Petyuk, Jing Chen, Ernesto S. Nakayasu, Mark A. Atkinson, Rohit N. Kulkarni, Clayton E Mathews, Ying Zhu, Martha Campbell-Thompson, Wei-Jun Qian

**Affiliations:** Biological Sciences Division, Pacific Northwest National Laboratory, Richland, WA; Environmental Molecular Sciences Division, Pacific Northwest National Laboratory, Richland, WA; Department of Pathology, Immunology, and Laboratory Medicine, College of Medicine, University of Florida, Gainesville, FL; Department of Infectious Disease and Immunology, College of Veterinary Medicine, University of Florida, Gainesville, FL; Section of Islet Cell Biology and Regenerative Medicine, Joslin Diabetes Center and Department of Medicine, Beth Israel Deaconess Medical Center, Harvard Stem Cell Institute, Harvard Medical School, Boston, MA; Department of Proteomic and Genomic Technologies, Genentech Inc., 1 DNA Way, South San Francisco, 94080, USA

## Abstract

Progressive β-cell dysfunction precedes the onset of type 1 diabetes (T1D), yet the molecular mechanisms driving early T1D development remain poorly understood. Although single-cell RNA-sequencing has uncovered transcript-level changes in human islet cells, it offers limited insight into the heterogeneity of distinct islet microenvironments. Here, we applied a single-islet proteomics workflow to profile intra-donor islet heterogeneity in three stage 1 T1D cases with matched non-diabetic controls and define in situ protein signatures of pseudo-temporal islet dysfunction. Intra-donor analyses of ∼100 individual islets per donor revealed highly consistent proteomic patterns reflecting pseudo-time progression of islet immune responses and β-cell dysfunction. Several pathways, including extracellular matrix remodeling and mRNA processing, were identified as closely associated with progressive islet immune activation and loss of β-cell function. These findings provide robust proteome-wide evidence of the progression of islet dysfunction, offer a valuable resource for investigating early mechanisms of T1D pathogenesis— including novel candidates for functional studies—and underscore the utility of single-islet spatial proteomics for examining islet heterogeneity in T1D.

## Introduction

Type 1 diabetes (T1D) is a chronic autoimmune disease characterized by the selective destruction of insulin-producing β-cells within the islets of Langerhans, leading to insulin deficiency and chronic hyperglycemia^1,2^. A prolonged asymptomatic phase typically precedes clinical onset, marked by the presence of islet autoantibodies and a decline in β-cell function^3,4^. Longitudinal studies have revealed that defects in glucose-stimulated insulin secretion and first-phase insulin response may emerge 4-6 years before clinical diagnosis^5,6^. These findings highlight the importance of understanding molecular mechanisms during this preclinical stage to develop therapeutic strategies that can prevent or delay disease progression.

Mouse models of diabetes and cytokine-treated in vitro systems used over the past 40 years have provided insights into β-cell stress and immune signaling pathways^7,8^. However, these findings often fail to fully capture the complexity and heterogeneity of human T1D pathogenesis. Recent research has increasingly shifted to investigating human pancreatic tissues obtained from organ donors, utilizing advanced ‘omics techniques to uncover insights into disease mechanisms. For instance, multiple studies have employed single-cell multi-omics approaches to produce comprehensive cell type-specific atlases of pancreatic islet cells from non-diabetic, AAb+, recent-onset T1D and long-standing T1D donors^9–11^. These studies uncovered cell–type–specific expression changes, with β-cells exhibiting the strongest immune-related alterations and revealed a potential broader remodeling of the islet and exocrine microenvironments in T1D. Similarly, a whole pancreas tissue gene expression profiling study of patients with recently diagnosed T1D highlights the role of the exocrine pancreas with higher expression of digestive enzyme genes (such as *AMY2A*, *AMY2B*, *CELA3A*, and *CELA2B*) in T1D compared to non-diabetic controls^12^. The importance of tissue microenvironments is underscored by recent spatial imaging using imaging mass cytometry (IMC)^13,14^ and CODEX^15^, which allow simultaneous imaging of dozens of protein markers in pancreatic tissue from T1D, autoantibody (AAb) positive (at risk), and non-diabetic donors, revealing alterations in islet architecture, increased immune cell infiltration, selective β-cell marker loss, and cytotoxic T cell recruitment preceding full β-cell destruction. Other spatial studies, including multiplexed immunofluorescence ^16^, extracellular matrix (ECM) mapping^17^, and functional slice physiology^18^, analyzing T1D donors at various stages of the disease, demonstrated that insulitic lesions and immune aggregates are often confined to specific islets, while other islets remain unaffected. However, our recent results have demonstrated that β-cell dysfunction is not associated with insulitis^18^. This emphasizes the significant gaps that remain in our understanding of the molecular mechanisms driving the progression of T1D. These limitations arise in part from the heterogeneous nature of the disease^19^ and the cross-sectional study design or “snapshot” nature of data obtained from cadaveric donor samples. Furthermore, while omics studies on human pancreatic tissues have primarily relied on transcriptomics, few investigations have applied proteomics^20,21^, even though transcript levels often fail to predict protein abundance, and proteins more directly reflect tissue functional states.

Another notable feature of T1D is the observed intra-donor islet heterogeneity and lobularity within pancreatic tissue from individual donors^22,23^. This intra-donor islet heterogeneity represents a valuable, yet underutilized, opportunity to identify pseudo-temporal disease progression information and its underlying pathways through spatial and molecular analyses at the level of individual islets—even without longitudinal sampling. In this study, we employed single-islet proteomics to examine intra-donor islet heterogeneity and early T1D pathogenesis. Using laser microdissection (LMD) and nanoPOTS-based proteomics workflow^24^, we profiled approximately 100 individual islet sections per donor, including donors with stage 1 T1D (multiple AAb positive; mAAb+) and non-diabetic (ND) controls. Multiplex immunohistochemistry (mIHC) was used to annotate immune infiltration (CD3⁺ cells) and β-cell content (INS⁺ cells). Through unsupervised clustering and machine learning analyses, we uncovered distinct trajectories of islet immune response states and β-cell function within stage 1 T1D donors, representing a “pseudo-temporal” progression of islet dysfunction. These analyses identified two key molecular signatures: an Islet Immune Response Signature and a β-Cell Profile, each comprising a panel of ∼40 proteins.

Our findings provide protein-level molecular insights into the activated islet immune response, β-cell function, and ECM remodeling within individual stage 1 T1D donors. These results corroborate and expand prior transcriptomics and single-cell multi-omics studies and captured molecular responses to pro-inflammatory cytokines (e.g., IL-1β, IFNγ) in vitro^25^. Furthermore, our data highlight ECM remodeling and mRNA processing as potential pathways that contribute to the loss of β-cell function and progression of T1D, to offer new perspectives on the interplay between tissue context as well as immune cell and endocrine cell dynamics in T1D pathogenesis.

## Results

### Mapping intra-donor islet heterogeneity by single-islet proteomics

Our approach focuses on resolving intra-donor islet heterogeneity at the proteome level by profiling ∼100 islets obtained from each of three stage 1 T1D donors (mAAb+) and ∼50 islets each from three age, sex, and race-matched ND donors ((**Fig.1A & Table 1**). We propose that such heterogeneity within a single donor, before clinical onset, represents a pseudo-temporal trajectory of islet dysfunction that mirrors early disease progression. Multiple fresh frozen tissue sections from each donor were obtained from nPOD. Adjacent tissue sections were subjected to islet phenotyping (mIHC) for detection of insulin (INS: β-cells), glucagon (GCG: α-cells), and cell differentiation marker 3 (CD3: T cells) or LMD for isolation of individual islet sections. The isolated islets from stage 1 T1D donors were phenotype-matched based on mIHC CD3 status. The individual islets were then processed by nanoPOTS and analyzed by LC-MS/MS with label-free quantification (**Fig. 1B**). An average of ∼5800 proteins were identified and quantified per donor (**Extended Data Fig.1A, Supplementary File 1**). For all proteomic data from single islet sections, the data were log2 transformed, normalized, and batch-corrected for intra-donor tissue blocks (**Extended Data Fig.1A**). The robustness of the analytic workflow was assessed based on data from randomly distributed reference samples as processing replicates on the nanoPOTS chip. Pearson correlation coefficients >0.96 were observed between any pairs of reference samples, while correlation values between 0.87-0.95 for pairs between single islet samples (**Extended Data Fig.1B**). We also examined the quality of LMD isolation of islet sections by analyzing the cell type composition information based on selected cell-type specific markers. As expected, the proteomics data indicate that α- and β-cells are consistently shown to be the dominant cell types across all six donors, with only a minimal contribution of other cell types (<2% based on proteomic estimation) (**Fig.1C)**. Next, we also plotted the abundances of insulin) and glucagon for individual islets across the six donors to examine intra-donor variances with insulin or glucagon levels (**Fig. 1D and E**). A cutoff of insulin level 4-fold lower than the median (-2 in log2 scale) was used to identify low insulin islets. We observed 7 and 16 low-insulin islets from cases 6450 and 6521, respectively. This is consistent with our anticipation of potential loss of β-cells in stage 1 T1D, with triple AAb+ case 6521 having the greatest number of low-insulin islets. For GCG, most islets have abundances within 4-fold of the median, with only 9 islets having levels lower than that range. These low-GCG islets were not associated with any donor type. Upon plotting the GCG vs INS abundances for individual islets, we observe that INS abundances are more varied across the islets than GCG, with GCG abundances in low-insulin islets being close to the median (<-2 in log2 scale) (**Fig. 1F**). Overall, these results establish a robust single-islet proteomics framework capable of capturing intra-donor heterogeneity, providing a molecular window into the earliest events of T1D pathogenesis.

**Figure 1.**
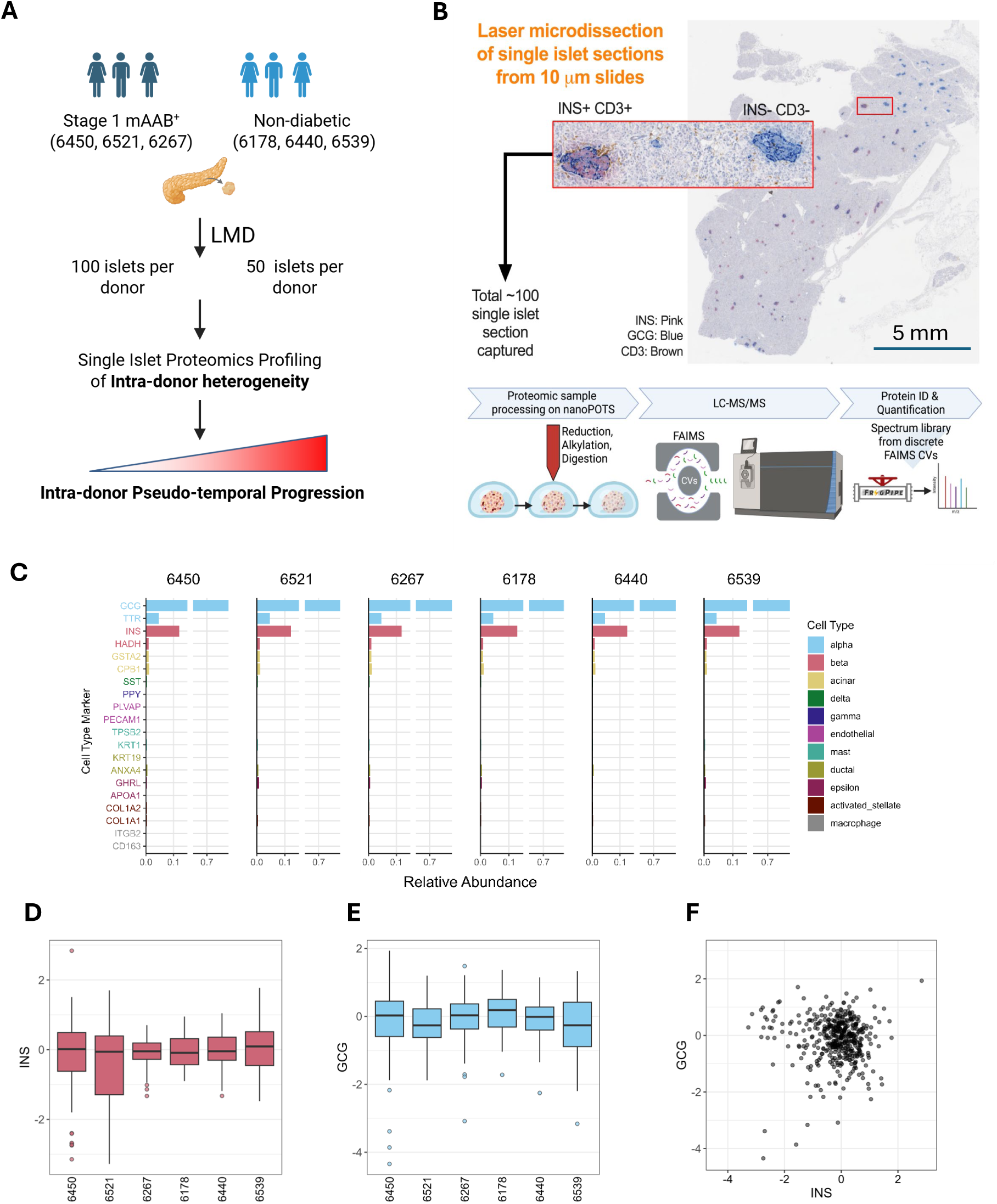
Experimental workflow and quality control for single-islet proteomics profiling. **A.** Overall experimental design. Tissue sections (3-5) from each age, sex, and race-matched nPOD donor were obtained for islet selection and analysis of intra-donor islet heterogeneity. **B.** Experimental workflow. Adjacent tissue sections were mIHC stained for INS, GCG, and CD3 to inform islet selection prior to LMD, nanoPOTS sample processing, and LC-MS/MS data acquisition. **C.** Quantitative analysis of cell-type protein markers based on the intensity-based absolute quantification (iBAQ) information. The markers were selected based on scRNA-seq data. Relative abundance is the non-log-transformed intensity relative to the summed intensity across all cell type markers. **D.** Median-centered log2 insulin intensity distribution across donors. **E.** Median-centered log2 glucagon intensity distribution across donors. **F.** Comparison of insulin and glucagon abundance in LMD isolated islets.

**Table 1:** Age and sex matched donor information and clinical description obtained from nPOD.

| nPOD ID | Case Type | Insulinitis Status | AAb detected | Race | Age | Sex | HbA1C | BMI | HLA |  |  |  |  |
| --- | --- | --- | --- | --- | --- | --- | --- | --- | --- | --- | --- | --- | --- |
|  |  |  |  |  |  |  |  |  | A | B | DRB1 | DQA1 | DQB |
| 6450 | mAAb+ | Yes | GADA+ ZnT8A+ | Caucasian | 22.0 | F | 5.7 | 24.4 | 01:01 | 08:01 | 03:01 | 05:01 | 02:01 |
|  |  |  |  |  |  |  |  |  | 33:01 | 14:02 | 03:01 | 05:01 | 02:01 |
| 6521 |  |  | GADA+ IA-2A+ ZnT8A+ | Hispanic/Latino | 19.8 | M | 5.8 | 24.1 | 02:01 | 08:01 | 03:01 | 05:01 | 02:01 |
|  |  |  |  |  |  |  |  |  | 02:01 | 44:03 | 04:02 | 03:01 | 03:02 |
| 6267 |  |  | GADA+ IA-2A+ | Caucasian | 23.0 | F | 5.0 | 23.5 | 01:01 | 39:01 | 04:04 | 03:01 | 03:02 |
|  |  |  |  |  |  |  |  |  | 11:01 | 60:01 | 04:04 | 03:01 | 03:02 |
| 6178 | No Diabetes | - | - | Caucasian | 24.5 | F | 5.0 | 27.5 | 02:01 | 27:01 | 04:02 | 03:01 | 03:02 |
|  |  |  |  |  |  |  |  |  | 24:01 | 44:01 | 15:01 | 01:02 | 06:02 |
| 6440 |  |  |  | Caucasian | 15.7 | F | 5.0 | 20.5 | 02:01 | 14:02 | 01:02 | 01:01 | 05:01 |
|  |  |  |  |  |  |  |  |  | 33:01 | 15:01 | 13:01 | 01:03 | 06:03 |
| 6539 |  |  |  | Hispanic/Latino | 24.0 | M | 5.7 | 19.3 | 02:01 | 27:02 | 04:03 | 03:01 | 03:02 |
|  |  |  |  |  |  |  |  |  | 11:01 | 49:01 | 13:02 | 01:02 | 06:04 |

### Clustering analysis identifies protein modules associated with CD3 phenotype

To more closely examine intra-donor heterogeneity, we applied the Weighted Gene Co-expression Network Analysis (WGCNA) as a clustering approach to the single-islet proteomics data from each donor to identify protein modules (i.e., eigenproteins) that are significantly associated with a given phenotype, including CD3 status and INS intensity data (from proteomics). **Fig. 2A** shows the eigenprotein-phenotype association plots for all three stage 1 T1D donors with each module assigned with a color code. One or multiple modules were identified with significant association (adjusted p-value <0.05) with CD3 or INS intensity for each of the stage 1 T1D donors. To identify underlying pathways associated with these modules, we performed an Over-Representation Analysis (ORA) based on Gene Ontology Biological Process (GO:BP) terms for each module.

**Figure 2.**
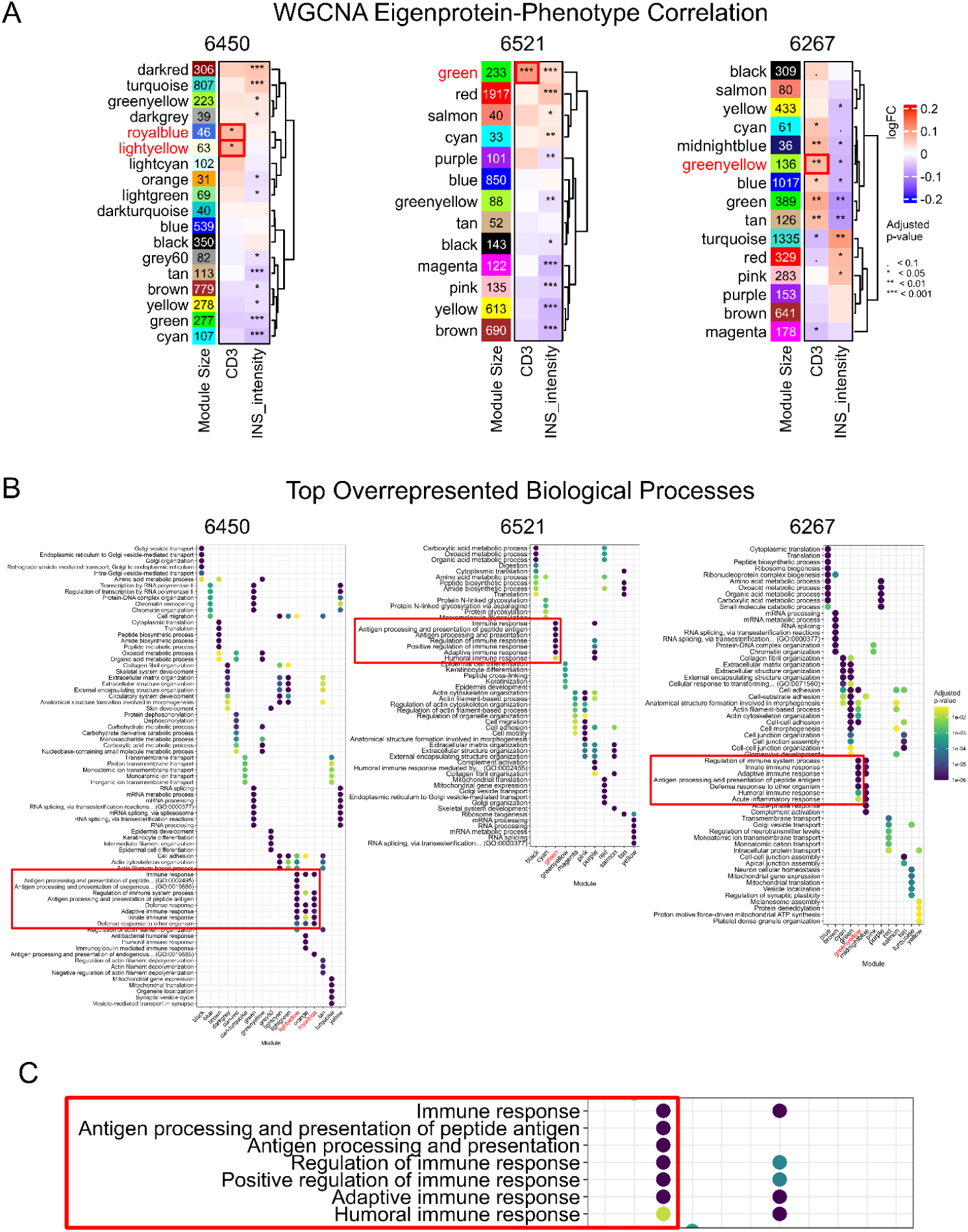
Identification of immune-associated protein modules by clustering and over-representation analysis. **A.** WGCNA identified protein modules correlated with CD3 status and insulin (INS) intensity for stage 1 T1D donors with case ID 6450, 6521, & 6267. The red boxes on the heatmap highlight modules with positive correlation with the CD3 phenotyping data. **B.** Top 5 overrepresented biological processes for each of the WGCNA protein modules with significant enrichment in at least one GO:BP term, with color scale representing the adjusted p-value. The red box enclosing the pathway terms highlights modules associated with CD3 are also highly enriched with pathways associated with innate and adaptive immune response. **C.** Highlighted immune response pathways enriched in the green module of donor 6521.

Initially, we focused on modules that are significantly associated with CD3 to identify immune-mediated biological processes in the islets. Notably, for case 6521, a “green” module with 233 proteins was identified with the most significant association with CD3 (**Fig. 2A)**. This module also displayed clear enrichment of biological processes associated with adaptive and innate immune responses (**Fig. 2B and C, Supplementary File 2**). Similarly, protein modules significantly associated with CD3 were also enriched with adaptive and innate immune response processes for case 6450 and 6267 (**Fig. 2B**). Antigen processing and presentation pathways are particularly notable because it was consistently observed in all three stage 1 donors, but not in any ND controls (**Extended Data Fig. 2**). Overall, these findings indicate that intra-donor proteomic heterogeneity reflects localized immune engagement within the islet microenvironment, highlighting early activation of antigen presentation and innate–adaptive crosstalk within stage 1 T1D individuals.

### An Islet Immune Response Signature capturing immune-mediated pseudo-temporal progression

Since CD3 staining only provides limited binary information about T-cell infiltration, we aim to identify a panel of protein markers as an islet immune response signature (IIRS). Conceptually, a proteomic panel would offer a more comprehensive and reliable representation of islet immune response compared to relying solely on the single pan-T cell marker, CD3. The IIRS could serve as a robust tool for characterizing immune-mediated pseudo-temporal progression across islets within each donor. To pursue this, proteins from all modules from WGCNA with significant enrichment of immune response pathways across the islets of stage 1 T1D donors were combined to provide a list of 329 candidate proteins. These proteins were then subjected to a random forest machine learning model (**Extended Data Fig. 3A**) that allow us to down select a final list of the top 40 ranked proteins as IIRS.

**Fig. 3A** shows the abundance profile of the 40 IIRS proteins across individual islets from stage 1 T1D and ND donors. The relative abundance level of IIRS represents the level of immune-mediated response from islet cells, especially β-cells, at the individual islet level. The proteins are ranked based on feature importance determined by the random forest model. Within each stage 1 T1D donor, a pronounced and coherent abundance shift is observed with a portion of individual islets displaying upregulated (red) features, reflecting a pseudo-temporal progression of the islet immune response. In contrast, no such abundance shift was observed in ND controls. Notably, the observed magnitudes of islet immune response among stage 1 T1D donors appear to align with their AAb status. For instance, donor 6521, classified as triple AAb+, shows the highest percentage of islets with elevated IIRS protein abundances, indicating more advanced immune-mediated progression compared to other two stage 1 donors. **Fig. 3B** further highlights the abundance ranges of IIRS for each donor. Stage 1 T1D donors exhibit substantially larger abundance ranges compared to ND donors, reinforcing the hypothesis that significant intra-donor progression in islet dysfunction can be captured at the single-islet level in stage 1 T1D. This aligns with the proposed model of immune-mediated progression of islet dysregulation being detectable even in early stages prior to clinical onset. **Fig. 3C** displays a pseudo-temporal map of individual islets from all six donors based on the relative abundance of the IIRS. The overall distribution can be broadly categorized into clusters of low, moderate, and high inflammation. When examining individual donors (**Fig. 3D**), each stage 1 donor exhibits a distinct cluster of highly inflamed islets, where such clusters are absent in ND donors.

**Figure 3.**
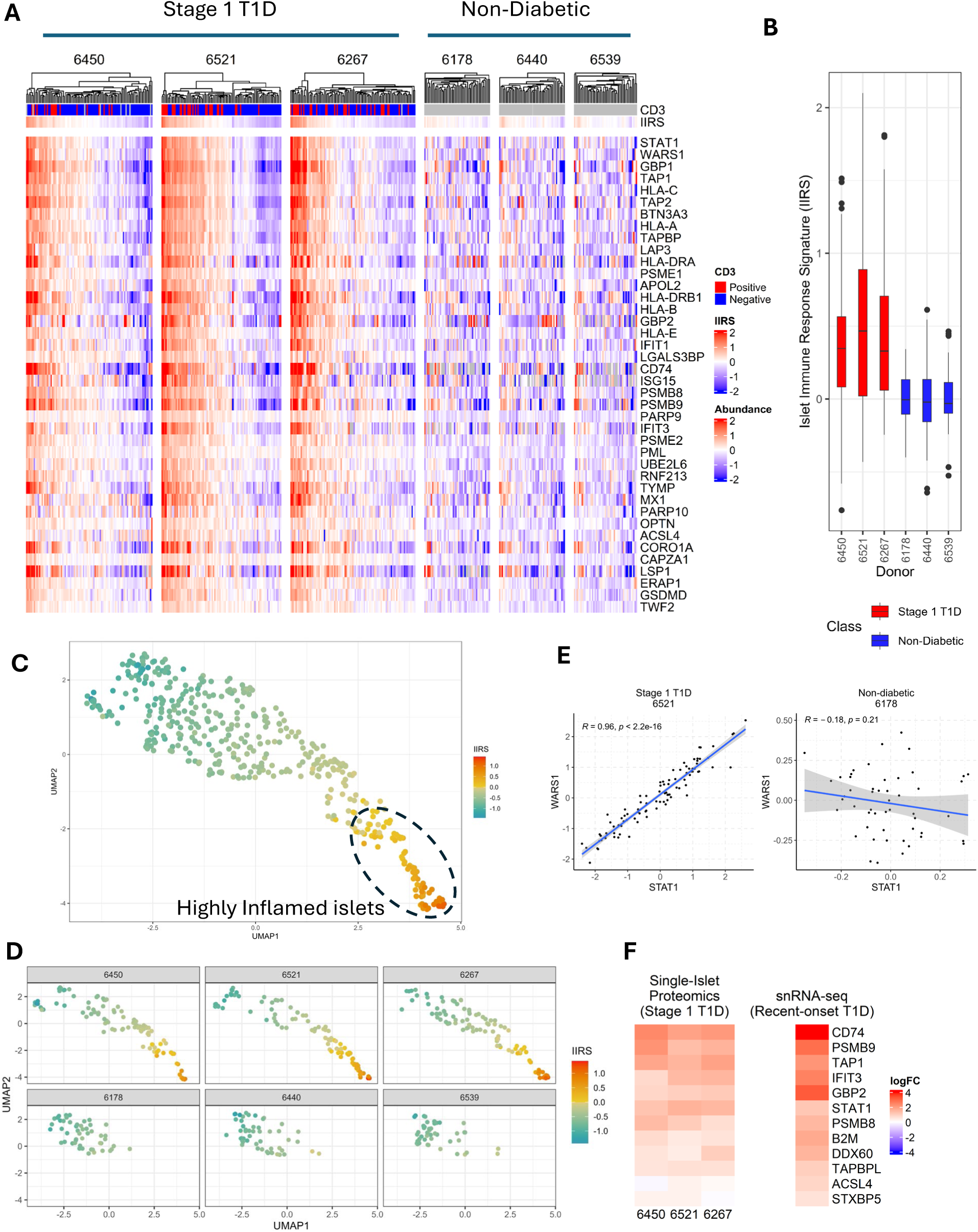
An Islet Immune Response Signature (IIRS) with a 40-protein panel. **A.** Heatmap of protein abundances of the 40 IIRS proteins for individual islet samples across each donor. Protein abundances are log2-transformed and median-centered. Note that the median-centering across the donors was performed under the assumption that the islets exhibiting lower half of IIRS expression levels in stage 1 T1D were comparable to ND control islets. Proteins are ranked in descending order according to feature importance derived from random forest models consolidated through robust rank aggregation. CD3 binary status for each islet is represented in the top, with red as positive and blue as negative. Additionally, the overall IIRS relative abundance for each islet is represented as the median abundance of the 40-protein panel. **B.** Box and whisker plot, representing the dynamic range of the IIRS abundance per donor. **C.** Pseudo-temporal map of islets from all donors based on IIRS abundance. **D.** Donor level islet maps based on IIRS. **E.** Contrasting co-regulation of STAT1 vs WARS1 between stage 1 and ND donors. *R* denotes the Pearson correlation coefficient, and *p* is the corresponding p-value assessing the statistical significance of the correlation. **F.** Overlap of IIRS proteins with differentially expressed genes identified by scRNA-seq. Left: Log2FC (slope) based on a linear model against IIRS for each stage 1 donors in this study. Right: Log2FC between recent-onset T1D and ND patients. Genes are those that are included in the IIRS and also significantly (adjusted p-value <0.1) differentially expressed in β-cells by scRNA-seq.

Moreover, significant co-regulation among the members of IIRS is apparent in islets of stage 1-T1D donors but is substantially less correlated in islets of ND donors, as indicated by the correlation heatmaps (**Extended Data Fig. 3B**). **Fig. 3E** further illustrates high correlation (*R=0.95*) between two members (STAT1 and WARS1) of the IIRS only in stage 1. Upon further examination of the proteins comprising the IIRS, we identified a combination of well-established markers and lesser-known proteins in the context of T1D. Key components include HLA-A, HLA-B, HLA-C, HLA-E, HLA-DRA, HLA-DRB1, CD74, TAP1, TAP2, TAPBP, PSMB8, PSMB9, PSME1, PSME2, and ERAP1, which are integral to the major histocompatibility complex (MHC) class I and II machinery, as well as antigen processing. Notably, increased HLA-ABC expression was confirmed in inflamed islets from both mAAb+ and T1D donors in the recent CODEX imaging study^15^. The expression of these proteins is also markedly upregulated in islets exposed to IFN-γ and other cytokines, highlighting their connection to autoimmune-mediated targeting of β-cells in T1D^20,26^. Furthermore, we observed a significant enrichment of proteins involved in interferon-mediated immune response, including STAT1, GBP1/2, ISG15, IFIT1/3, MX1, PML, and UBE2L6, and WARS1. These findings align closely with observations of overexpression of interferon-stimulated genes in insulitic islets from living patients with recent-onset T1D^27^. Our data are also consistent with the recent findings from single cell or single nuclear RNA-Seq profiling^11^ where differential gene expression was identified when comparing recent-onset T1D cases to ND controls (**Fig. 3F**). Notably, unlike in T1D, the previous study only detected a few individual genes with significant expression changes in β-cells of stage 1 mAAb+ compared to ND controls^11^, underscoring the advantage of intra-donor analysis in revealing robust activation of immune response pathways in stage 1 donors.

In addition to these established disease-associated proteins, novel IIRS proteins with limited established association in the context of T1D were identified. For example, PARP10 and OPTN are known to regulate inflammatory signaling and cytokine response^28^, whereas LAP3 has been implicated in the MHC class I antigen presentation pathway^29^; however, no direct studies have established their association or roles in T1D. The IIRS also captured programmed cell death proteins like GSDMD^30^ (pyroptosis) and ACSL4^30^ (ferroptosis), which have been associated with β-cell death in in vivo models, providing the first direct evidence of their involvement in T1D development in human patients. Another set of proteins that has limited evidence in the context of T1D are TWF2, CORO1A, CAPZA1, and LSP1, which play a role in regulating cytoskeleton dynamics and immune cell migration. Finally, IIRS proteins like RNF213 and APOL2^31^ can be broadly classified as β-cell intrinsic modulators mainly involved in promoting β-cell dysfunction, whereas PARP9, LGALS3BP, TYMP, and BTN3A3 can be extrinsic modulators, increasing islet inflammation/autoimmunity. Overall, the intra-donor analysis, along with the IIRS, reveals a clear trajectory of islet inflammation within the stage 1 T1D donors, and the IIRS provides a more robust estimate of islet immune response than the single CD3 marker by providing a more comprehensive representation of the activation of islet immune response pathways.

### Pathways underlying the progression of the islet immune response in stage 1 T1D

To gain further insights into the molecular changes that contribute to or are affected by islet immune response, we applied linear models (LIMMA) to test the relationships of each protein with the IIRS within each donor (**Supplementary File 3**). The volcano plots in **Fig. 4A** summarize the significant results with IIRS as the predictor of protein abundance, where the x-axis is the log_2_(fold-change) (i.e., slope of each fitted line), and the y-axis is the –log_10_(adjusted p-value). Each stage 1 T1D donor showed much larger fold-changes and more significant correlation with IIRS than the ND controls, indicating the IIRS as a “driver” for islet protein abundances. Next, a CAMERA-PR-based gene set enrichment analysis^32^ was conducted on the above results to identify significant pathways associated with IIRS across the donors. **Fig. 4B** shows a curated version of the top 30 biological processes from this analysis. Most of the represented gene sets show a clear donor type association. For example, immune-related pathways, including antigen processing and presentation, humoral immune response, T cell-mediated cytotoxicity, adaptive immune response, lymphocyte-mediated immunity, interleukin-10 production, and response to Type I and II interferon, all have substantially higher Z-scores (i.e., slope of correlation) in stage 1 T1D compared to ND donors, indicating stronger associations of immune response-associated gene sets with the IIRS in stage 1 T1D donors. Moreover, nearly all observed immune-related pathways were also reported among the significantly enriched pathways in T1D β-cells in a previous report of an integrated single-cell RNA-sequencing (scRNA-seq) map^10^. In contrast, ECM-related gene sets such as ECM organization, integrin-mediated signaling, cell-matrix adhesion, and collagen fibril organization were observed with lower Z-scores (or lower association) with IIRS in stage 1 T1D compared to ND donors, presenting evidence of ECM dysregulation in stage 1 T1D donors. To further support the ECM observations, both the hyaluronan metabolic and glycosaminoglycan catabolic processes displayed higher associations with IIRS in the stage 1 donors, suggesting both processes are dysregulated in connection with ECM remodeling. In terms of negatively associated pathways, mitochondrial translation and ER-Golgi-related pathways emerged.

**Figure 4.**
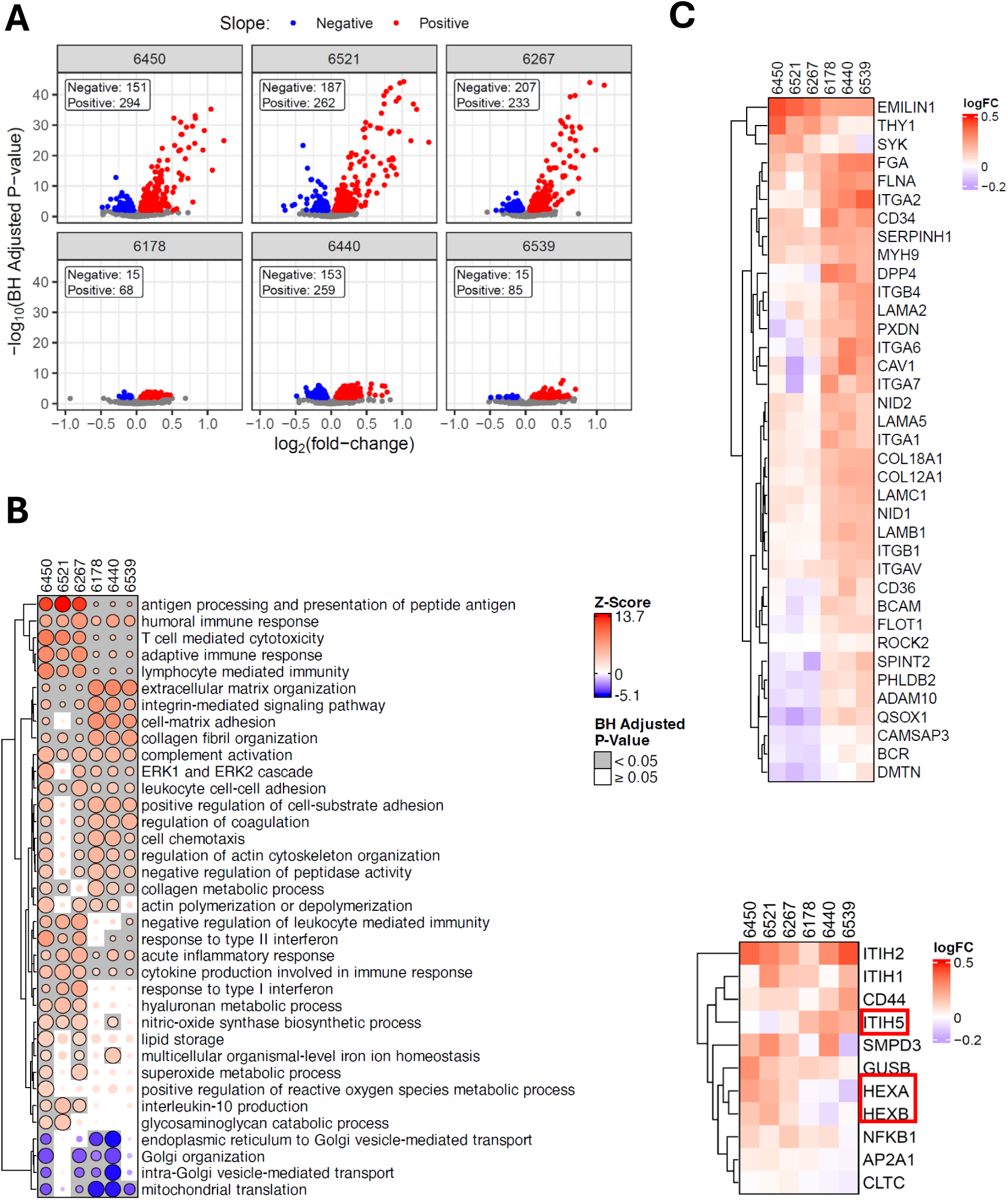
Pathways associated with the immune-mediated pseudo-temporal progression. **A.** Volcano plots of LIMMA analysis with the IIRS as the predictor of protein abundance for individual donors. The x-axis is the log_2_(fold-change) (i.e., slope of each fitted line), and the y-axis is the –log_10_(adjusted p-value). Red indicates a positive slope and blue indicates a negative slope in the protein-IIRS correlation. Colored points indicate adjusted p-value <0.01. **B.** Bubble heatmaps summarizing the top GO Biological Process terms from the CAMERA-PR analysis applied to LIMMA results against IIRS. The bubble sizes are scaled by the –log_10_(adjusted p-value) and then re-scaled relative to the most significant result within each gene set to better observe relationships across donors. The colors are determined by the z-scores, which were derived from the LIMMA logFC (i.e., slope). **C. & D.** Heatmaps for proteins in ECM organization and Hyaluronan metabolic process, respectively, showing the log2FC of each protein across individual donors.

Further zooming in on the individual pathways to gain a protein-level view (**Fig. 4C**), we observe that proteins associated with ECM display significant differences between stage 1 T1D and ND donors. Interestingly, these proteins generally have larger fold-changes in relation to IIRS within ND donors, suggesting that ECM regulation is closely associated with normal cellular immune response. However, the observed fold-changes are substantially lower in stage 1 T1D donors as compared to ND donors, providing evidence of ECM dysregulation in stage 1 T1D. Other ECM-related pathways include hyaluronan metabolic process and glycosaminoglycan catabolic process, both of which are closely intertwined with the organization of ECM (**Fig. 4D, Extended Data Fig. 4A**). In both pathways, several proteins were shown with distinct levels of association with IIRS in stage 1 vs ND donors. Specifically, ITIH5^33^, a hyaluronan (HA) modifier/stabilizer, has a negative slope, whereas HEXA/B^34^, responsible for finishing breaking down HAs, has a positive slope with IIRS in stage 1 donors. These observations support a loss of ECM under islet inflammation. Finally, we observed a distinct association of the interleukin-10 (IL-10) production pathway with IIRS in stage 1 T1D (**Extended Data Fig. 4B**). Given the protective and anti-inflammatory role of IL-10 in T1D, we observed a marked positive IIRS association of the IFN-stimulated gene (ISG15), a known upstream regulator of IL-10. In contrast, thrombospondin-1 (THBS1) and CD46 exhibited no IIRS association in stage 1 T1D donors (**Extended Data Fig. 4B**). Together, these patterns point to potential insufficient IL-10 activity to resolve inflammation.

Complementary analyses using Ingenuity Pathway Analysis (IPA) based on the list of significant proteins correlated with IIRS converge on the same conclusion that antigen presentation, interferon signaling, and IL-10 signaling are among the top pathways (**Extended Data Fig. 4C).** Moreover, Interferon alpha family is predicted to be one of the top upstream regulators by IPA (**Extended Data Fig. 4D**).

### Progressive loss of β-cell function in stage 1 T1D

Beyond the islet immune response, one key question is whether there is molecular evidence of progression in the loss of β-cell function or other phenotypes within individual stage 1 T1D donors. As discussed earlier, low-insulin islets were observed within several of the stage 1 T1D cases, indicating the potential loss of β-cell mass or function. However, one challenge in analyzing the loss of β-cell function is the lack of reliable functional biomarkers. Alternatively, we seek to identify proteins that can potentially serve as a signature panel for β-cell function or identity. To explore this, we conducted a fine-grained clustering analysis across all data to identify functionally clustered protein modules. Among the final 75 clusters (**Supplementary File 4**), we confirmed several clusters (e.g., the TAP-binding) were strongly correlated with the IIRS, and a cluster consisting of the constitutive subunits (PSMB5/6/7) of the proteasome was anti-correlated with IIRS^35^ (**Extended Data Fig. 5A and B**). Interestingly, a cluster of proteins (**Extended Data Fig. 5C)** mainly consisting of β-cell-specific markers (e.g., ENTPD3) also shows substantially larger intra-donor heterogeneity in stage 1 T1D compared to ND controls, suggesting β-cell-dysfunction present in stage 1 T1D.

We then sought to identify a panel of proteins as a <u>β</u>-<u>c</u>ell <u>p</u>rofile (BCP) by selecting the top 41 proteins with the highest average correlation coefficient against INS and ENTPD3. **Fig. 5A** shows the heatmap of the 41 proteins of the BCP across individual islets of stage 1 T1D and ND donors.

**Figure 5.**
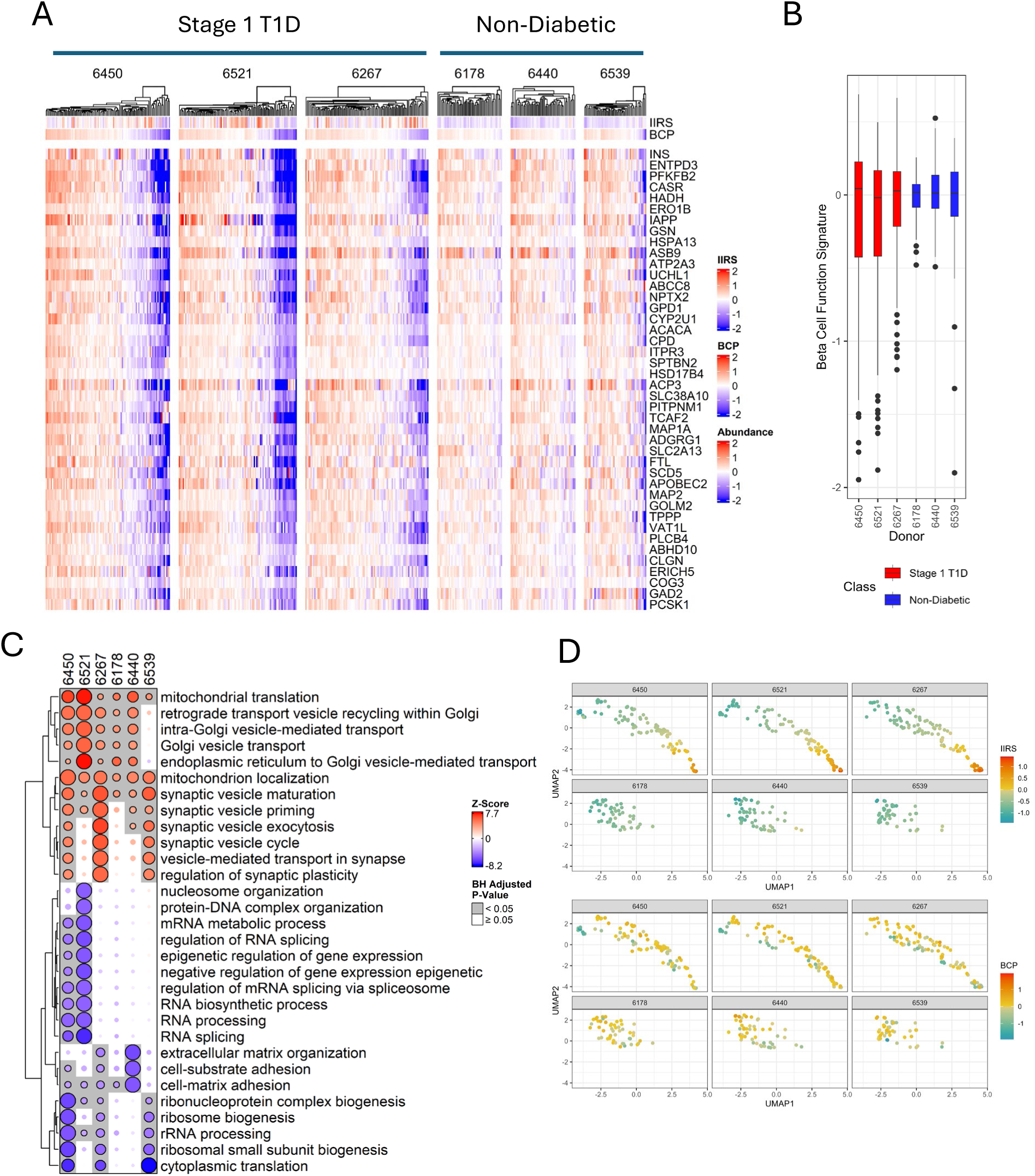
The β-Cell Profile (BCP) with 41-proteins. **A.** Heatmap of protein abundances of the 41 BCP proteins for individual islet samples across each donor. Proteins are log2-transformed and median-centered across all samples. Proteins are ranked by mean correlation with INS and ENTPD3. **B.** Box and whisker plot, representing the dynamic range of the BCP abundance per donor. **C.** Bubble heatmaps summarizing the top GO Biological Process terms from the CAMERA-PR analysis applied to LIMMA results against BCP. The bubble sizes are scaled by the – log_10_(adjusted p-value) and then re-scaled relative to the most significant result within each gene set to better observe relationships across donors. The colors are determined by the z-scores, which were derived from the LIMMA logFC (i.e., slope). **D.** Comparison of IIRS and BCP abundance at the single islet level. Top: Pseudo-temporal map of islets from each donor, arranged by IIRS abundance. Bottom: The same map visualized according to BCP abundance.

Among these, several are recognized β-cell markers or ranked among the most β-cell-enriched genes identified in single-cell transcriptomic analyses^36^, including IAPP, NPTX2, PCSK1, ERO1B, HADH, and UCHL1. In all stage 1 T1D donors, a markedly greater proportion of islets exhibited low BCP expression or diminished insulin levels compared to ND controls, although islets with low BCP expression were also present in each control subject (**Fig. 5B**). Reduced BCP expression in these islets suggests a loss of β-cell mass, function, or identity in the affected patients.

A LIMMA analysis was also applied to test the linear relationships of each protein with BCP for each donor to identify proteins and pathways associated with the loss of β-cell function. Substantially larger numbers of proteins were shown to be significant in stage 1 donors than in ND donors based on the volcano plots (**Extended Data Fig. 6, Supplementary File 5**). Next, a CAMERA-PR-based gene set enrichment analysis was conducted to identify pathways associated with BCP across the donors (**Fig. 5C**). Notably, mitochondrial translation and gene expression were observed to have the highest positive correlation with BCP, which is not surprising in that pancreatic β-cells are known to have especially high mitochondrial content and metabolic activity, and the loss of mitochondrial translation would correlate with the loss of β-cell function^37^. Other pathways positively correlating with BCP are related to hormone secretion (e.g., synaptic vesicle cycle, Golgi vesicle transport). It should be noted that these pathways positively correlating with BCP are generally not differentiating between stage 1 T1D and control cases, presumably reflecting a common degree of intra-donor heterogeneity in islet β-cell mass across all donors.

Among the pathways showing the most significant negative correlation with BCP (**Fig. 5C**), several related to mRNA processing and RNA-splicing were prominently enriched in two of three stage 1 T1D cases, both of which containing islets with depleted insulin levels (**Fig.1D**). Examining the above mRNA processing cluster (**Extended Data Fig. 7A**), high expression levels of core components of the mRNA splicing machinery (SF3B1, SF3B2, SRSF9, SNRNP70, SFPQ, SMC3, SMC1A, PNN, ACIN1, THRAP3, SART1, SRSF9, IK, ZC3H14, SAFB, MATR3, RBMX) are most notable in islets with low insulin levels in donors 6450 and 6521, suggesting that altered mRNA processing is strongly linked with the loss of β-cell function^38^.

Another group of pathways exhibited significant negative correlation with BCP is again related to ECM remodeling. Like the observations with IIRS, the ECM-associated gene sets show negative correlation with BCP in both stage 1 T1D and ND donors. Further examining the abundance profiles of several clusters associated with ECM remodeling (**Extended Data Fig. 7B-D**), the abundances of these proteins are significantly higher in the islets with low expression of BCP, suggesting a potential significant role of ECM remodeling in the demise of β-cells.

Finally, we note that the abundance profiles of BCP and IIRS are weakly correlated (Pearson correlation r = 0.13) in stage 1 T1D, consistent with our recent report indicating that β-cell dysfunction can occur independently of insulitis in T1D pathogenesis^18^. The pseudo-temporal islet maps displayed in either IIRS or BCP expression levels (**Fig. 5D)** further illustrates that islets with low BCP expression do not align with pronounced islet inflammation but rather with either minimal or variable IIRS expression.

## DISCUSSION

Although decades of clinical research have documented progressive loss of β-cell function prior to the clinical onset of T1D, the molecular mechanisms driving early islet dysfunction remain poorly defined. Recent scRNA-seq studies have begun to reveal gene-level changes in islet cells^9,10^; however, our knowledge on the protein-level remains limited. Advanced spatial techniques such as imaging mass cytometry and CODEX have revealed structural and cellular changes in T1D islets, including immune cell infiltration and alterations in islet architecture^13,15^. Nevertheless, these methods are limited by their targeted marker panels (typically around 40-50 markers), resulting in a lack of comprehensive proteome-wide data from human T1D islets. Additionally, progress in understanding T1D pathogenesis is hindered by the limited access to human pancreatic tissue samples, especially donors with multiple autoantibodies and high-risk HLA alleles, and the heterogeneity of the disease.

In this work, we applied a spatial single-islet proteomics workflow to deeply profile intra-donor islet heterogeneity and identify proteome-level signals driving pseudo-temporal progression of islet dysfunction. Our approach combined mIHC with LMD thereby preserving spatial context to enable high-resolution in situ single-islet proteomic analysis of ∼100 individual islets per donor. While previous studies have observed activated islet inflammation in recent-onset or established T1D donors^11,15^, our intra-donor single islet analyses demonstrated clear and compelling evidence of activated islet immune response and β-cell dysfunction in all three stage 1 T1D cases. These detailed analyses allow us to define IIRS and BCP and to identify clusters of pathways linked to both the progression of islet immune response and the loss of β-cell function.

The most compelling pseudo-temporal pattern of progression in stage 1 T1D cases was observed as the islet immune response pathway highlighted by the 40 IIRS proteins (**Fig. 3**). Most of these proteins are known players within the antigen presentation or innate/adaptive immune response pathways, and approximately 50% have previously been reported as major contributors to T1D progression based on transcriptome studies^27^. We acknowledge that intra-donor pseudo-temporal progression of islet inflammation was recently reported in one stage 1 T1D case (6267)^15^, which is also included among the three stage 1 cases analyzed here. Our study not only corroborates these earlier findings but extends them by providing direct proteomic evidence demonstrating the involvement of both innate and adaptive immune responses in the progression of islet immune activation across all three stage 1 T1D cases examined.

A subset of IIRS proteins can be considered novel or understudied in T1D. Among them, PARP10, pyroptosis executor GSDMD, and OPTN stand out as the least established in T1D. Pyroptosis^39^ and ADP-ribosylation^40^ have been shown to play a role in β-cell death, and intriguingly, inflammasome activation connects these molecules to a common mechanistic pathway. GSDMD, a pore-forming executor of pyroptosis^41^, is cleaved by NLRP3 inflammasome, which is primed by NF-κB/PARP10 regulation dynamics^42^. Similarly, OPTN-mediated autophagy helps restrain pyroptosis by inhibiting inflammasome activation^43^. Together, these findings suggest a coordinated regulatory network wherein these molecules balance immune defense and inflammatory β-cell death.

LGALS3BP is another IIRS protein that is of particular interest. LGALS3BP, known as galectin-3 binding protein (Gal-3BP or Mac-2-binding protein/90K), is a ubiquitously expressed secreted glycoprotein with diverse physiological roles. It normally helps modulate cell adhesion and immune surveillance, but its levels increase during infections or immune challenges^44^. A recent targeted proteomics study on TEDDY plasma samples found that LGALS3BP levels rise about 6 months before seroconversion, followed by fluctuations thereafter^45^, suggesting it might serve as an early biomarker of immune activation in T1D. In our study, we observed markedly elevated levels and a strong IIRS association of LGALS3BP in islets of stage 1 T1D compared to ND donors. This observation aligns with a prior report showing increased LGALS3BP levels across multiple islet cell types in AAB^+^ patients^9^. Intriguingly, higher expression of LGALS3, the binding partner of LGALS3BP, was also observed in islets exhibiting either diminished BCP or elevated IIRS (**Extended Data Fig. 8**). LGALS3 has been shown to promote β-cell dysfunction and apoptosis, and immune cell recruitment ^46^. Thus, the elevation of LGALS3 in islets with lower BCP or higher IIRS indicates its dual role in driving both immune activation and β-cell dysfunction. While the precise interplay of these gelatin proteins in islet biology requires further investigation, they represent promising candidates for biomarker development and potential immunotherapeutic strategies in T1D.

During stage 1 diabetes, a progressive loss of β-cell function has been observed in longitudinal studies^5,6^. Here we report changes to the islet proteome that suggest pathways that contribute to this loss of β-cell function or identity in a larger proportion of islets in stage 1 T1D cases compared to ND controls. The variations in the expression of BCP (**Fig. 5**) even within ND controls suggest the existence of islet heterogeneity in ND individuals. However, the observation of a higher proportion of islets with low expression of BCP in stage 1 donors compared to ND controls does present evidence supporting a loss of β-cell function or identity in these donors. Our data are also consistent with prior reports about β-cell dedifferentiation, where selective loss of β-cell identity markers (e.g., INS, IAPP, PTPRN) occurs before extensive β-cell loss^13^. The most notable pathways associated with BCP are mRNA processing and RNA splicing. Altered mRNA processing was reported as strongly linked with the loss of β-cell function^38^. mRNA processing and alternative splicing in pancreatic β cells have also been suggested as a new frontier in the pathogenesis of T1D^47^. Our in-situ proteomic data further support the involvement of mRNA processing and alternative splicing in β-cell dysfunction and loss, offering new insights into the molecular mechanisms underlying T1D pathogenesis.

ECM-related pathways are another category consistently identified in correlation analyses with both IIRS and BCP. Islet ECM serves as both a barrier and a conduit for immune cell migration. During T1D, the ECM undergoes continuous cycles of degradation and rebuilding, driven by both infiltrating immune cells and β-cells responding to inflammatoin^48^. Breakdown of specific ECM components, including type IV collagen (COL IV) via MMP-3, laminin, hyaluronan, and heparan sulfate, has been reported in T1D-prone NOD mice and in human donors with mAAb⁺ or long-standing T1D^49^. Notably, ECM-related pathways displayed a stronger correlation with BCP in ND controls than stage 1 T1D cases, both in terms of fold changes and statistical significance, suggesting that ECM regulation is protective in healthy islets but becomes dysregulated in stage 1 T1D. We identified significantly lower log₂ fold changes of COL IVA2, a core islet basement membrane component, in stage 1 T1D compared to ND donors. This finding corroborates an observation by Johansen et al.^17^, who reported decreased COL IV levels in the same set of stage 1 mAAb⁺ donors. Building on this, we also detected disruptions in additional collagen isoforms alongside remodeling of actin cytoskeleton components and hyaluronan-associated proteins.

In contrast, ECM-related pathways were negatively correlated with BCP, particularly the higher expression levels of these proteins observed in islets with reduced β-cell function, suggesting that ECM remodeling contributes to β-cell dysfunction or loss (**Extended Data Fig. 7B-D**). Among them, we observed high FAP and DPP4 expressions in BCP low islets. Both proteins have been previously known to be present in human islets^50^, with potential implications of local peptide processing. FAP is known to cleave and inactivate human FGF21, a hormone that supports β-cell survival or function^51^ and is a known marker for activated fibroblast/stellate cells that remodel peri-islet ECM^52^. Whereas DPP4 is known to protect immortalized β-cells (EndoC-βH1) from cytokine toxicity (partly GLP-1-independent), improve GSIS, and reduce apoptosis^53^. We also identified S100A10, SH3PXD2B (TKS4), LPCAT2, and MMRN2 as some of the understudied ECM remodeling/organization proteins in the context of β-cell dysfunction. Together, these findings support the notion that ECM plays a bifunctional role, providing protective effects against insulitis and inflammation under normal conditions, while certain conditions exacerbate β-cell dysfunction.

There are several limitations of the present study to be acknowledged. First, due to the limited throughput of the current proteomics workflow, our study analyzed only ∼100 islets per donor and a total of ∼450 islets from three stage 1 T1D cases with three matched ND controls. Second, pancreas tissue inherently represents a single “snapshot” of disease, and thus, the pseudo-temporal progression reconstructed here does not fully reflect the natural history of T1D development. Moreover, the apparent “independent” occurrence of β-cell dysfunction and islet inflammation observed in this and previous study^18^ likely reflects the cross sectional or ‘snapshot’ nature of the samples. Islet inflammation (IIRS) probably represents a transient state of activated immune response, whereas β-cell dysfunction reflects the chronic, cumulative impact of signals within the islet microenvironment. Despite inherent limitations, our intra-donor, single-islet analyses substantially enhanced statistical power and uncovered reproducible proteomic patterns reflecting the progressive immune response and β-cell dysfunction in T1D. This high-resolution approach enabled the identification of in situ molecular pathways and novel protein candidates that warrant further functional investigation. Together, these findings demonstrate the potential of single-islet spatial proteomics to resolve intra-donor heterogeneity, providing a deeper understanding of how local immune and cellular processes drive islet pathology in early stages of T1D.

## MATERIALS AND METHODS

### Demographic information of organ donors

Organ donors from the Network for Pancreatic Organ Donors with Diabetes (nPOD) were selected for those who tested positive for multiple autoantibodies (mAAb+: against Insulin, GAD65, ZnT8, and IA-2A) and age/sex/ethnicity-matched ND controls (**Table 1**). Ethical permission was obtained from the Institutional Review Board at the University of Florida (NH00048338). All donors were between the ages of 15-25 and were otherwise healthy with no co-morbidities. The Body Mass Index (BMI) range for these donors was 19-28, with a mean and standard deviation of 23.2 ± 2.9.

### Multiplex immunohistochemistry and Laser Microdissection of Individual Islet Sections

Pancreatic tissue sections from fresh frozen Optimal Cutting Temperature (OCT)-embedded blocks were obtained from selected donors. From each tissue block, one section was used for mIHC, and adjacent serial sections were subjected to LMD. mIHC of insulin (INS), Glucagon (GCG), and CD3 was performed using methods previously described^54^. Sections were air-dried, followed by rinsing in phosphate-buffered saline (PBS) to remove OCT. Sections were fixed in 10% formalin for 10 minutes, followed by washing in PBS. Blocking was carried out for 1 hour at room temperature in 10% normal serum. Whole slide images were acquired using a Zeiss 710 confocal microscope with tile scanning. Individual islet images were acquired using a Zeiss 710 confocal microscope and 20x objective (Plan-Apochromat, 0.8 NA, 0.12µm/pixel, 8-bit depth with 16 frame averaging).

For LMD, frozen sections (10 µm) were placed on PEN slides using previously described methods ^55^. Sections were dehydrated in 100% methanol for 1 min, rinsed with diethylpyrocarbonate (DEPC)-treated water, followed by incubation for 1 min in 75% ethanol, 1 min in 95% ethanol, and two 1 min washes in 100% ethanol. All solutions were prepared fresh on the day of use and stored on ice. Following the last 100% ethanol incubation, slides were blotted, placed in an RNAase-free desiccator until dry (>8 mins) at room temperature under vacuum, and immediately subjected to LMD using a Leica LMD7000 microscope. Islets were identified by inherent autofluorescence and morphology within exocrine regions. Dissections were performed using a 10X objective using the following settings: speed 5, balance 15, head correct 100, less 4755, and offset 50. Approximately 100 individual islet sections were obtained from each mAAb+ case, and ∼50 individual islet sections were collected for each ND case. Each islet section was collected by gravity into the cap of AdhesiveCap 200 opaque (D) (Zeiss). Additional tissue sections, if available, were used to collect pooled samples of islet sections (∼100 sections per pool). All samples were immediately frozen on dry ice, then stored at -80°C until shipment on dry ice to PNNL.

### Islet transfer onto chips

Clean polypropylene microPOTS chips^56^ with 2.2 mm wells were preloaded with HPLC-grade dimethyl sulfoxide (DMSO). A total of ten Islet sections from different donors were pooled into each well. The 2.2 mm microPOTS chip was secured and stored at -80 °C until ready for further sample processing. For profiling of a single islet section, a series of nanoPOTS chips (1.4 mm diameter wells) were preloaded with 400 nL of HPLC-grade DMSO. The pre-sorted islet section samples were removed from -80 °C and placed on ice. The cap of AdhesiveCap 200 opaque (D) was removed from ice and brought inside an AirClean® Systems dead air box. Using a Dino-Lite USB camera, the patient islets observed on the Adhesive cap were transferred to each 1.4 mm nanowell with Excelta 4-CO tweezers. Once all tissues had been sorted, samples were stored at −80 °C until ready for sample processing.

### NanoPOTS and microPOTS sample processing

Before sample preparation, the nanoPOTS and microPOTS chips were removed from −80 °C and left at room temperature for 15 min. The chips were placed uncovered in an incubator set to 70 °C until all DMSO had evaporated. Using our in-house nanodroplet dispensing robotic system^24^, 200 nL of cell lysis buffer containing 0.1% (w/v) n-dodecyl-ß D-maltoside (DDM) and 2 mM DTT in 50 mM ammonium bicarbonate (ABC) buffer was dispensed into each 1.4 mm nanowell. For micropots, 1.6 µL of the cell lysis buffer was added to the 2.2 mm microwells. The nanoPOTS and microPOTS chips were sealed with a sterile microslide cover, placed in a humidity chamber, and incubated at 70 °C for 60 min. Then, the chips were removed from the incubator and left at room temperature for 15 min. Next, 50 nL of alkylation reagent containing 20 mM iodoacetamide (IAA) in 50 mM ABC was dispensed into each nanowell, and 200 nL of 10 mM IAA in 50 mM ABC buffer was dispensed into each 2.2 mm microwell. The samples were sealed and placed into a humidity chamber for 30 min at room temperature in the dark. Next, 50 nL of an enzyme solution containing 0.04 ng Trypsin and 0.02 ng Lys-C in 50 mM ABC was dispensed into each nanowell, and 200 nL were added to each 2.2 mm microwell before the chips were sealed, placed into a humidity chamber, and incubated for 10 hours at 37 °C. The samples were quenched with 50 nL formic acid (5%, v/v) into the wells for 15 min at room temperature. Finally, the droplets were left uncovered in a vacuum desiccator until all droplets had evaporated. All chips were secured and stored in −20 °C in a freezer before LC-MS analysis.

### LC-MS/MS Analysis

LC-MS/MS analyses were performed using our in-house automated nanoPOTS LC system^57^ coupled with a ThermoFisher Orbitrap Fusion Lumos Tribrid mass spectrometer (MS) with a high field asymmetric waveform ion mobility spectrometry (FAIMS) interface. The Thermo Ultimate 3000 LC system was configured for on-line trapping with reverse elution onto the analytical column. The nanoPOTS-collected samples were loaded at 3.5 μL/min onto an in-house packed Solid Phase Extraction (SPE) trapping column (100 μm i.d., 4.25 cm length, Jupiter C18, 300 Å pore size, 5 μm particles; Phenomenex, Torrance, CA, USA). Peptides were separated at 100 nL/min using a 25 cm long × 50 µm i.d. analytical column with integrated emitter (PicoFrit column, New Objective, Woburn, USA) packed in-house using Waters BEH 1.7 μm particles (Milford, MA). The analytical column was heated with a 15 cm Agile Sleeve capillary heater at 50 °C (Sales and Services, Inc., Flanders, NJ, USA). Reverse mobile phases consisted of (A) 0.1% formic acid in water and (B) 0.1% formic acid in acetonitrile with the following gradient profile for elution of peptides (min., %B): 0, 2; (sample loaded at 3.5 µl/min for 5 minutes); 1, 8; 45, 22; 60, 35; 62, 80; 63, 2; 66, 35; 68, 2; 71, 35; 73, 2; 75, 80; 83, 80; 84, 2; and 90, 2.

Samples were analyzed using data-dependent acquisition (DDA) mode using an Orbitrap Fusion Lumos Tribrid MS (Thermo Scientific) coupled with a FAIMS interface. To enhance the proteomic depth from a single islet, transferring identification based on the FAIMS filtering (TIFF) method was employed^58,59^. Peptides were ionized using a voltage of 2.4 kV, an ion transfer tube temperature at 200°C, and with 3.5 L/min carrier gas flow. Data acquisition time was 90 min following a 10 min delay to avoid a delay between injection and elution of peptides.

The microPOTS samples of pooled islet sections were acquired to generate a spectral library, each injection with a discrete compensation voltage (CV) of 45, –60, or –75 V with a cycle time of 2.5 s. The precursor ions with masses within 350-1500 m/z were scanned by Orbitrap at 120,000 FWHM resolution with an IT of 118 ms and an AGC target of 1E6. The selected precursors with charges from 2+ to +6 were fragmented at 30% HCD and scanned in an ion trap with an AGC of 2E4 and an IT of 100 ms.

For single islets analysis, the ionized peptides were fractionated by the FAIMS interface using three FAIMS compensation voltages (CV, –45, –60, and –75 V) for each LC-MS/MS analysis with cycle times of 0.8 s. The precursor ions with masses within 350–1500 m/z were scanned by Orbitrap at a 120,000 resolution with an injection time (IT) of 246 ms and an automatic gain control (AGC) target of 1E6. Selected precursor ions with +2 to +6 charges and intensities above 1E4 were selected for fragmentation by 30% high collision dissociation (HCD) and scanned in an ion trap with an AGC of 2E4 and an IT of 100 ms. Cycle times of 0.8 s were used for the combined 3-CV method.

### Data Processing

#### Preprocessing

All LC-MS/MS raw files were processed using the FragPipe (v21.1) computational platform with MSFragger (v4.0)^60,61^, and Philosopher (v5.1.0)^62^ for FDR (false discovery rate) filtering and reporting, and IonQuant (v1.10.12)^63^ for label-free quantification (LFQ) with matching between runs (MBR). All MS/MS spectra were searched against a human UniProt (Homo sapiens) database (Oct 9, 2024 downloaded) containing 20,476 protein sequences, 115 common contaminant sequences, and decoy protein sequences.

For LFQ-MBR analysis, the following search parameters were used for MS/MS search: full tryptic specificity up to three missed cleavage sites, carbamidomethylation (57.0214 Da) on cysteine as a fixed modification, and methionine oxidation (+15.9949) and protein N-terminal acetylation (42.0105 Da) as variable modifications. The initial precursor and fragment mass tolerances were set to ±20 ppm. After mass calibration, MSFragger adjusted the tolerances automatically. The identification results were filtered with sequential 1% PSM-, peptide-, and protein-level FDR. The MBR algorithm in IonQuant was activated with a matching RT tolerance of 0.4 min, m/z tolerance of 10 ppm, and an ion-level FDR threshold of 5%. MBR candidate ions were selected from pooled library samples from their respective mass spectrometry run batches. Normalization was disabled, and the MaxLFQ algorithm was used to quantify ion-level intensities.

Peptides were rolled to protein level with <1% FDR, log2-transformed, and protein features zero-centered, before sample normalization with medpolish^64^ based on complete cases, for each donor. Features with unusually high missingness were removed. Batch correction for pancreas tissue block location was applied to datasets within each donor using ComBat^65^. Samples across donors were combined (n = 439), and features were filtered to those with >50% completeness within each donor (n = 4519 protein groups).

#### Clustering analysis

WGCNA^66^ clustering analysis was initially applied on single islet datasets from each donor independently to identify potential modules with significant association with a given phenotype (e.g., CD3 or insulin intensity from MS data). To test these associations, the limma (v3.58.1) framework was employed: moderated t-tests were applied to assess differential expression of module eigengenes (MEs; principal eigenvectors of each module) across the dichotomous CD3 phenotype, while moderated linear models were fit to evaluate correlations of MEs with insulin intensity. ORA based on Gene Ontology Biological Process (GO:BP) terms was applied to each module to identify significantly enriched BP terms. ORA was performed with the enrichGO function from the clusterProfiler R package^67^ to test GO:BP gene sets^68,69^.

Clustering analyses were also applied to identify protein modules by combining data across all donors. In this case, samples were first separated by disease status (stage 1 T1D vs. ND) to handle missing values in clustering analyses. Within each class, missing data were imputed using single value decomposition (SVD) via the svdImpute function from the pcaMethods package (v1.94.0). The imputation was performed on z-scored features, and the original feature scales were restored afterwards. To improve cluster interpretability, clustering settings were optimized with the following settings implemented. Clustering was performed by calculating the topological overlap matrix (TOM) of type signed Nowick 2 as implemented in the WGCNA package (v1.73). Feature correlation was evaluated using the biweight midcorrelation, and soft power was set to 12. The TOM was converted to a distance and subjected to hierarchical clustering using Ward’s D linkage. The resulting dendrogram was cut to produce 75 final clusters.

#### Selection of protein markers for the Islet Immune Response Signature (IIRS)

Following the observation of protein modules significantly enriched with innate and adaptive immune response from all three stage 1 T1D (mAAb+) donors, a machine-learning-based feature selection approach was applied to identify members for the Islet Immune Response Signature (IIRS). Proteins from all modules enriched with innate and adaptive immune response were combined and used as input for feature selection. Protein intensity data of the six donors were combined and batch corrected for donor effects using ComBat^65^ before the feature selection process.

The R mlr3 machine learning ecosystem (mlr3verse v0.3.1)^70^ was used to tune, train, and evaluate the machine learning model. A random forest classification model was chosen because tree-based models inherently provide knowledge of feature importance and are robust to multicollinearity. Protein abundances were used in modeling after filtering proteins for those included in WGCNA immune modules and requiring features to contain fewer than 50% missing values within each donor (n = 238). A random forest learner from the ranger package was employed, leveraging permutation-based feature importance scores. The learner was configured for probabilistic prediction and stratified resampling based on prediction class to ensure the same proportion of class labels. Hyperparameters were tuned using a random search with 10 evaluations. Tuned hyperparameters include the number of features considered as candidates for splitting at each node (mtry) and the minimum number of samples required to form a terminal node (min.node.size). The number of trees grown in each ensemble was fixed at 1000. Nested cross-validation was employed with tuning and feature selection embedded in the inner loop (5-fold CV) to segregate parameter selection from model evaluation. Features were selected using recursive feature elimination, iteratively removing features with the lowest 10% importance scores, with a target of 60 features. Hyperparameter tuning was performed for each iteration of feature selection, and model performance was assessed using area under the ROC curve. The top-performing features from the inner folds were evaluated on the hold-out set from the outer loop (10-fold CV). This feature selection procedure was repeated for each of the 10 outer folds, providing 10 independent feature sets. Each feature set is ordered by the feature importance score derived from the final model. These sets were integrated using robust rank aggregation (RRA)^71^, as implemented in the RobustRankAggreg package, ranking proteins based on the consistency of their relative importance across all sets. The Islet Immune Response Signature (IIRS) was selected by taking the 40 proteins with the lowest RRA scores (p-values). These IIRS markers were aggregated to provide a single representative feature by taking sample-wise median.

#### Differential abundance analysis

Linear Models for Microarray Data (LIMMA) was used to fit a model with the centroid of the islet immune response signature (IIRS) as the sole predictor of the log_2_-transformed and median-centered protein abundance^72,73^. LIMMA borrows information about the entire ensemble of features being tested to augment the available degrees of freedom, thereby increasing statistical power. Moderated t-tests were performed to evaluate the strength of the linear relationships, and the moderated t-statistics were then transformed into their standard normal equivalents (z-scores) for use with pre-ranked Correlation Adjusted MEan RAnk (CAMERA-PR) gene set testing. CAMERA-PR is an extension of the two-sample t-test that accounts for correlation between genes^32^, as failure to account for such correlation has been shown to drastically inflate the type I error rate^32,74^. Gene sets from the Gene Ontology Biological Processes database^68,69^ provided by the org.Hs.eg.db R/Bioconductor package (v3.20.0)^75–77^, were tested only if they contained at least 5 genes that overlapped with the proteomics data. The CAMERA-PR t-statistics were converted to z-scores, and p-values were adjusted separately by donor using the method of Benjamini and Hochberg to control the false discovery rate (FDR)^78^. The TMSig R/Bioconductor package was used to perform CAMERA-PR with the default inter-gene correlation of 0.01 and generate a bubble heatmap to visualize the top gene sets per donor^75,76,79^. The bubble diameter is controlled by the –log_10_(adjusted p-value) and scaled relative to the most significant result by row (gene set) to better observe inter-donor differences. The color of each bubble indicates the strength and direction of the association between the immune signature centroid and the genes in the set relative to all other measured genes. Results with adjusted p-values below 0.05 are identified by a gray background.

#### Selection of markers for β-cell profile

Markers for β-cell profile were identified by association with insulin and ENTPD3. All features were evaluated for correlation with INS and ENTPD3, and candidate markers were ranked by their mean correlation. These markers were aggregated to form a single representative feature by taking the sample-wise median.

## Acknowledgements

We thank the donors and families of the donors for their invaluable contribution to our research and their help to further understand and hopefully cure type 1 diabetes. This research was performed with the support of the Network for Pancreatic Organ donors with Diabetes (nPOD; RRID:SCR_014641), a collaborative type 1 diabetes research project supported by Breakthrough T1D and The Leona M. & Harry B. Helmsley Charitable Trust (Grant#3-SRA-2023-1417-S-B). The content and views expressed are the responsibility of this article’s authors and do not necessarily reflect the official view of nPOD. Organ Procurement Organizations (OPO) partnering with nPOD to provide research resources are listed at https://npod.org/for-partners/npod-partners/. This research was supported by NIH Grants R01DK122160, R01DK135081, R01DK131059, R01DK123329, P01AI42288, and U01DK137113. Proteomics was performed in the Environmental Molecular Sciences Laboratory, a national scientific user facility sponsored by the DOE and located at Pacific Northwest National Laboratory, which is operated by Battelle Memorial Institute for the DOE under Contract DE-AC05-76RL0 1830. This work utilized a LEICA 7000 laser microdissection microscope purchased with a NIH shared instrumentation grant S10OD016350 and operated by the University of Florida Molecular Pathology Core (RRID:SCR_016601).

## Extended Figure Legends

**Extended data figure 1:**
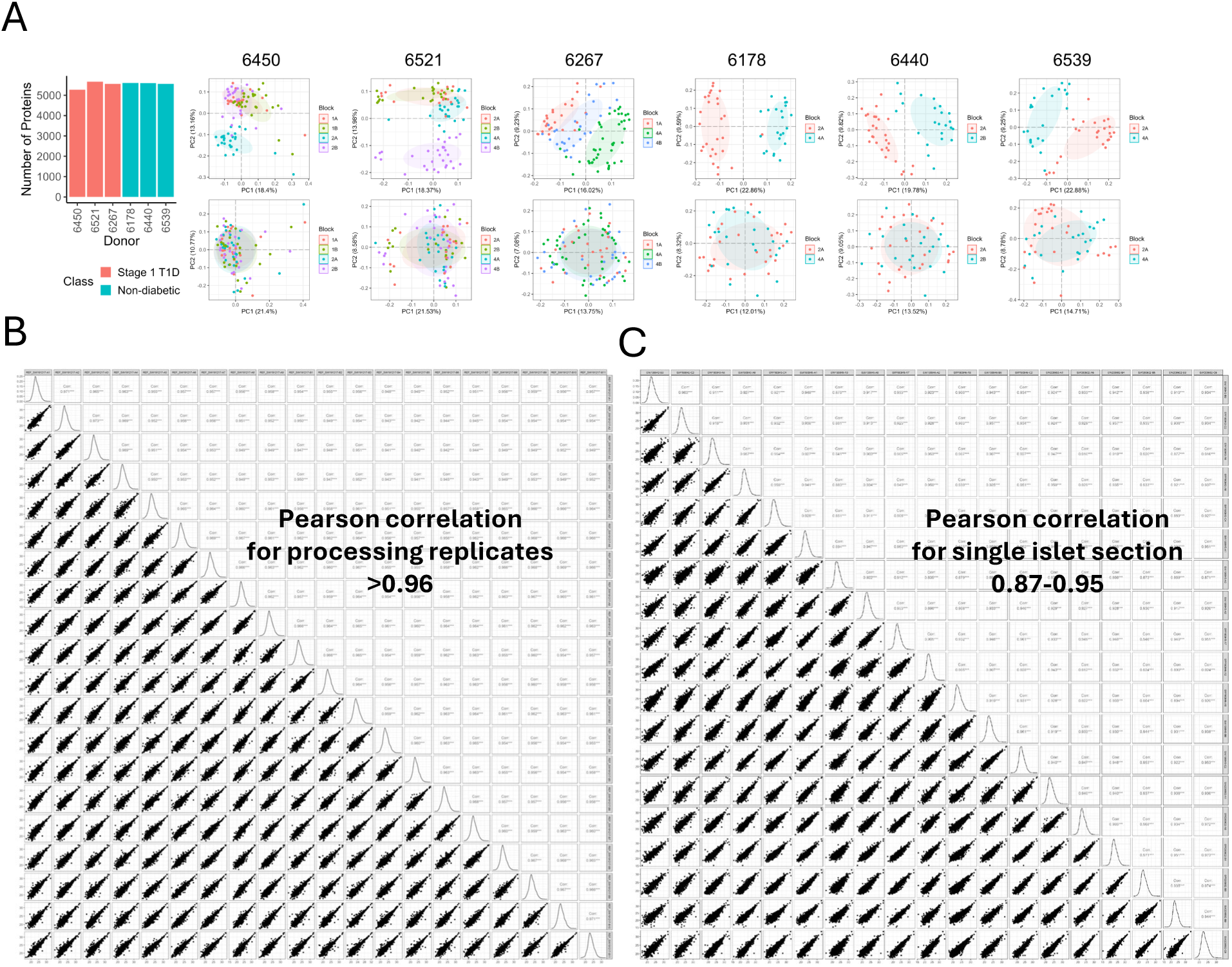
Assessment of overall data quality. **A.** Number of proteins quantified with greater than 50% completeness per donor along with PCA plots for each donor before and after batch correction for pancreas tissue block location. **B.** Sample-wise Pearson correlation among replicate reference samples inserted across samples for case 6450. **C.** Sample-wise Pearson correlation among randomly selected islet samples from case 6450.

**Extended data figure 2:**
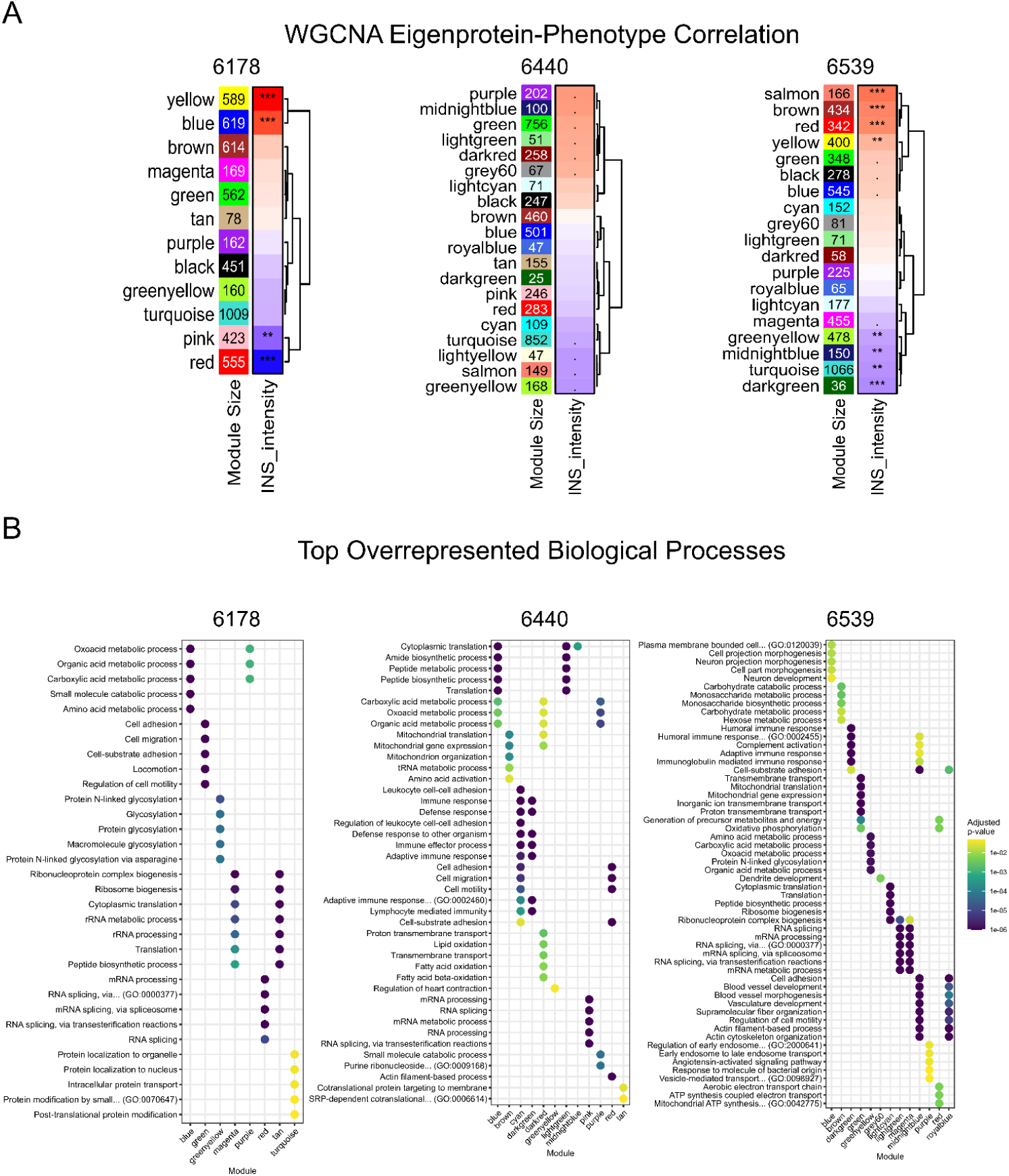
**A.** WGCNA identified protein modules correlated with insulin (INS) intensity for ND control donors with case ID 6178, 6440, & 6539. **B.** Top five overrepresented biological processes for each of the WGCNA protein modules with significant enrichment in at least one GO:BP term, with color scale representing the adjusted p-value.

**Extended data figure 3:**
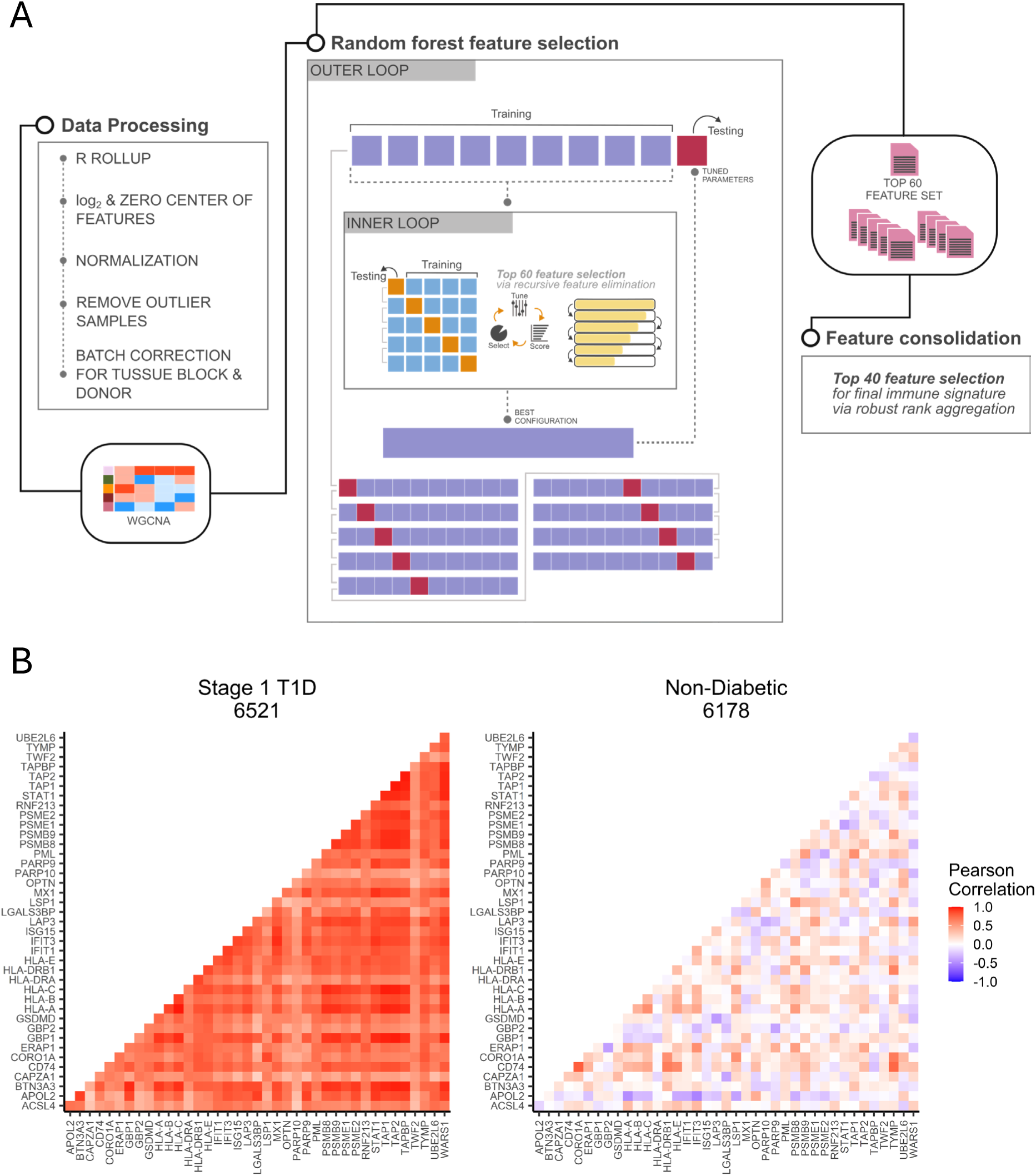
**A.** Workflow of machine learning feature selection pipeline including data processing, selection of candidate features by WGCNA, feature selection by random forest classification of single islet disease status with nested cross-validation for hyperparameter tuning and recursive feature elimination, culminating in consolidation of the outer fold feature sets by robust rank aggregation to produce the IIRS protein panel. **B.** Intra-donor correlation between pairs of IIRS proteins, highlighting coregulation of immune response proteins in stage 1 T1D (case 6521), which is absent in ND donors (case 6178).

**Extended data figure 4:**
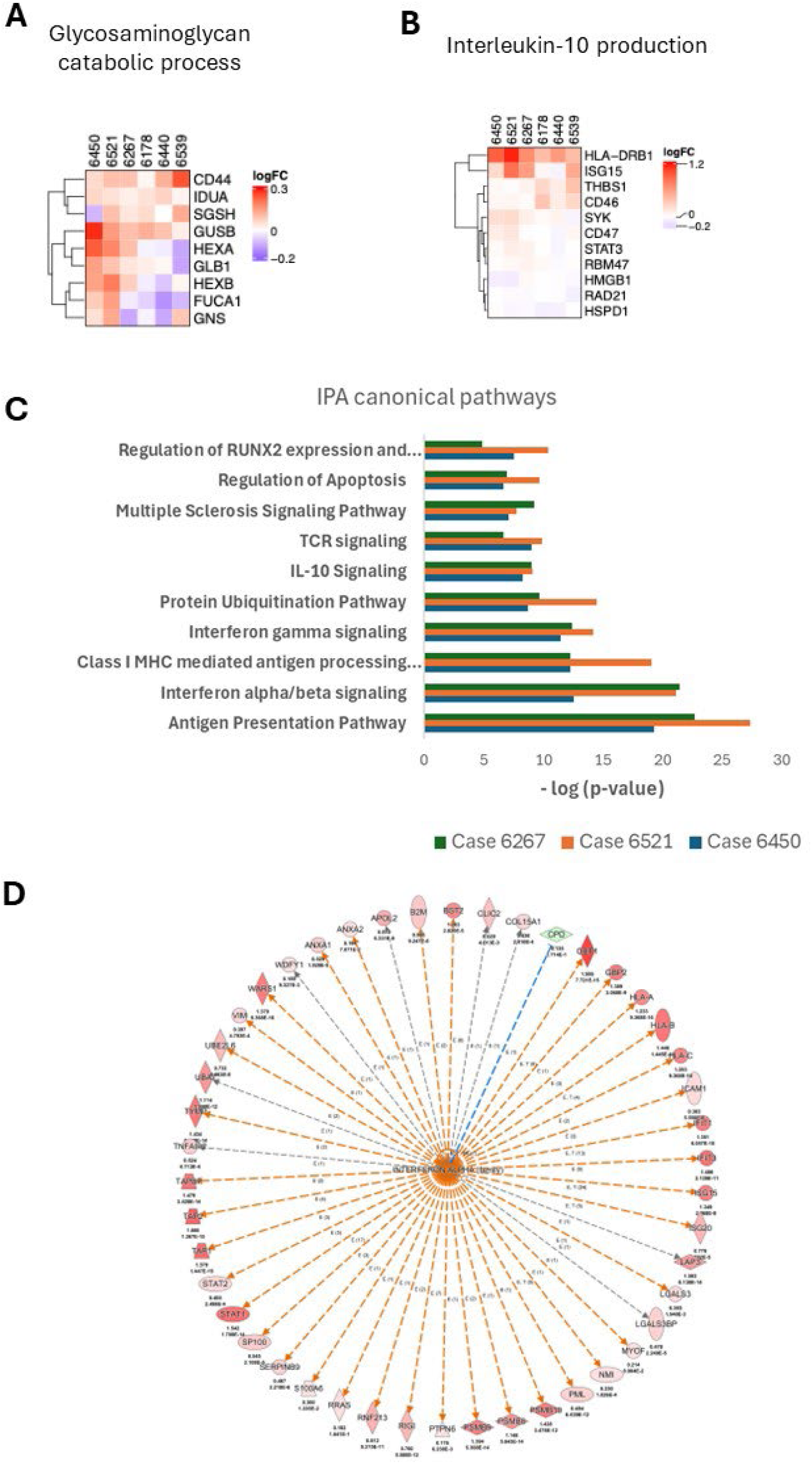
**A. & B.** Heatmaps for proteins in Glycosaminoglycan catabolic process and Interleukin-10 production, respectively, showing log2FC of each protein across individual donors. **C.** Top canonical pathways from Ingenuity Pathway Analysis (IPA) based on the significant resulst from LIMMA analysis against IIRS. **D.** Protein network of the interferon alpha family constructed from our dataset and identified as the top upstream regulator by IPA.

**Extended data figure 5:**
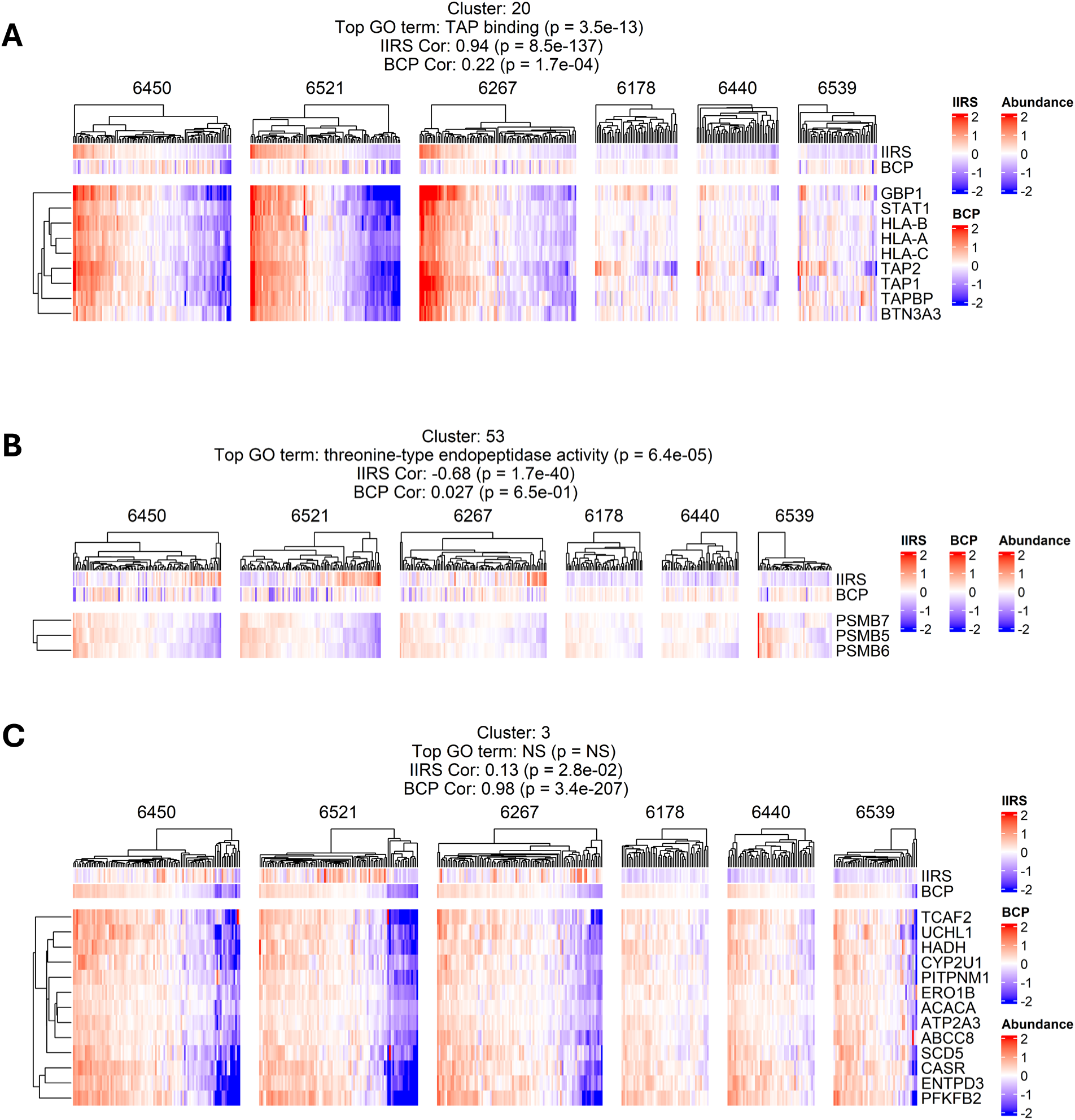
**A. & B.** Functionally distinct clusters with strong IIRS correlation, but weak association with BCP. Protein abundances are log2-transformed and median-centered across all samples. **C.** Protein cluster with strong BCP correlation, but weak association with IIRS.

**Extended data figure 6:**
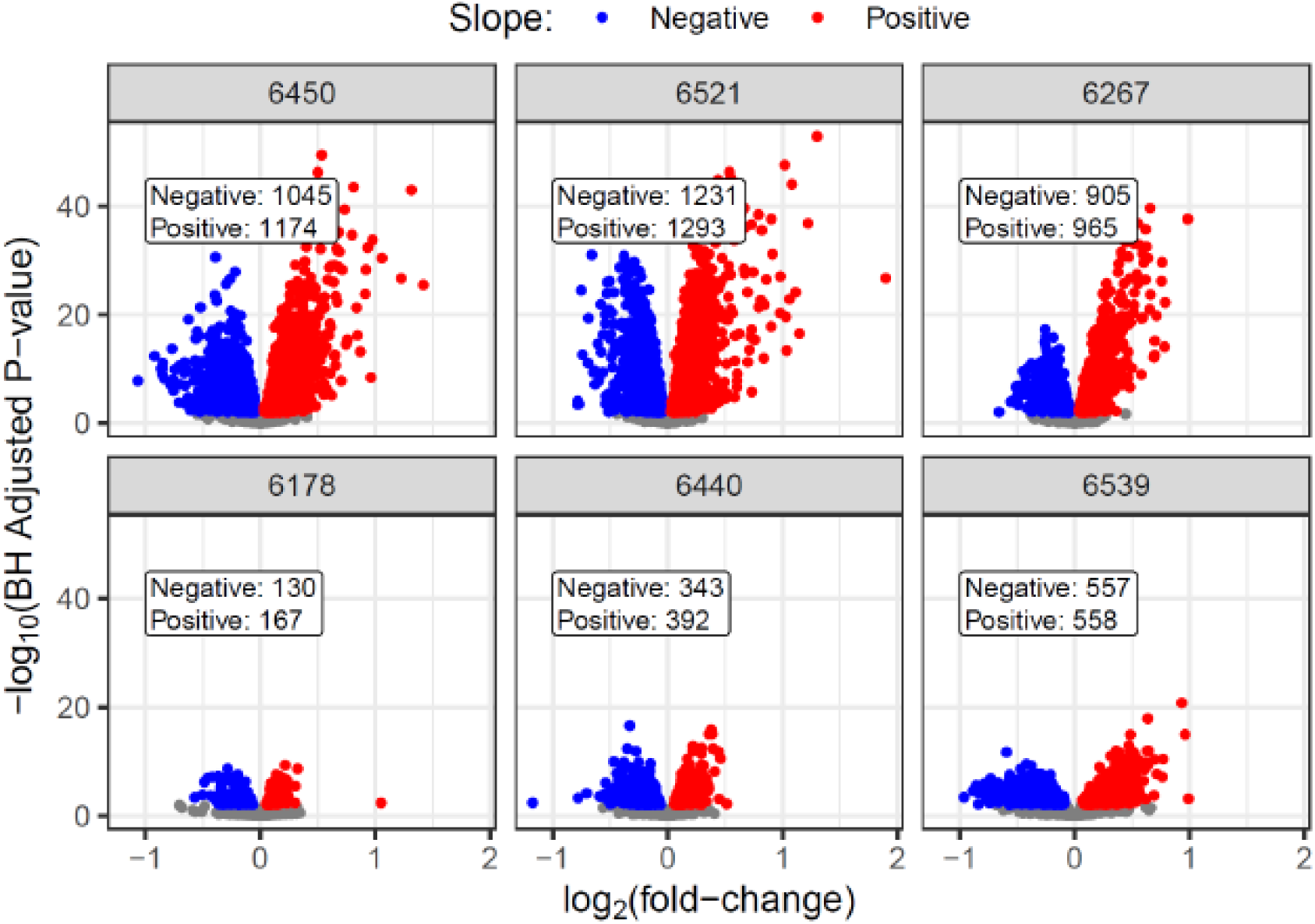
Volcano plots of the LIMMA analysis results against the BCP as the predictor of protein abundance for individual donors. The x-axis is the log2(fold-change) (i.e., slope of each fitted line), and the y-axis is the –log_10_(adjusted p-value). Red indicates a positive slope and blue indicates a negative slope in the protein-IIRS correlation. Colored points indicate adjusted p-value <0.01.

**Extended data figure 7:**
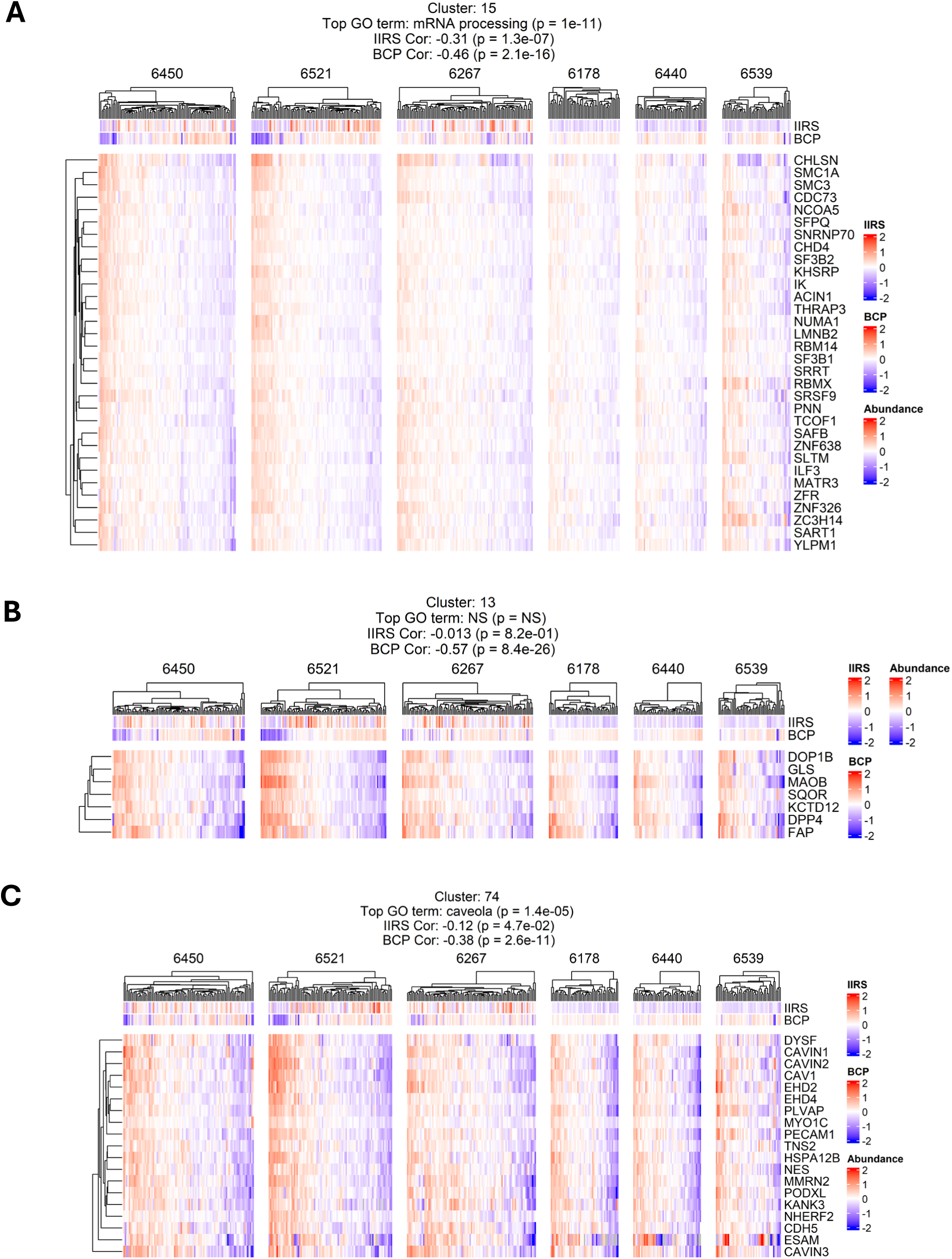
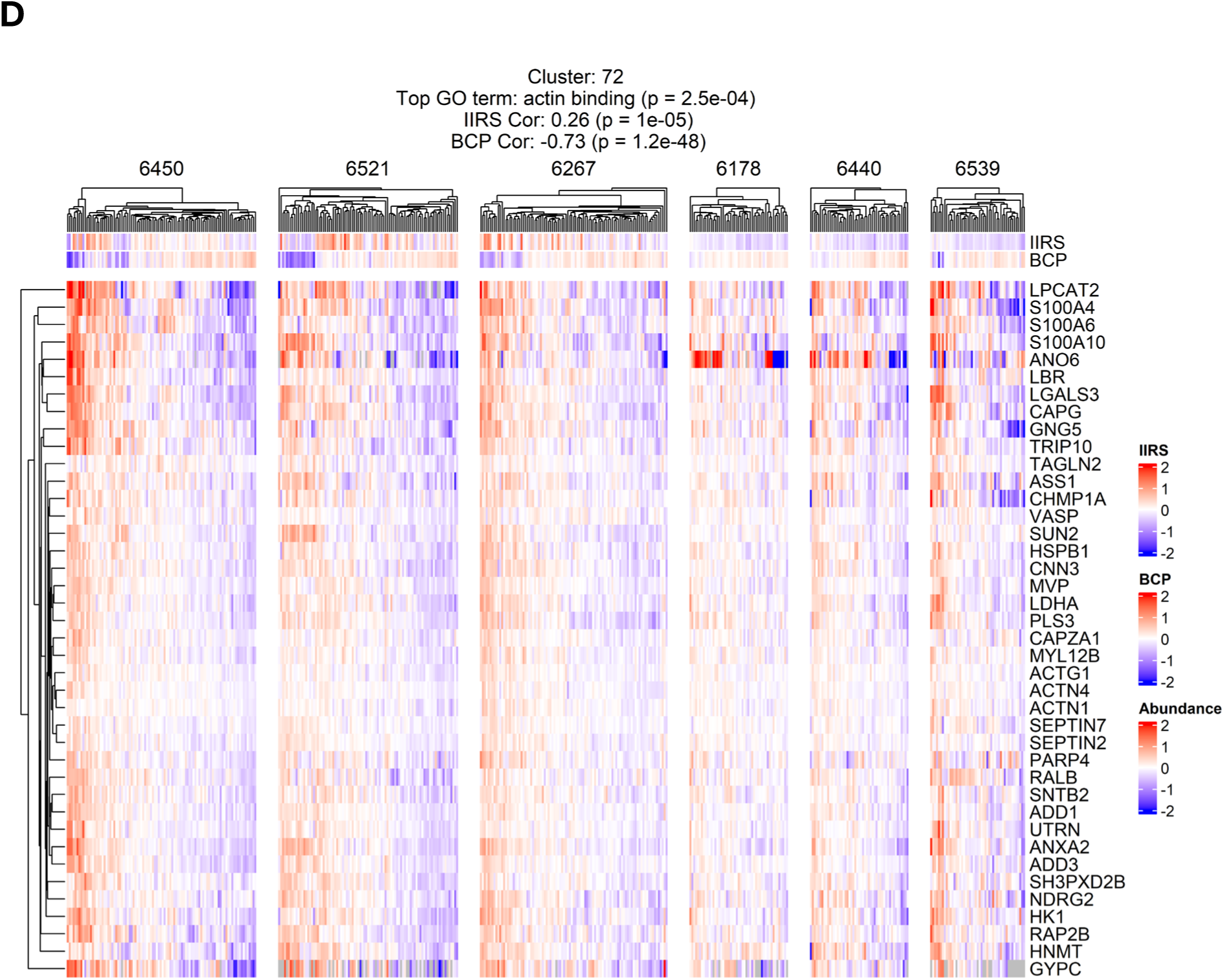
Clusters negatively correlated with BCP. **A.** Abundance of co-regulated proteins enriched in mRNA processing, which exhibits a negative association with BCP. Protein abundances are log2-transformed and median-centered across all samples. **B. C. & D.** ECM-associated clusters with negative BCP correlation.

**Extended data figure 8:**
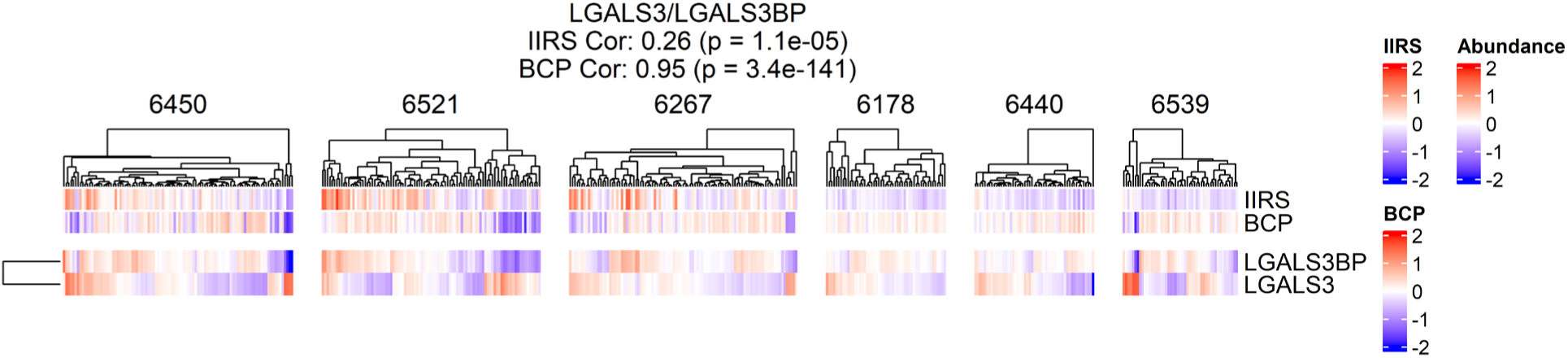
Abundance heatmap of LGAL3BP and LGALS3.

## Supplementary Information

The supplementary material includes 5 additional supplementary files.

