## Supplementary File 4 for "Single-Islet Proteomics Maps Pseudo-Temporal Islet Immune Responses and Dysfunction in Stage 1 Type 1 Diabetes"

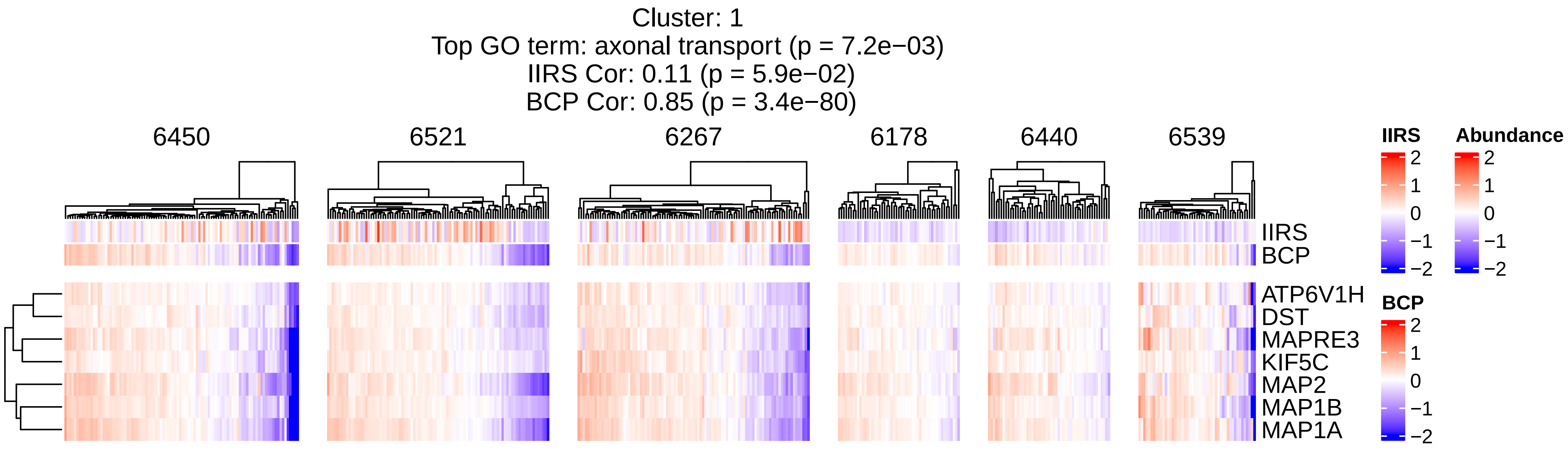

Cluster: 1
Top GO term: axonal transport (p = 7.2e−03)
IIRS Cor: 0.11 (p = 5.9e−02)
BCP Cor: 0.85 (p = 3.4e−80)
6450
6521
6267
6178
6440
6539
IIRS
BCP
ATP6V1H
DST
MAPRE3
KIF5C
MAP2
MAP1B
MAP1A
IIRS
2
1
0
−1
−2
Abundance
2
1
0
−1
−2
BCP
2
1
0
−1
−2

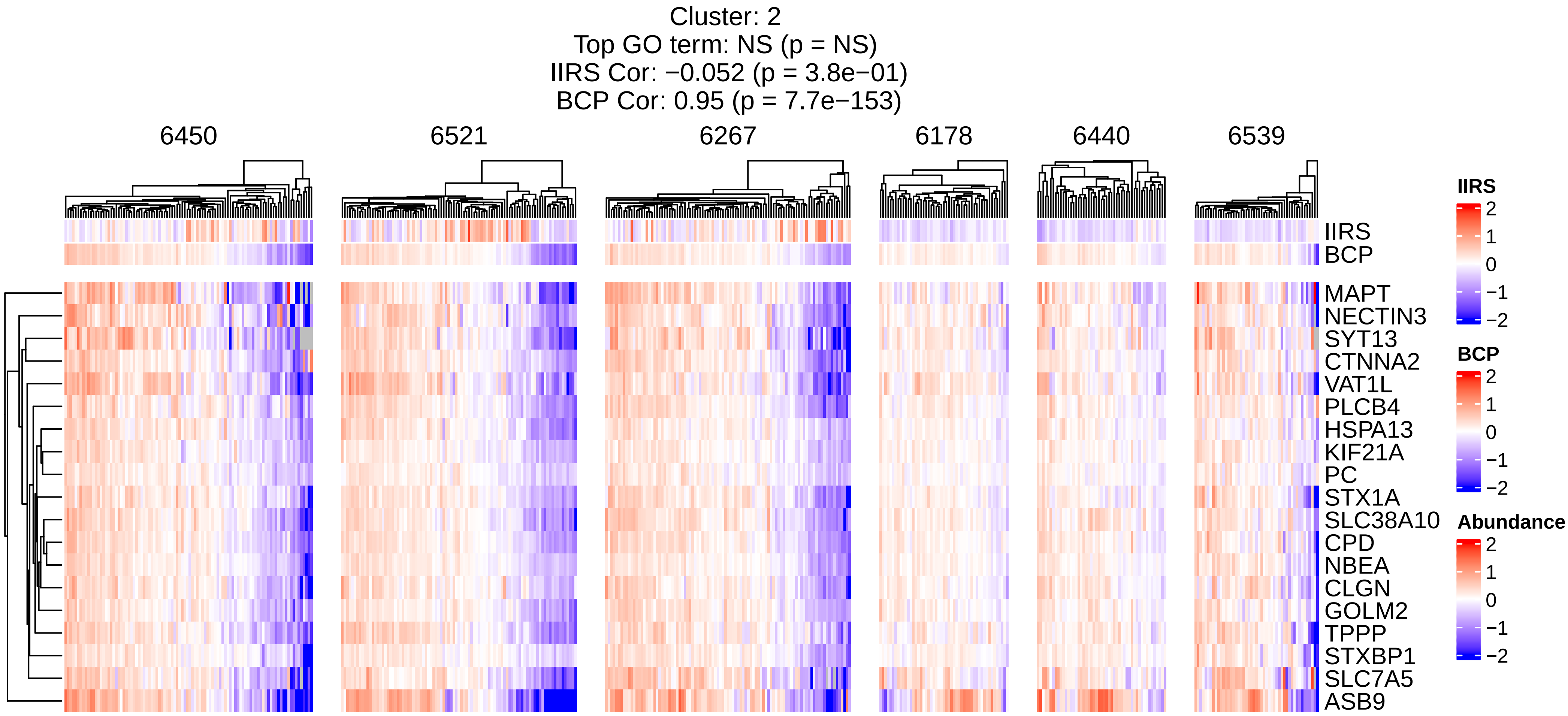
Cluster: 2
Top GO term: NS (p = NS)
IIRS Cor: −0.052 (p = 3.8e−01)
BCP Cor: 0.95 (p = 7.7e−153)
6450
6521
6267
6178
6440
6539
IIRS
BCP
MAPT
NECTIN3
SYT13
CTNNA2
VAT1L
PLCB4
HSPA13
KIF21A
PC
STX1A
SLC38A10
CPD
NBEA
CLGN
GOLM2
TPPP
STXBP1
SLC7A5
ASB9
IIRS
2
1
0
−1
−2
BCP
2
1
0
−1
−2
Abundance
2
1
0
−1
−2

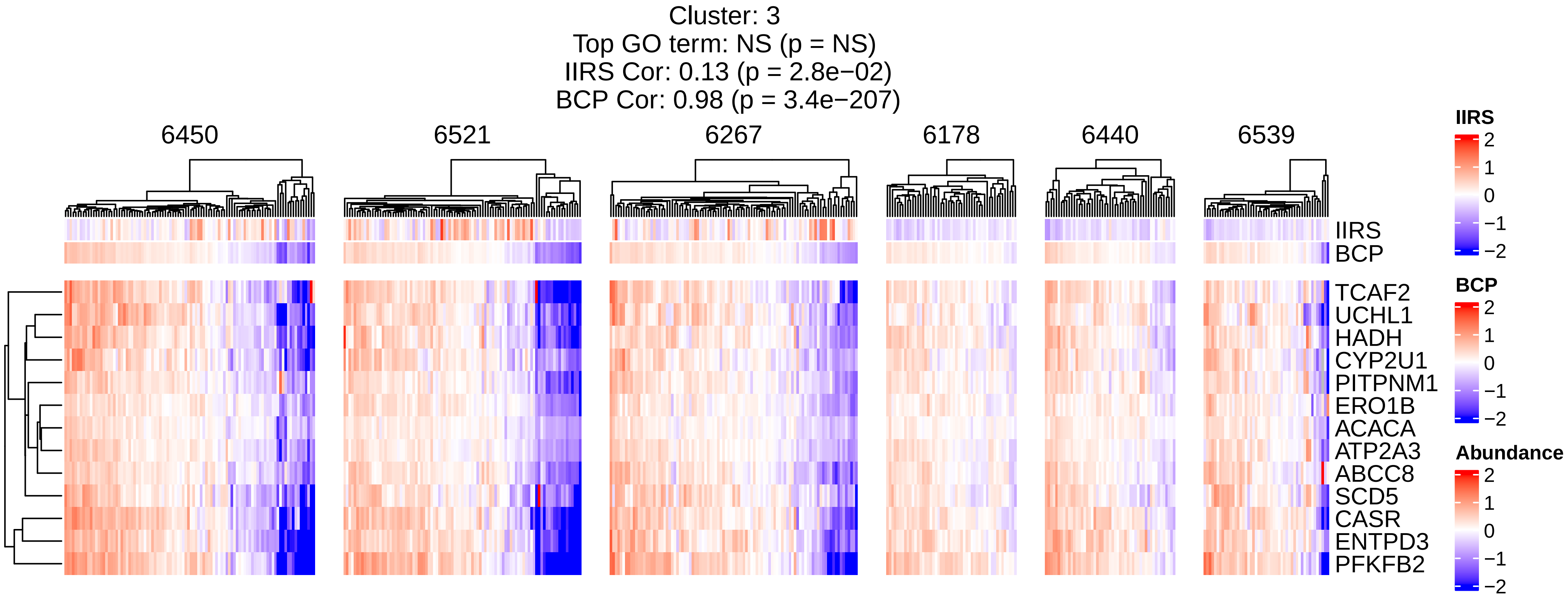

Cluster: 3
Top GO term: NS (p = NS)
IIRS Cor: 0.13 (p = 2.8e−02)
BCP Cor: 0.98 (p = 3.4e−207)
6450
6521
6267
6178
6440
6539
IIRS
BCP
TCAF2
UCHL1
HADH
CYP2U1
PITPNM1
ERO1B
ACACA
ATP2A3
ABCC8
SCD5
CASR
ENTPD3
PFKFB2
IIRS
2
1
0
−1
−2
BCP
2
1
0
−1
−2
Abundance
2
1
0
−1
−2

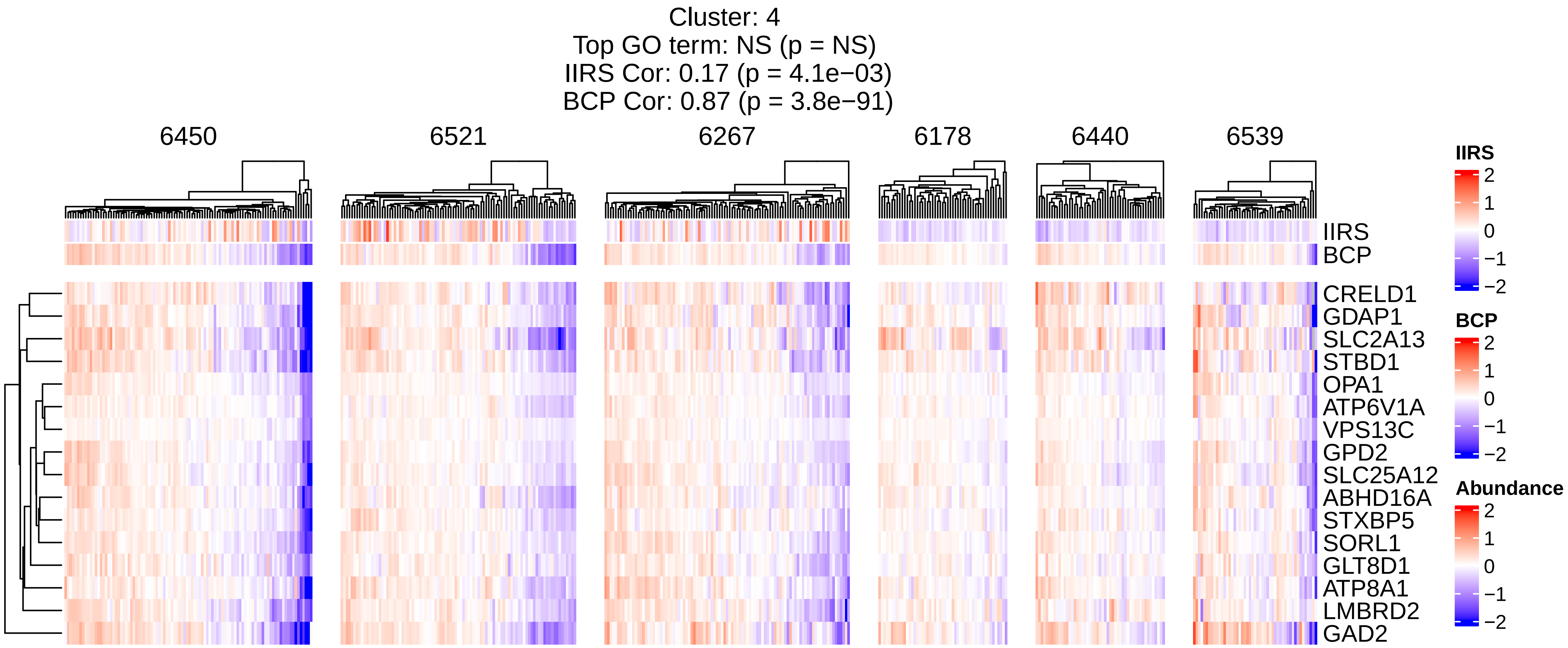
Cluster: 4
Top GO term: NS (p = NS)
IIRS Cor: 0.17 (p = 4.1e−03)
BCP Cor: 0.87 (p = 3.8e−91)
6450
6521
6267
6178
6440
6539
IIRS
BCP
CRELD1
GDAP1
SLC2A13
STBD1
OPA1
ATP6V1A
VPS13C
GPD2
SLC25A12
ABHD16A
STXBP5
SORL1
GLT8D1
ATP8A1
LMBRD2
GAD2
IIRS
2
1
0
−1
−2
BCP
2
1
0
−1
−2
Abundance
2
1
0
−1
−2

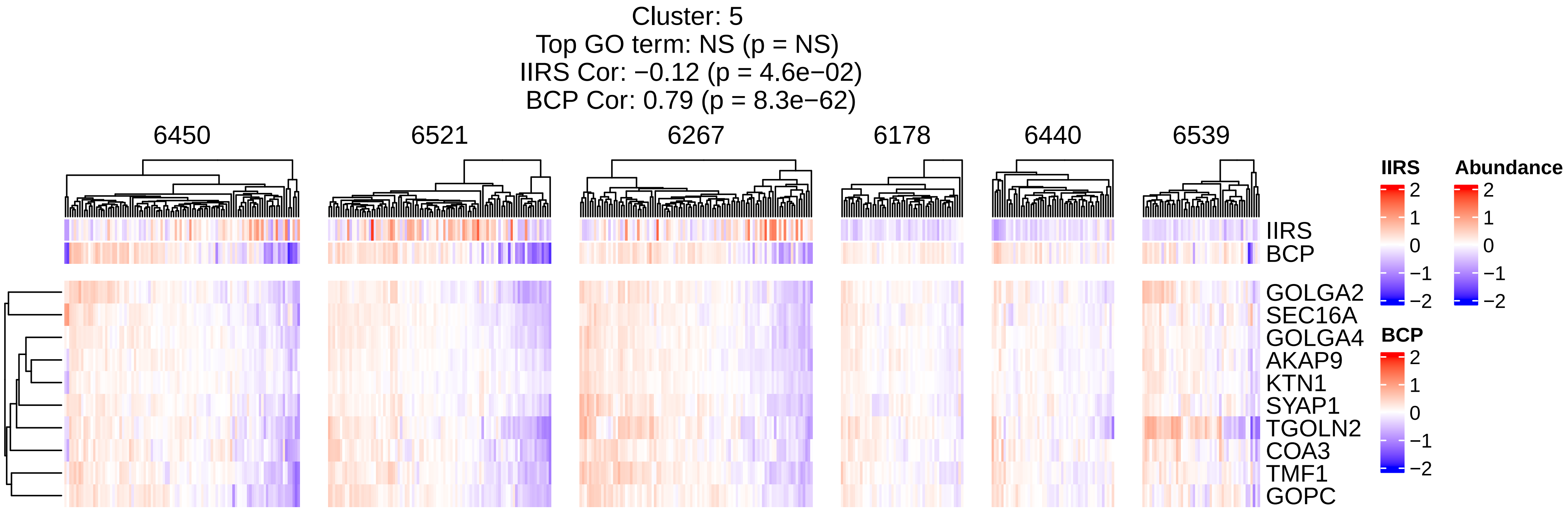
Cluster: 5
Top GO term: NS (p = NS)
IIRS Cor: −0.12 (p = 4.6e−02)
BCP Cor: 0.79 (p = 8.3e−62)
6450
6521
6267
6178
6440
6539
IIRS
BCP
GOLGA2
SEC16A
GOLGA4
AKAP9
KTN1
SYAP1
TGOLN2
COA3
TMF1
GOPC
IIRS
2
1
0
−1
−2
Abundance
2
1
0
−1
−2
BCP
2
1
0
−1
−2

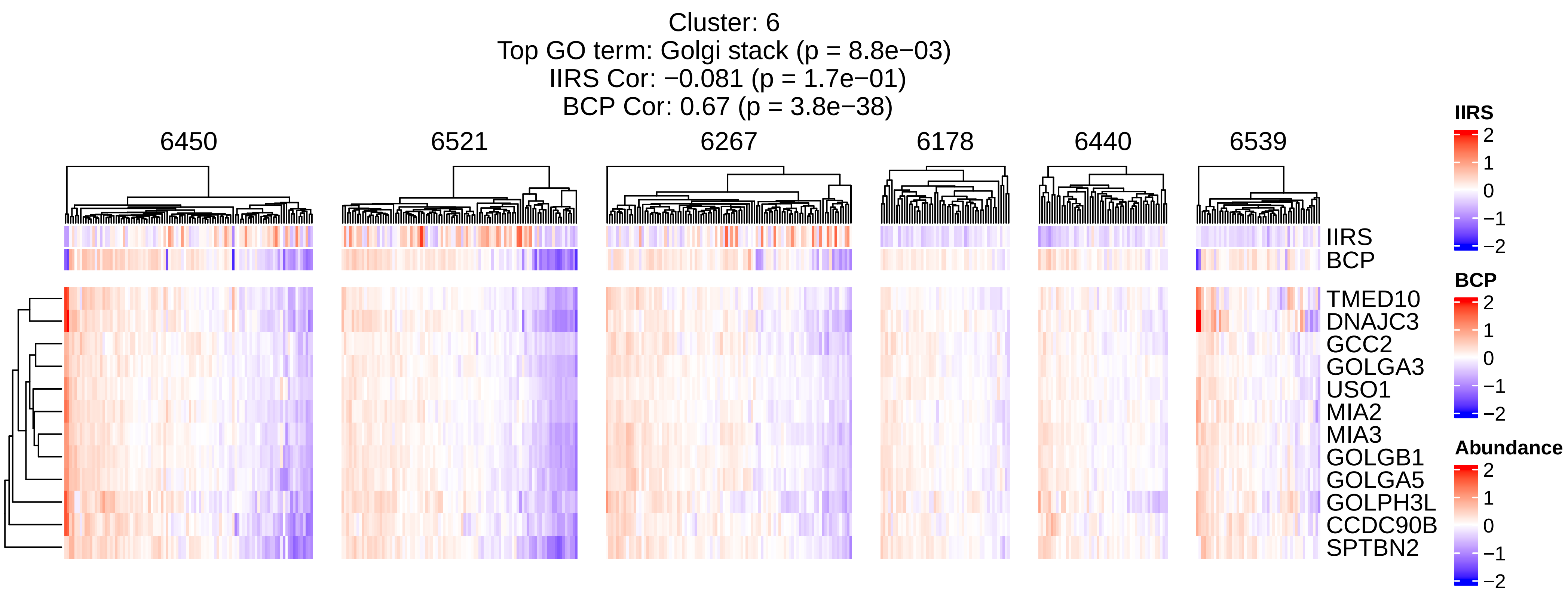
Cluster: 6
Top GO term: Golgi stack (p = 8.8e−03)
IIRS Cor: −0.081 (p = 1.7e−01)
BCP Cor: 0.67 (p = 3.8e−38)
6450
6521
6267
6178
6440
6539
IIRS
BCP
TMED10
DNAJC3
GCC2
GOLGA3
USO1
MIA2
MIA3
GOLGB1
GOLGA5
GOLPH3L
CCDC90B
SPTBN2
IIRS
2
1
0
−1
−2
BCP
2
1
0
−1
−2
Abundance
2
1
0
−1
−2

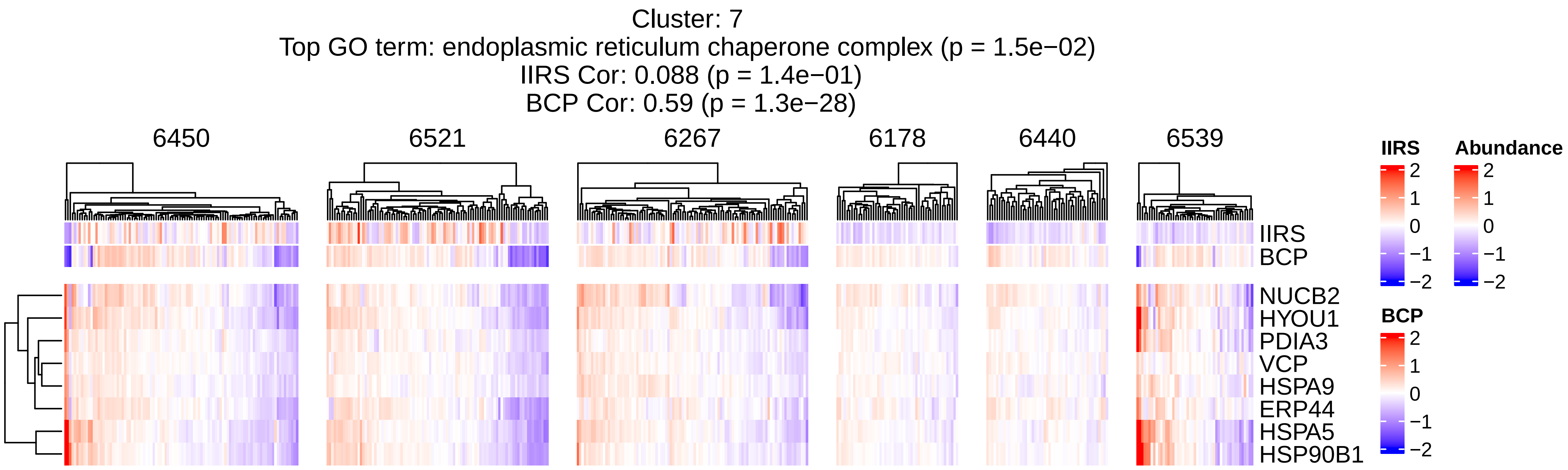
Cluster: 7
Top GO term: endoplasmic reticulum chaperone complex (p = 1.5e−02)
IIRS Cor: 0.088 (p = 1.4e−01)
BCP Cor: 0.59 (p = 1.3e−28)
6450
6521
6267
6178
6440
6539
IIRS
BCP
NUCB2
HYOU1
PDIA3
VCP
HSPA9
ERP44
HSPA5
HSP90B1
IIRS
2
1
0
−1
−2
Abundance
2
1
0
−1
−2
BCP
2
1
0
−1
−2

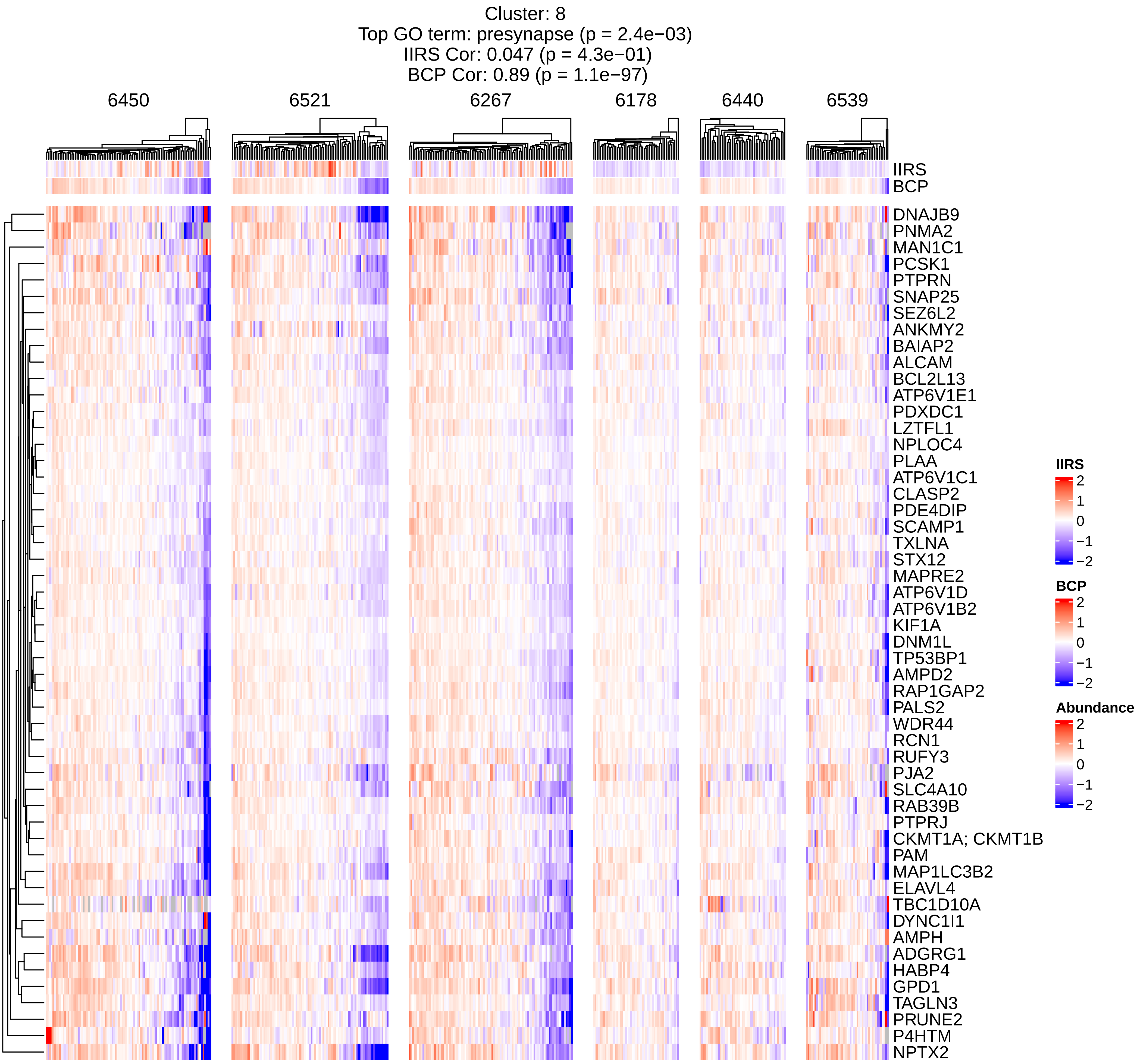

Cluster: 8
Top GO term: presynapse (p = 2.4e−03)
IIRS Cor: 0.047 (p = 4.3e−01)
BCP Cor: 0.89 (p = 1.1e−97)
6450
6521
6267
6178
6440
6539
IIRS
BCP
DNAJB9
PNMA2
MAN1C1
PCSK1
PTPRN
SNAP25
SEZ6L2
ANKMY2
BAIAP2
ALCAM
BCL2L13
ATP6V1E1
PDXDC1
LZTFL1
NPLOC4
PLAA
ATP6V1C1
CLASP2
PDE4DIP
SCAMP1
TXLNA
STX12
MAPRE2
ATP6V1D
ATP6V1B2
KIF1A
DNM1L
TP53BP1
AMPD2
RAP1GAP2
PALS2
WDR44
RCN1
RUFY3
PJA2
SLC4A10
RAB39B
PTPRJ
CKMT1A; CKMT1B
PAM
MAP1LC3B2
ELAVL4
TBC1D10A
DYNC1I1
AMPH
ADGRG1
HABP4
GPD1
TAGLN3
PRUNE2
P4HTM
NPTX2
IIRS
2
1
0
−1
−2
BCP
2
1
0
−1
−2
Abundance
2
1
0
−1
−2

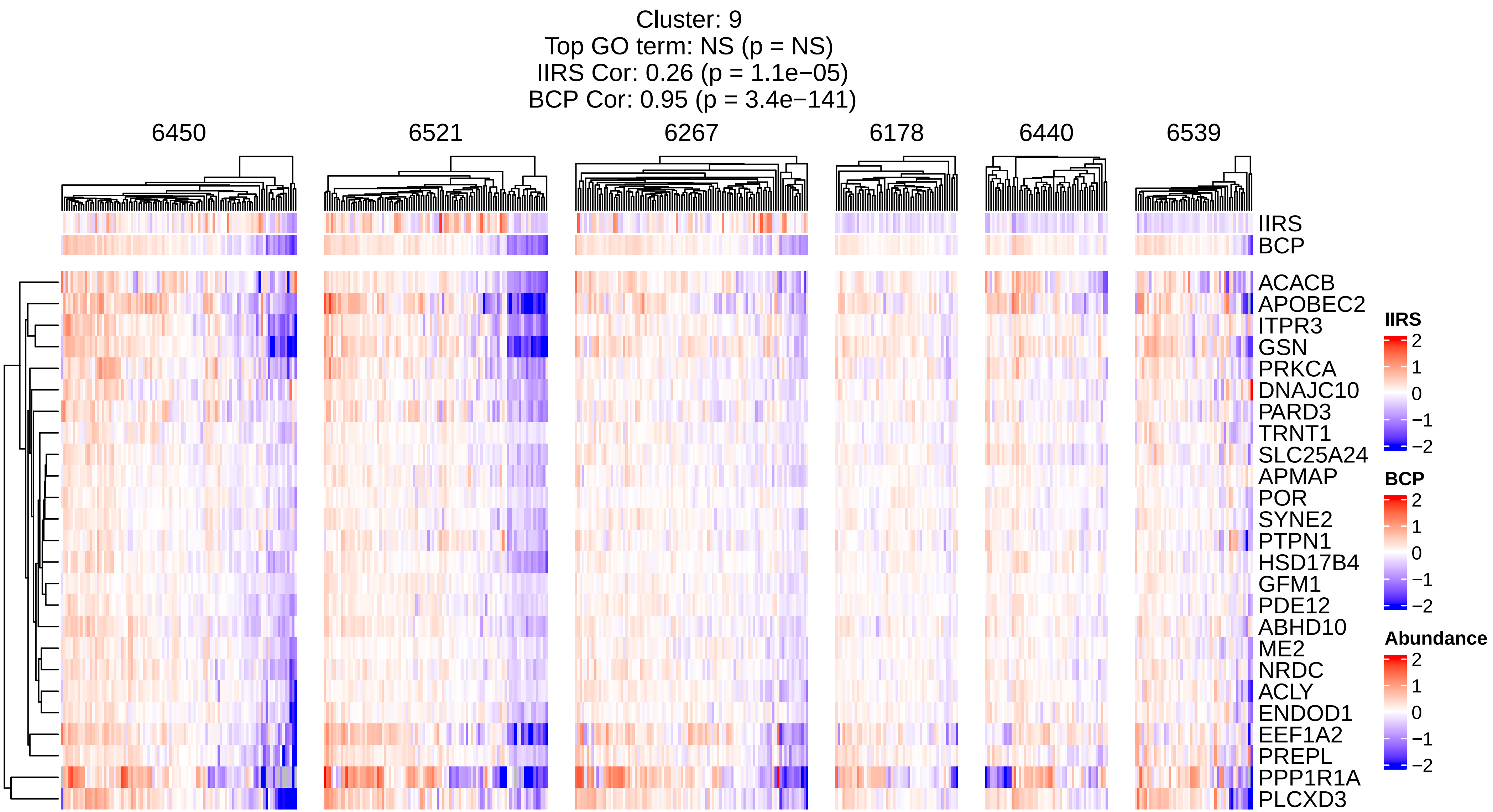

Cluster: 9
Top GO term: NS (p = NS)
IIRS Cor: 0.26 (p = 1.1e−05)
BCP Cor: 0.95 (p = 3.4e−141)
6450
6521
6267
6178
6440
6539
IIRS
BCP
ACACB
APOBEC2
ITPR3
GSN
PRKCA
DNAJC10
PARD3
TRNT1
SLC25A24
APMAP
POR
SYNE2
PTPN1
HSD17B4
GFM1
PDE12
ABHD10
ME2
NRDC
ACLY
ENDOD1
EEF1A2
PREPL
PPP1R1A
PLCXD3
IIRS
2
1
0
−1
−2
BCP
2
1
0
−1
−2
Abundance
2
1
0
−1
−2

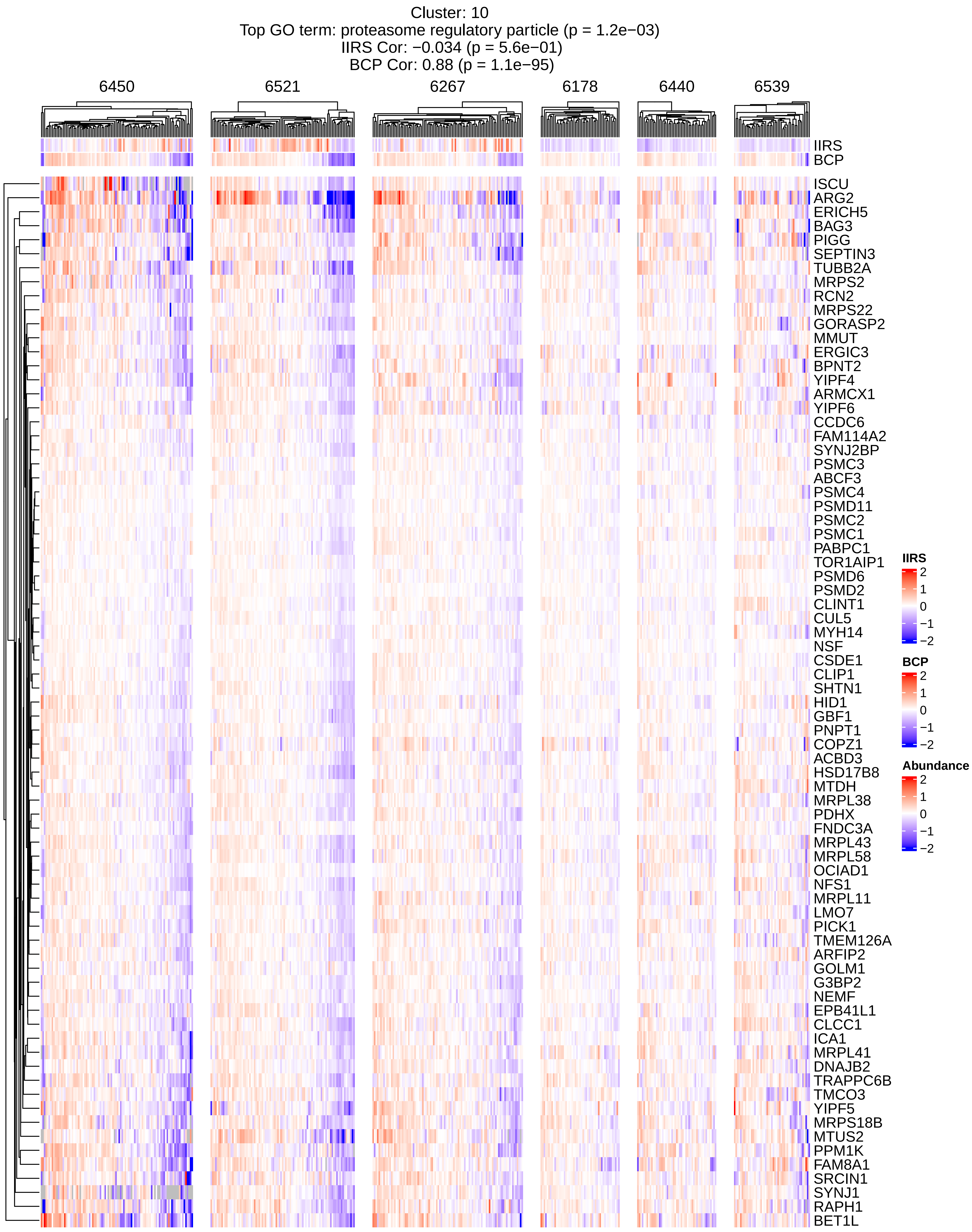
Cluster: 10
Top GO term: proteasome regulatory particle (p = 1.2e−03)
IIRS Cor: −0.034 (p = 5.6e−01)
BCP Cor: 0.88 (p = 1.1e−95)
6450
6521
6267
6178
6440
6539
IIRS
BCP
ISCU
ARG2
ERICH5
BAG3
PIGG
SEPTIN3
TUBB2A
MRPS2
RCN2
MRPS22
GORASP2
MMUT
ERGIC3
BPNT2
YIPF4
ARMCX1
YIPF6
CCDC6
FAM114A2
SYNJ2BP
PSMC3
ABCF3
PSMC4
PSMD11
PSMC2
PSMC1
PABPC1
TOR1AIP1
PSMD6
PSMD2
CLINT1
CUL5
MYH14
NSF
CSDE1
CLIP1
SHTN1
HID1
GBF1
PNPT1
COPZ1
ACBD3
HSD17B8
MTDH
MRPL38
PDHX
FNDC3A
MRPL43
MRPL58
OCIAD1
NFS1
MRPL11
LMO7
PICK1
TMEM126A
ARFIP2
GOLM1
G3BP2
NEMF
EPB41L1
CLCC1
ICA1
MRPL41
DNAJB2
TRAPPC6B
TMCO3
YIPF5
MRPS18B
MTUS2
PPM1K
FAM8A1
SRCIN1
SYNJ1
RAPH1
BET1L
IIRS
2
1
0
−1
−2
BCP
2
1
0
−1
−2
Abundance
2
1
0
−1
−2

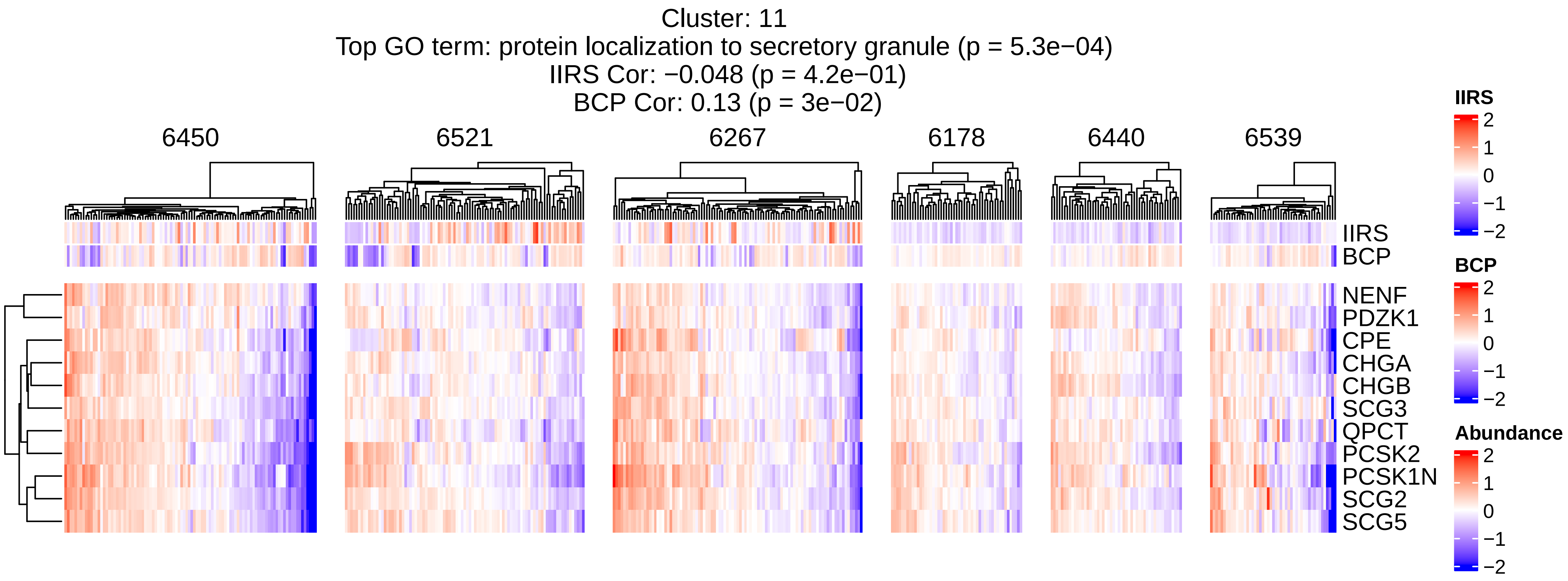

Cluster: 11
Top GO term: protein localization to secretory granule (p = 5.3e−04)
IIRS Cor: −0.048 (p = 4.2e−01)
BCP Cor: 0.13 (p = 3e−02)
6450
6521
6267
6178
6440
6539
IIRS
BCP
NENF
PDZK1
CPE
CHGA
CHGB
SCG3
QPCT
PCSK2
PCSK1N
SCG2
SCG5
IIRS
2
1
0
−1
−2
BCP
2
1
0
−1
−2
Abundance
2
1
0
−1
−2

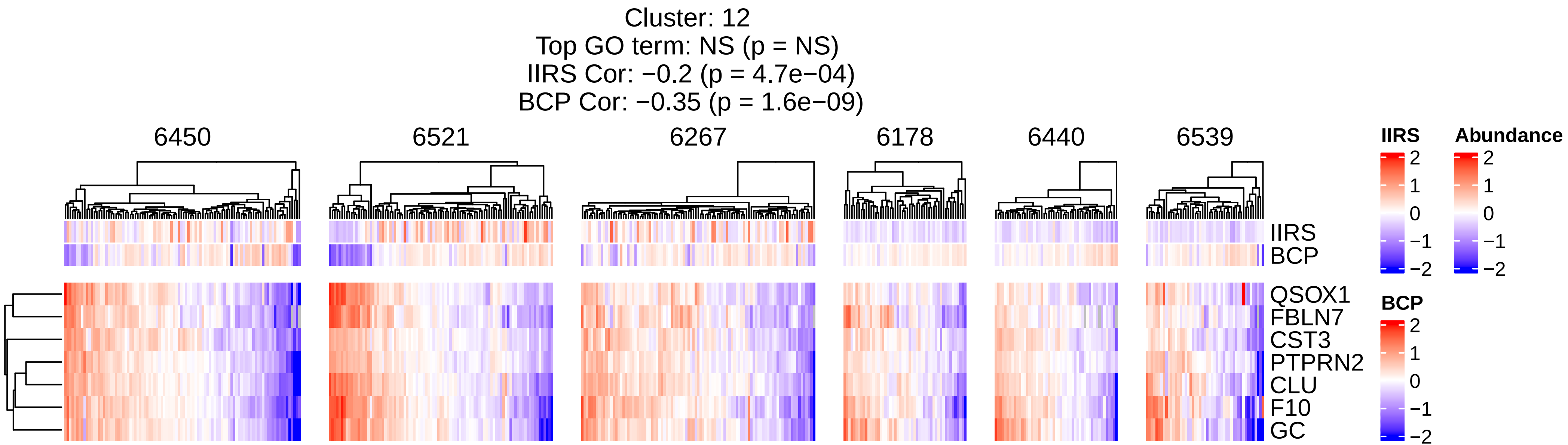
Cluster: 12
Top GO term: NS (p = NS)
IIRS Cor: −0.2 (p = 4.7e−04)
BCP Cor: −0.35 (p = 1.6e−09)
6450
6521
6267
6178
6440
6539
IIRS
BCP
QSOX1
FBLN7
CST3
PTPRN2
CLU
F10
GC
IIRS
2
1
0
−1
−2
Abundance
2
1
0
−1
−2
BCP
2
1
0
−1
−2

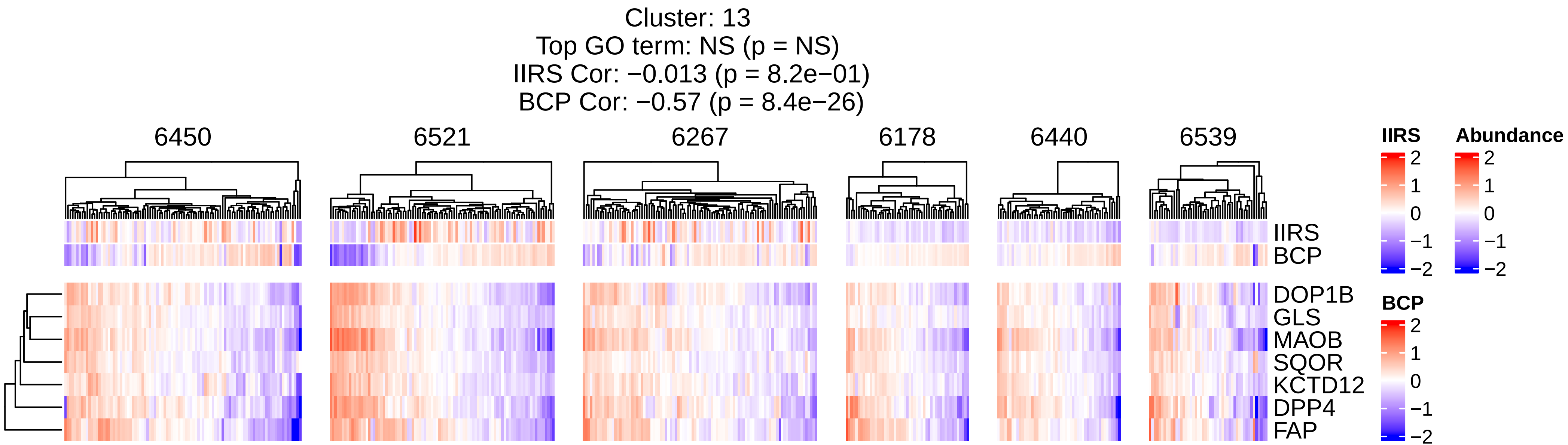

Cluster: 13
Top GO term: NS (p = NS)
IIRS Cor: −0.013 (p = 8.2e−01)
BCP Cor: −0.57 (p = 8.4e−26)
6450
6521
6267
6178
6440
6539
IIRS
BCP
DOP1B
GLS
MAOB
SQOR
KCTD12
DPP4
FAP
IIRS
2
1
0
−1
−2
Abundance
2
1
0
−1
−2
BCP
2
1
0
−1
−2

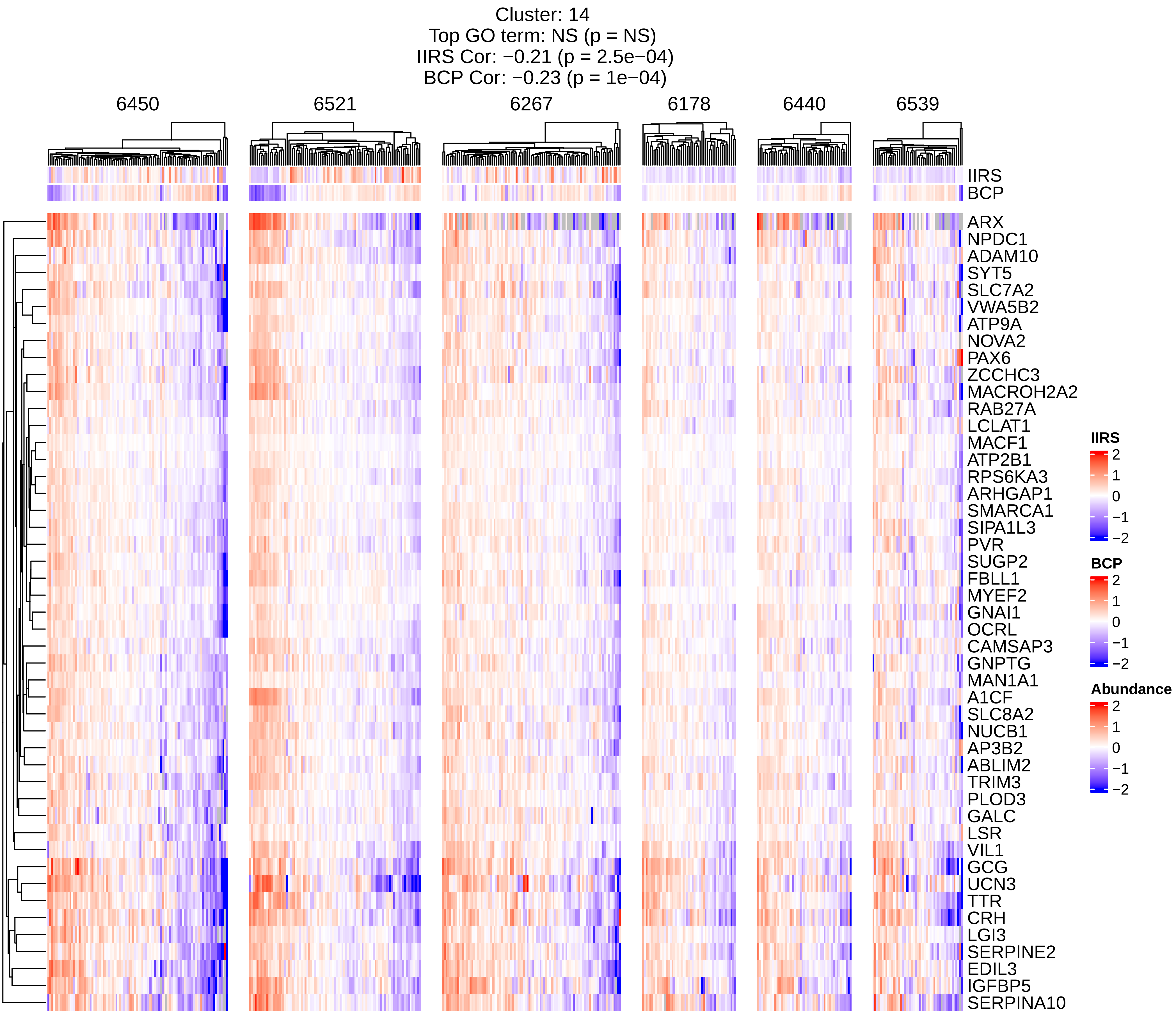
Cluster: 14
Top GO term: NS (p = NS)
IIRS Cor: −0.21 (p = 2.5e−04)
BCP Cor: −0.23 (p = 1e−04)
6450
6521
6267
6178
6440
6539
IIRS
BCP
ARX
NPDC1
ADAM10
SYT5
SLC7A2
VWA5B2
ATP9A
NOVA2
PAX6
ZCCHC3
MACROH2A2
RAB27A
LCLAT1
MACF1
ATP2B1
RPS6KA3
ARHGAP1
SMARCA1
SIPA1L3
PVR
SUGP2
FBLL1
MYEF2
GNAI1
OCRL
CAMSAP3
GNPTG
MAN1A1
A1CF
SLC8A2
NUCB1
AP3B2
ABLIM2
TRIM3
PLOD3
GALC
LSR
VIL1
GCG
UCN3
TTR
CRH
LGI3
SERPINE2
EDIL3
IGFBP5
SERPINA10
IIRS
2
1
0
−1
−2
BCP
2
1
0
−1
−2
Abundance
2
1
0
−1
−2

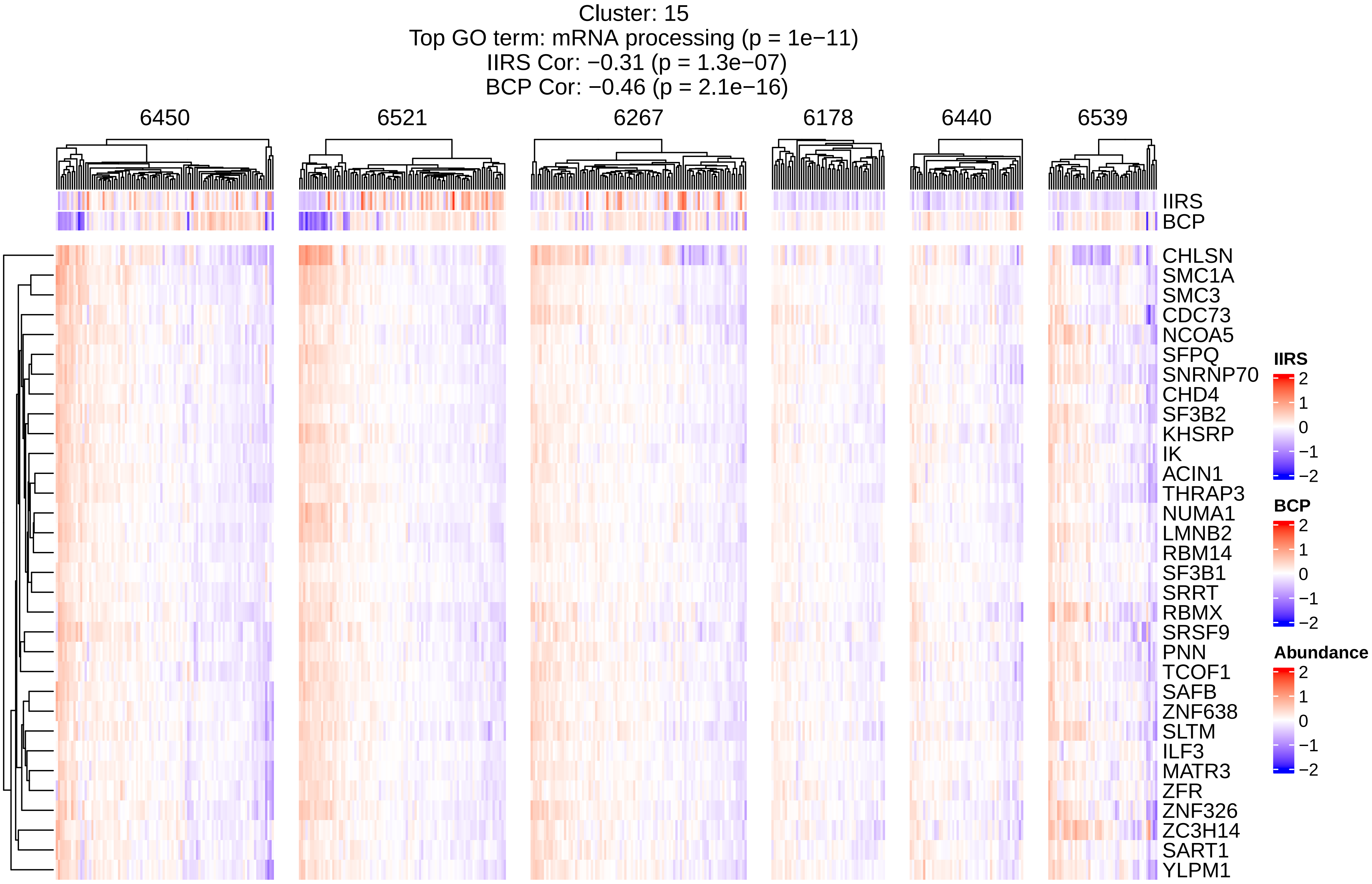
Cluster: 15
Top GO term: mRNA processing (p = 1e−11)
IIRS Cor: −0.31 (p = 1.3e−07)
BCP Cor: −0.46 (p = 2.1e−16)
6450
6521
6267
6178
6440
6539
IIRS
BCP
CHLSN
SMC1A
SMC3
CDC73
NCOA5
SFPQ
SNRNP70
CHD4
SF3B2
KHSRP
IK
ACIN1
THRAP3
NUMA1
LMNB2
RBM14
SF3B1
SRRT
RBMX
SRSF9
PNN
TCOF1
SAFB
ZNF638
SLTM
ILF3
MATR3
ZFR
ZNF326
ZC3H14
SART1
YLPM1
IIRS
2
1
0
−1
−2
BCP
2
1
0
−1
−2
Abundance
2
1
0
−1
−2

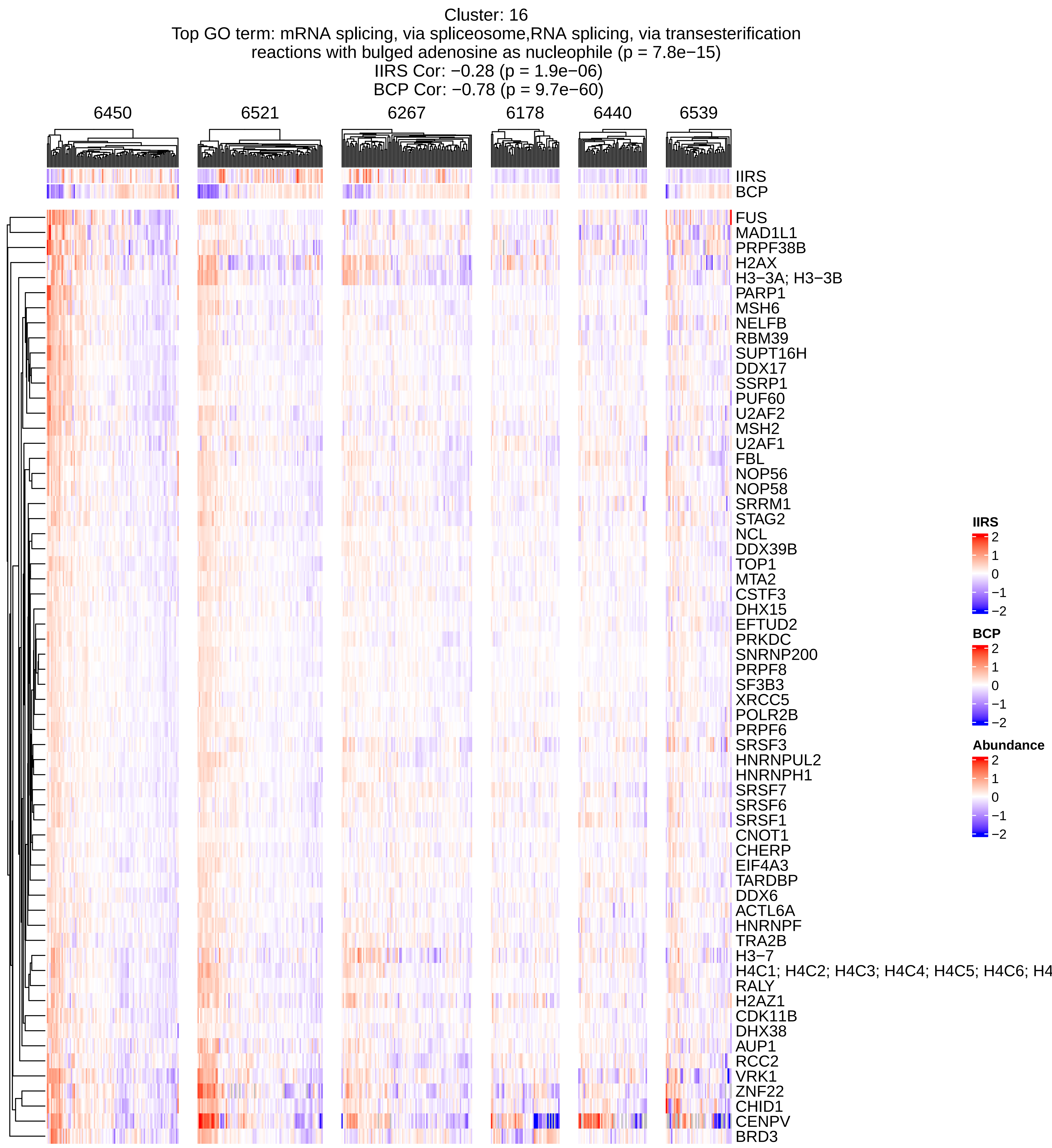
Cluster: 16
Top GO term: mRNA splicing, via spliceosome,RNA splicing, via transesterification reactions with bulged adenosine as nucleophile (p = 7.8e−15)
IIRS Cor: −0.28 (p = 1.9e−06)
BCP Cor: −0.78 (p = 9.7e−60)
6450
6521
6267
6178
6440
6539
IIRS
BCP
FUS
MAD1L1
PRPF38B
H2AX
H3−3A; H3−3B
PARP1
MSH6
NELFB
RBM39
SUPT16H
DDX17
SSRP1
PUF60
U2AF2
MSH2
U2AF1
FBL
NOP56
NOP58
SRRM1
STAG2
NCL
DDX39B
TOP1
MTA2
CSTF3
DHX15
EFTUD2
PRKDC
SNRNP200
PRPF8
SF3B3
XRCC5
POLR2B
PRPF6
SRSF3
HNRNPUL2
HNRNPH1
SRSF7
SRSF6
SRSF1
CNOT1
CHERP
EIF4A3
TARDBP
DDX6
ACTL6A
HNRNPF
TRA2B
H3−7
H4C1; H4C2; H4C3; H4C4; H4C5; H4C6; H4
RALY
H2AZ1
CDK11B
DHX38
AUP1
RCC2
VRK1
ZNF22
CHID1
CENPV
BRD3
IIRS
2
1
0
−1
−2
BCP
2
1
0
−1
−2
Abundance
2
1
0
−1
−2

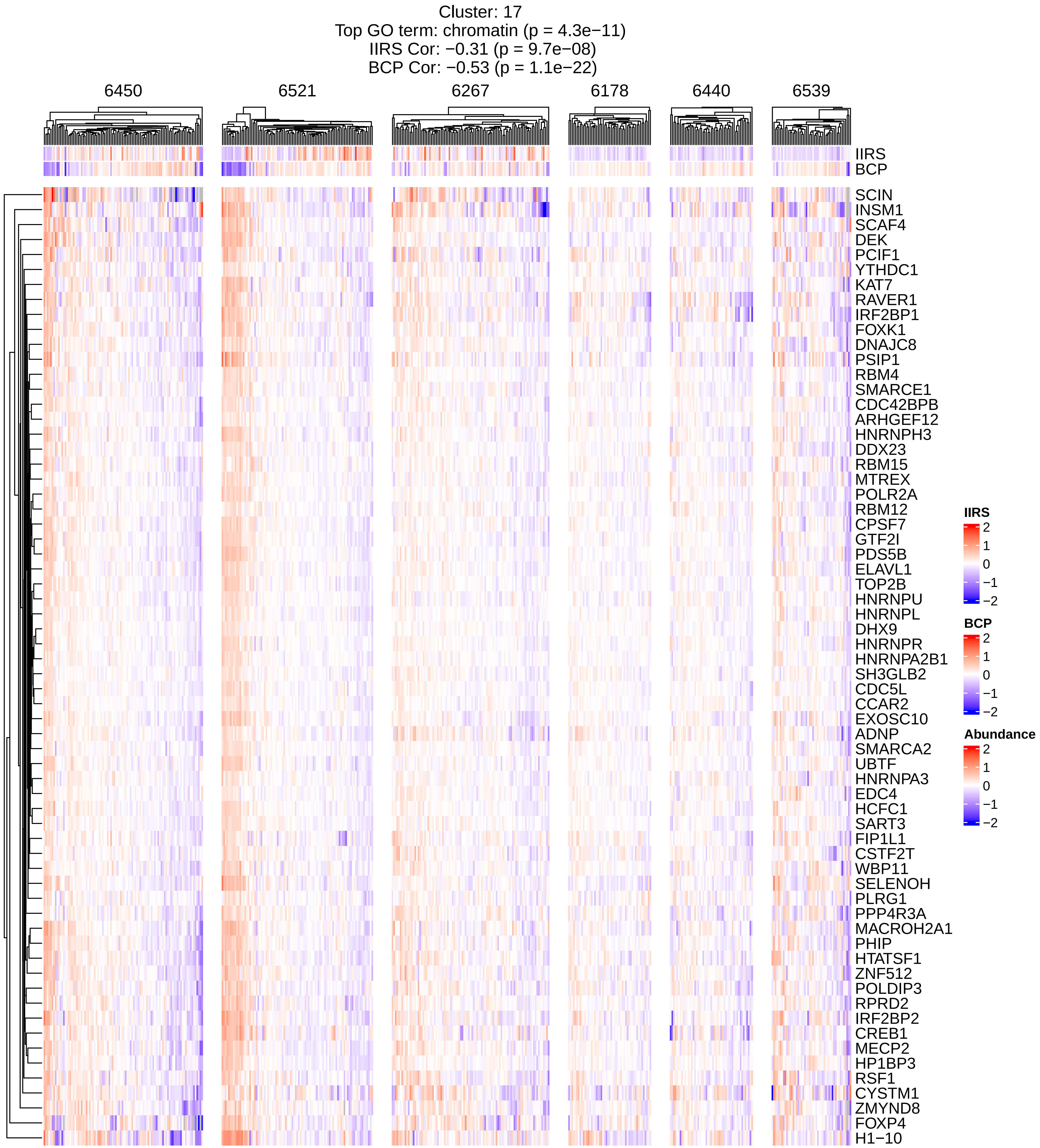
Cluster: 17
Top GO term: chromatin (p = 4.3e−11)
IIRS Cor: −0.31 (p = 9.7e−08)
BCP Cor: −0.53 (p = 1.1e−22)
6450
6521
6267
6178
6440
6539
IIRS
BCP
SCIN
INSM1
SCAF4
DEK
PCIF1
YTHDC1
KAT7
RAVER1
IRF2BP1
FOXK1
DNAJC8
PSIP1
RBM4
SMARCE1
CDC42BPB
ARHGEF12
HNRNPH3
DDX23
RBM15
MTREX
POLR2A
RBM12
CPSF7
GTF2I
PDS5B
ELAVL1
TOP2B
HNRNPU
HNRNPL
DHX9
HNRNPR
HNRNPA2B1
SH3GLB2
CDC5L
CCAR2
EXOSC10
ADNP
SMARCA2
UBTF
HNRNPA3
EDC4
HCFC1
SART3
FIP1L1
CSTF2T
WBP11
SELENOH
PLRG1
PPP4R3A
MACROH2A1
PHIP
HTATSF1
ZNF512
POLDIP3
RPRD2
IRF2BP2
CREB1
MECP2
HP1BP3
RSF1
CYSTM1
ZMYND8
FOXP4
H1−10
IIRS
2
1
0
−1
−2
BCP
2
1
0
−1
−2
Abundance
2
1
0
−1
−2

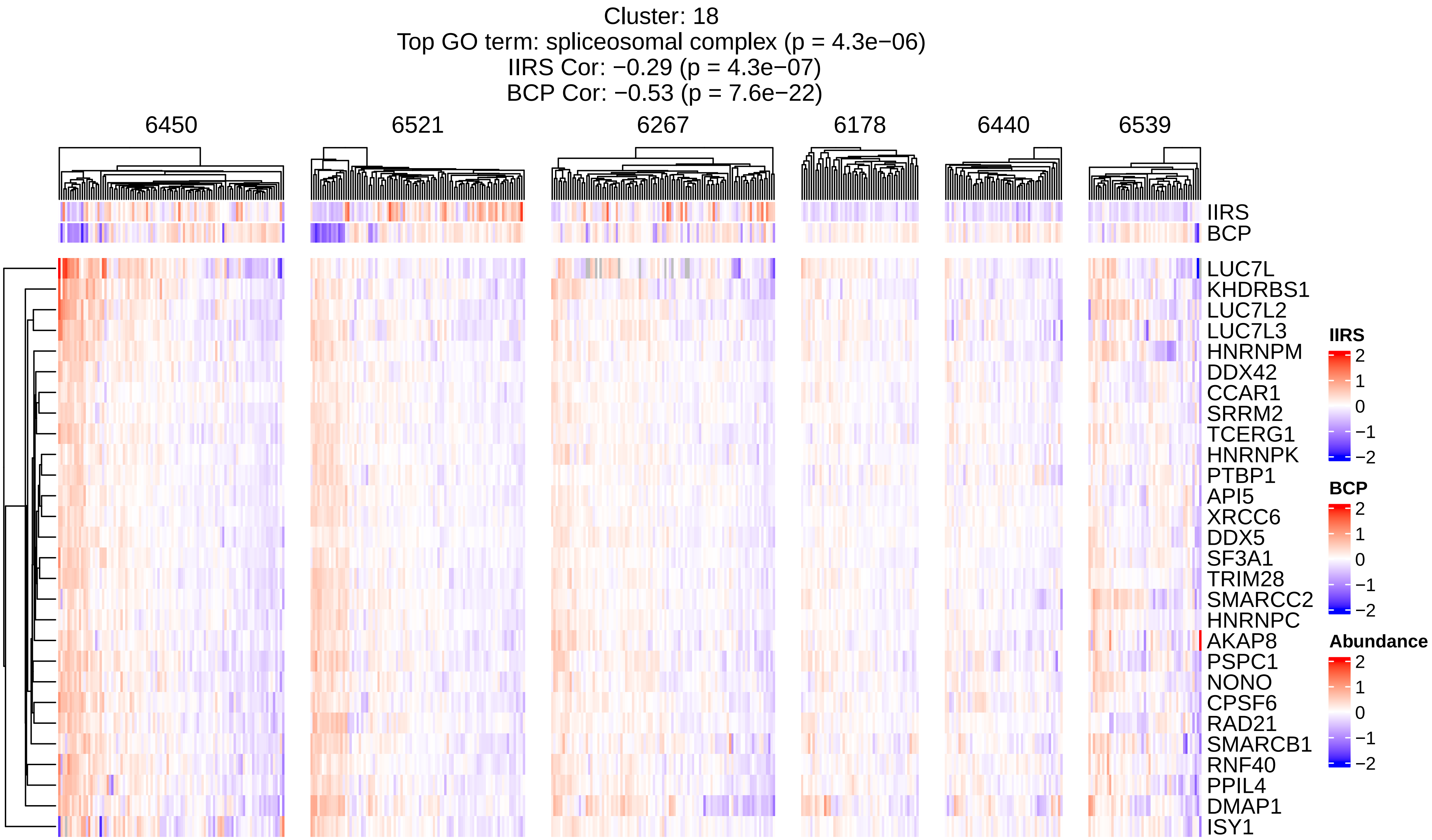
Cluster: 18
Top GO term: spliceosomal complex (p = 4.3e−06)
IIRS Cor: −0.29 (p = 4.3e−07)
BCP Cor: −0.53 (p = 7.6e−22)
6450
6521
6267
6178
6440
6539
IIRS
BCP
LUC7L
KHDRBS1
LUC7L2
LUC7L3
HNRNPM
DDX42
CCAR1
SRRM2
TCERG1
HNRNPK
PTBP1
API5
XRCC6
DDX5
SF3A1
TRIM28
SMARCC2
HNRNPC
AKAP8
PSPC1
NONO
CPSF6
RAD21
SMARCB1
RNF40
PPIL4
DMAP1
ISY1
IIRS
2
1
0
−1
−2
BCP
2
1
0
−1
−2
Abundance
2
1
0
−1
−2

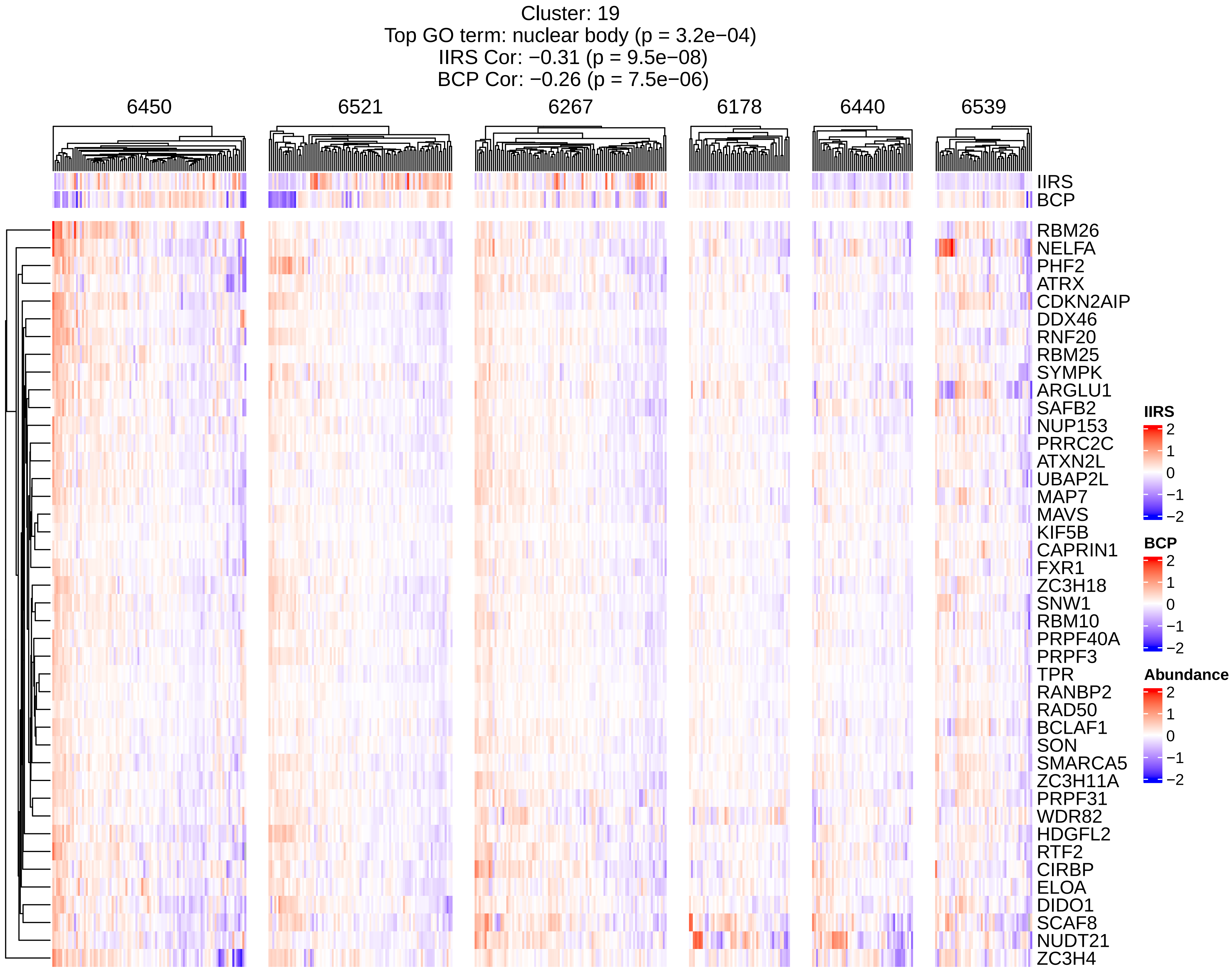

Cluster: 19
Top GO term: nuclear body (p = 3.2e−04)
IIRS Cor: −0.31 (p = 9.5e−08)
BCP Cor: −0.26 (p = 7.5e−06)
6450
6521
6267
6178
6440
6539
IIRS
BCP
RBM26
NELFA
PHF2
ATRX
CDKN2AIP
DDX46
RNF20
RBM25
SYMPK
ARGLU1
SAFB2
NUP153
PRRC2C
ATXN2L
UBAP2L
MAP7
MAVS
KIF5B
CAPRIN1
FXR1
ZC3H18
SNW1
RBM10
PRPF40A
PRPF3
TPR
RANBP2
RAD50
BCLAF1
SON
SMARCA5
ZC3H11A
PRPF31
WDR82
HDGFL2
RTF2
CIRBP
ELOA
DIDO1
SCAF8
NUDT21
ZC3H4
IIRS
2
1
0
−1
−2
BCP
2
1
0
−1
−2
Abundance
2
1
0
−1
−2

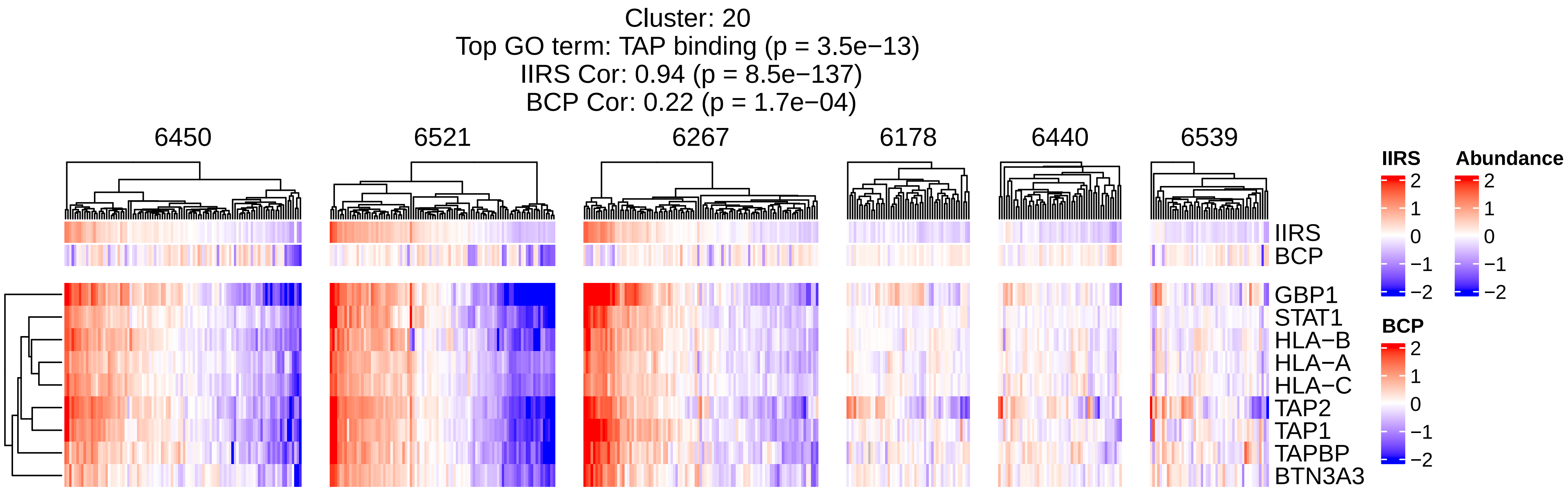

Cluster: 20
Top GO term: TAP binding (p = 3.5e−13)
IIRS Cor: 0.94 (p = 8.5e−137)
BCP Cor: 0.22 (p = 1.7e−04)
6450
6521
6267
6178
6440
6539
IIRS
BCP
GBP1
STAT1
HLA−B
HLA−A
HLA−C
TAP2
TAP1
TAPBP
BTN3A3
IIRS
2
1
0
−1
−2
Abundance
2
1
0
−1
−2
BCP
2
1
0
−1
−2

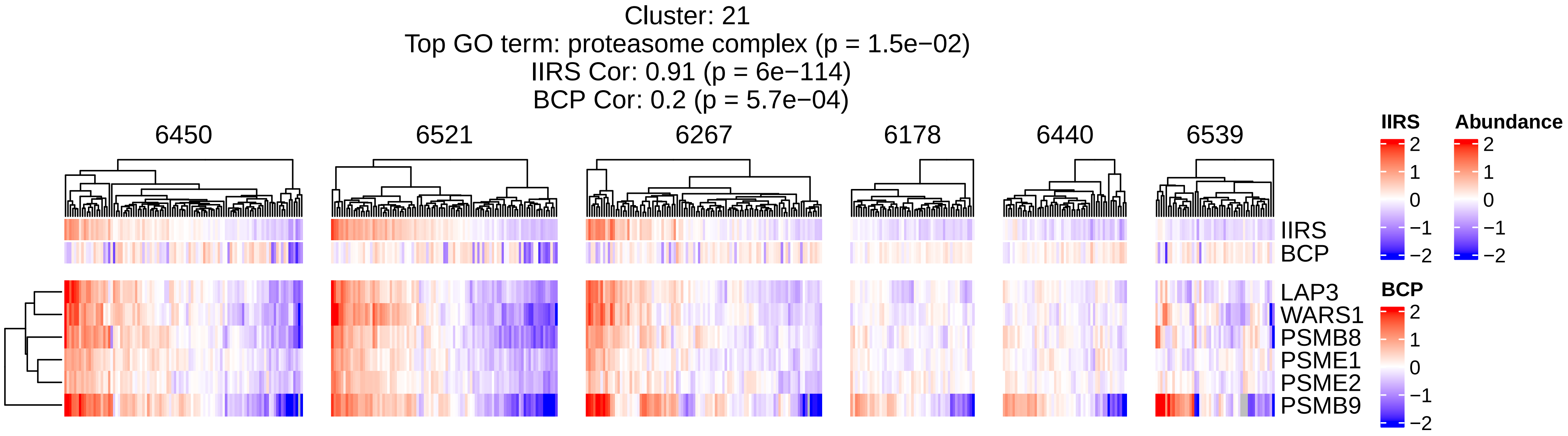

Cluster: 21
Top GO term: proteasome complex (p = 1.5e−02)
IIRS Cor: 0.91 (p = 6e−114)
BCP Cor: 0.2 (p = 5.7e−04)
6450
6521
6267
6178
6440
6539
IIRS
BCP
LAP3
WARS1
PSMB8
PSME1
PSME2
PSMB9
IIRS
2
1
0
−1
−2
Abundance
2
1
0
−1
−2
BCP
2
1
0
−1
−2

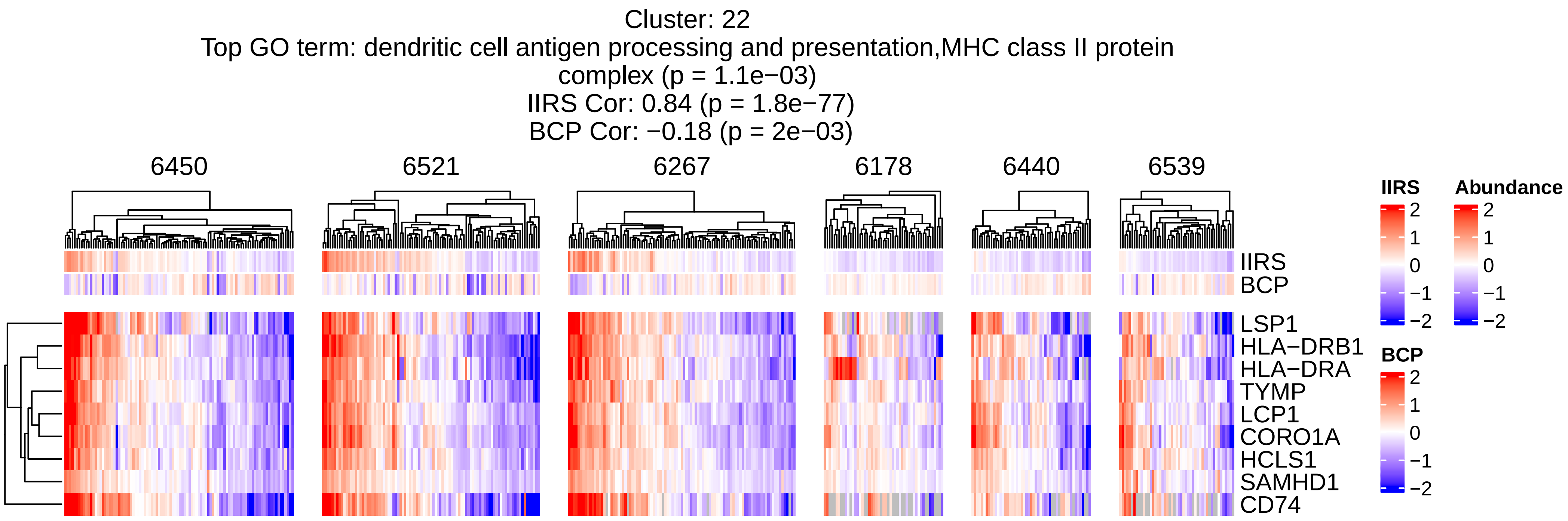
Cluster: 22
Top GO term: dendritic cell antigen processing and presentation,MHC class II protein complex (p = 1.1e−03)
IIRS Cor: 0.84 (p = 1.8e−77)
BCP Cor: −0.18 (p = 2e−03)
6450
6521
6267
6178
6440
6539
IIRS
BCP
LSP1
HLA−DRB1
HLA−DRA
TYMP
LCP1
CORO1A
HCLS1
SAMHD1
CD74
IIRS
2
1
0
−1
−2
Abundance
2
1
0
−1
−2
BCP
2
1
0
−1
−2

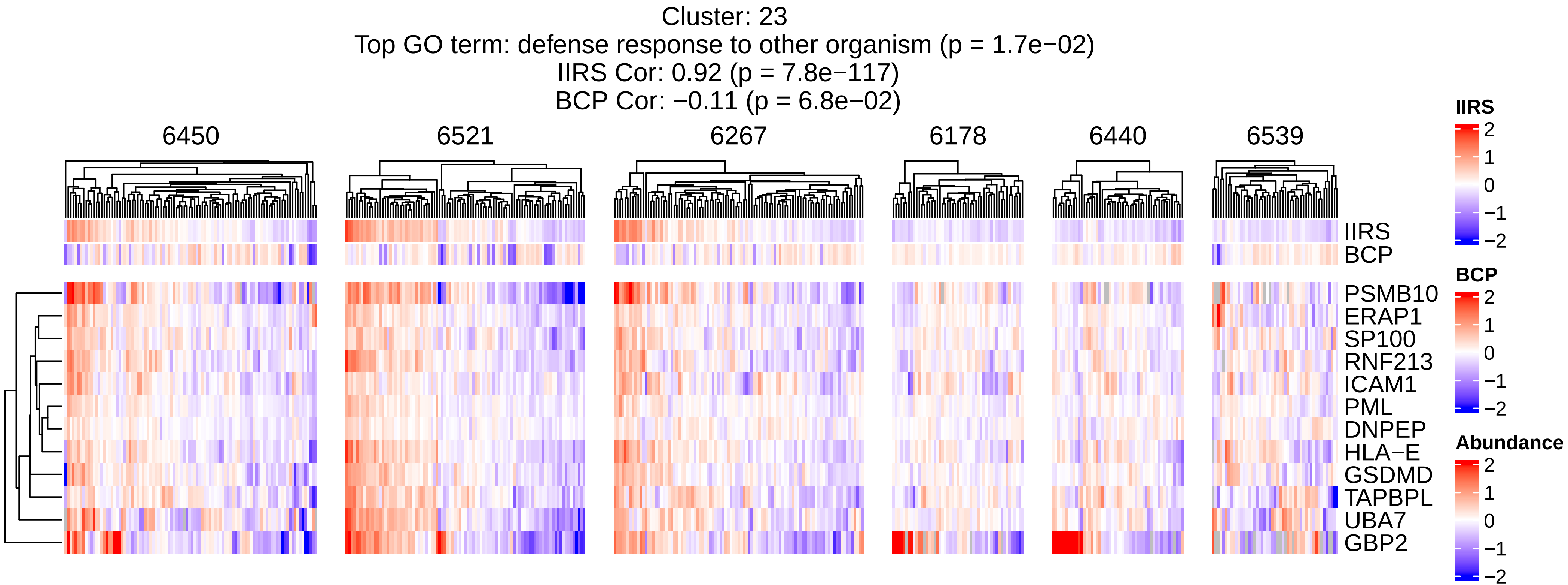

Cluster: 23
Top GO term: defense response to other organism (p = 1.7e−02)
IIRS Cor: 0.92 (p = 7.8e−117)
BCP Cor: −0.11 (p = 6.8e−02)
6450
6521
6267
6178
6440
6539
IIRS
BCP
PSMB10
ERAP1
SP100
RNF213
ICAM1
PML
DNPEP
HLA−E
GSDMD
TAPBPL
UBA7
GBP2
IIRS
2
1
0
−1
−2
BCP
2
1
0
−1
−2
Abundance
2
1
0
−1
−2

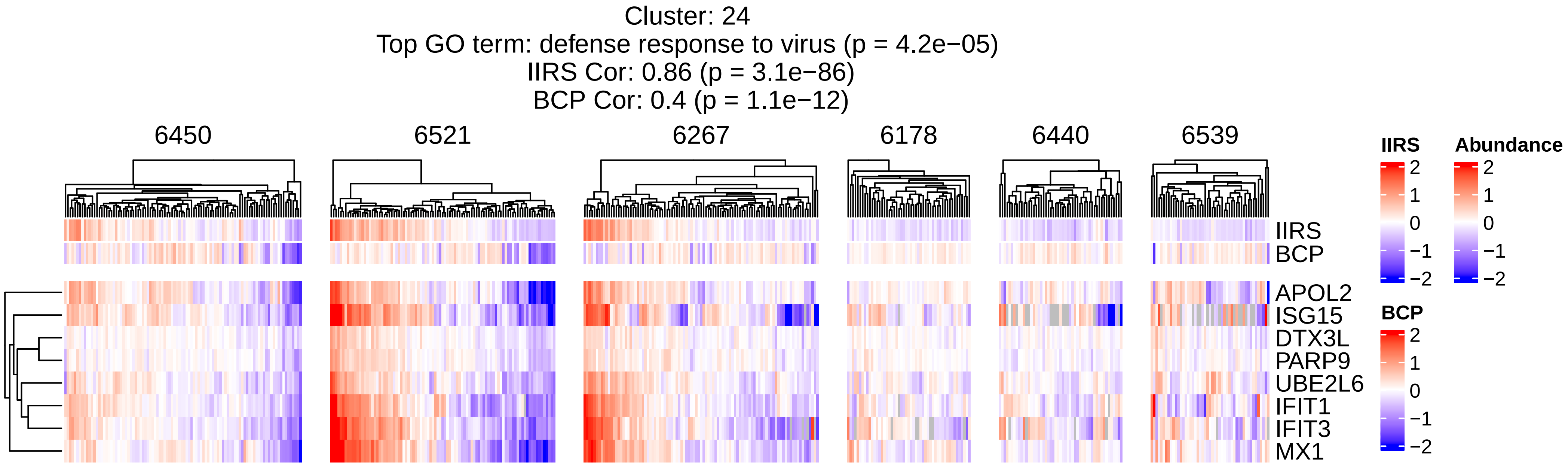
Cluster: 24
Top GO term: defense response to virus (p = 4.2e−05)
IIRS Cor: 0.86 (p = 3.1e−86)
BCP Cor: 0.4 (p = 1.1e−12)
6450
6521
6267
6178
6440
6539
IIRS
BCP
APOL2
ISG15
DTX3L
PARP9
UBE2L6
IFIT1
IFIT3
MX1
IIRS
2
1
0
−1
−2
Abundance
2
1
0
−1
−2
BCP
2
1
0
−1
−2

Cluster: 25
Top GO term: oligosaccharyltransferase complex (p = 9.9e−08)
IIRS Cor: −0.0097 (p = 8.7e−01)
BCP Cor: −0.11 (p = 6.2e−02)

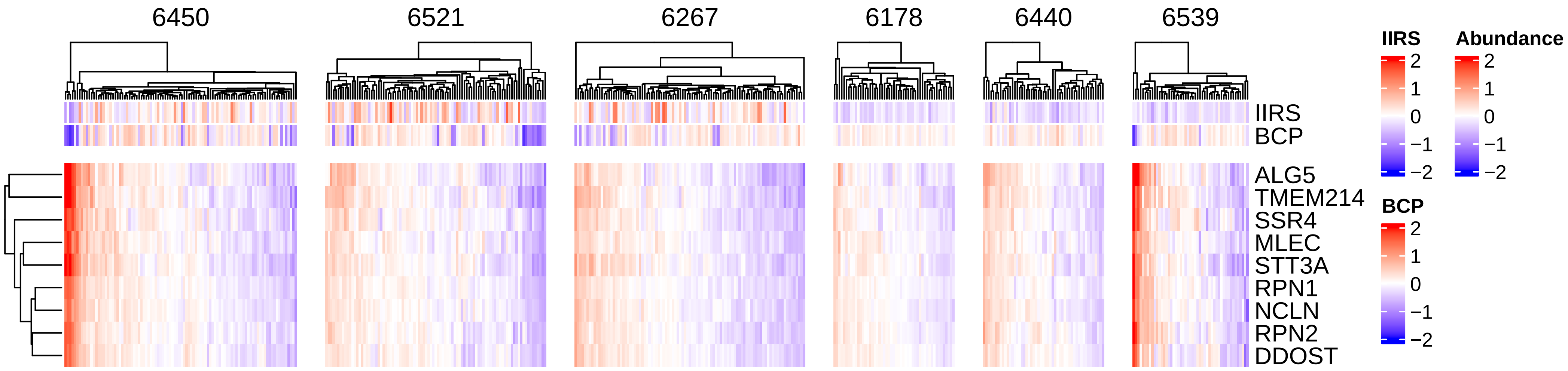

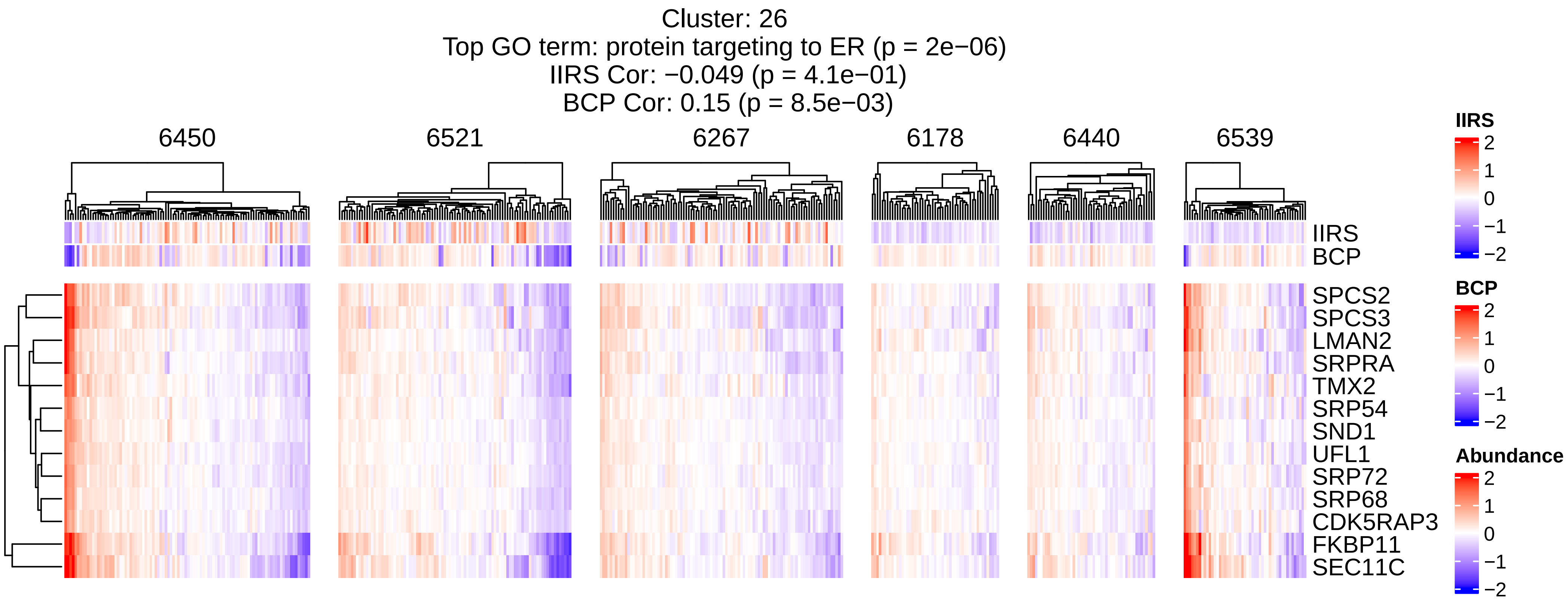
Cluster: 26
Top GO term: protein targeting to ER (p = 2e−06)
IIRS Cor: −0.049 (p = 4.1e−01)
BCP Cor: 0.15 (p = 8.5e−03)
6450
6521
6267
6178
6440
6539
IIRS
BCP
SPCS2
SPCS3
LMAN2
SRPRA
TMX2
SRP54
SND1
UFL1
SRP72
SRP68
CDK5RAP3
FKBP11
SEC11C
IIRS
2
1
0
−1
−2
BCP
2
1
0
−1
−2
Abundance
2
1
0
−1
−2

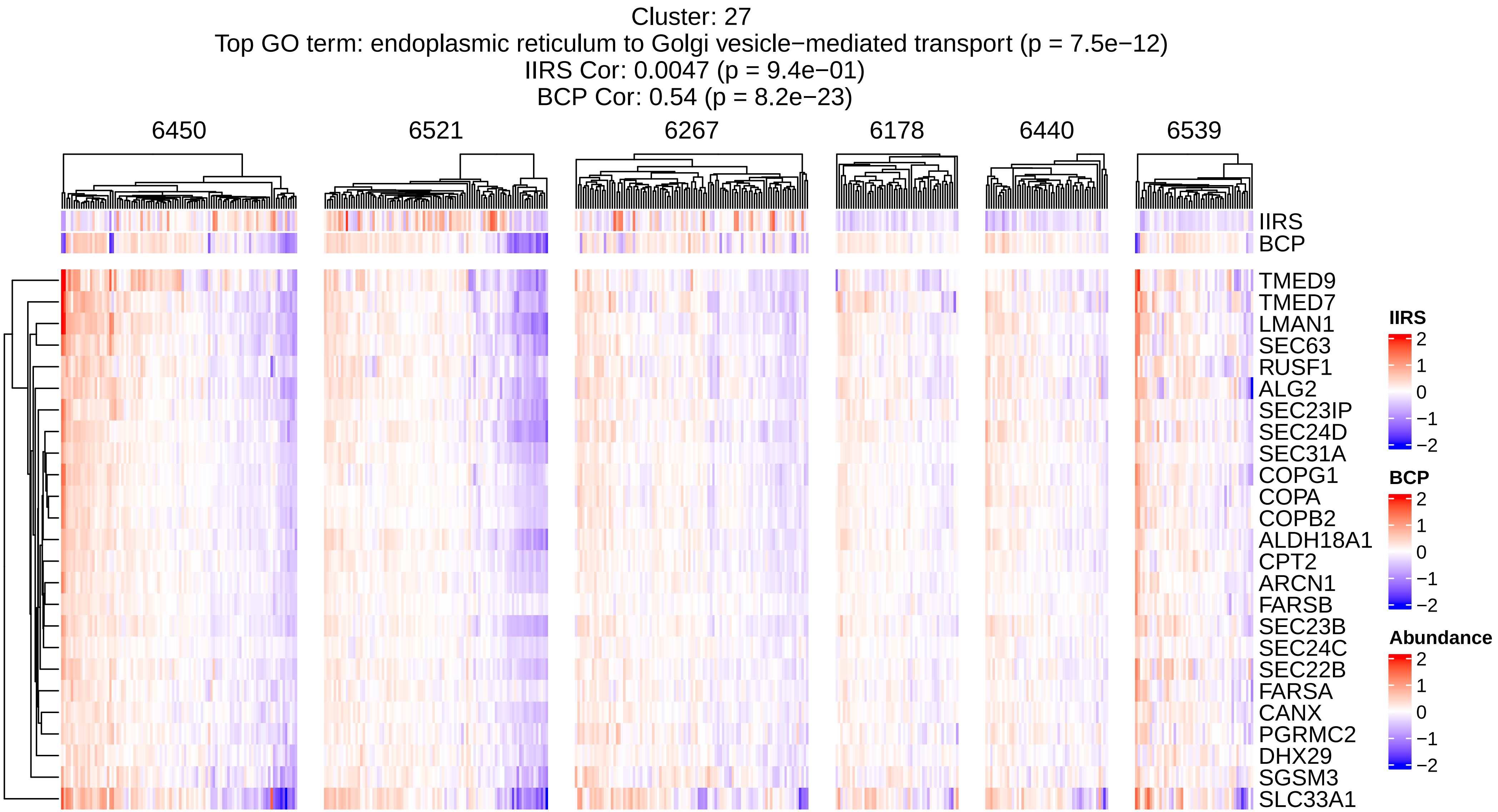
Cluster: 27
Top GO term: endoplasmic reticulum to Golgi vesicle−mediated transport (p = 7.5e−12)
IIRS Cor: 0.0047 (p = 9.4e−01)
BCP Cor: 0.54 (p = 8.2e−23)
6450
6521
6267
6178
6440
6539
IIRS
BCP
TMED9
TMED7
LMAN1
SEC63
RUSF1
ALG2
SEC23IP
SEC24D
SEC31A
COPG1
COPA
COPB2
ALDH18A1
CPT2
ARCN1
FARSB
SEC23B
SEC24C
SEC22B
FARSA
CANX
PGRMC2
DHX29
SGSM3
SLC33A1
IIRS
2
1
0
−1
−2
BCP
2
1
0
−1
−2
Abundance
2
1
0
−1
−2

Cluster: 28
Top GO term: protein disulfide isomerase activity,intramolecular oxidoreductase activity, transposing S−S bonds (p = 3.3e−02)
IIRS Cor: 0.1 (p = 8.5e−02)
BCP Cor: 0.25 (p = 1.8e−05)

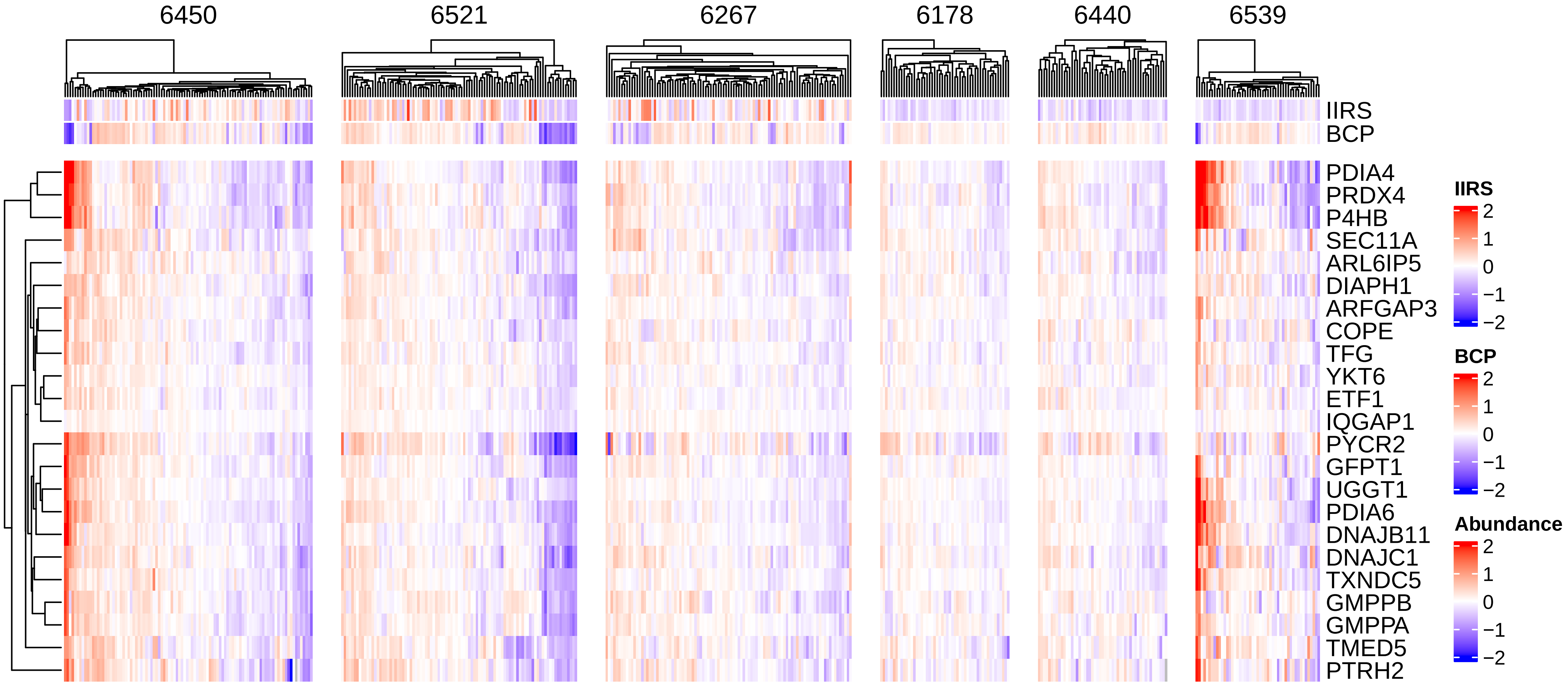

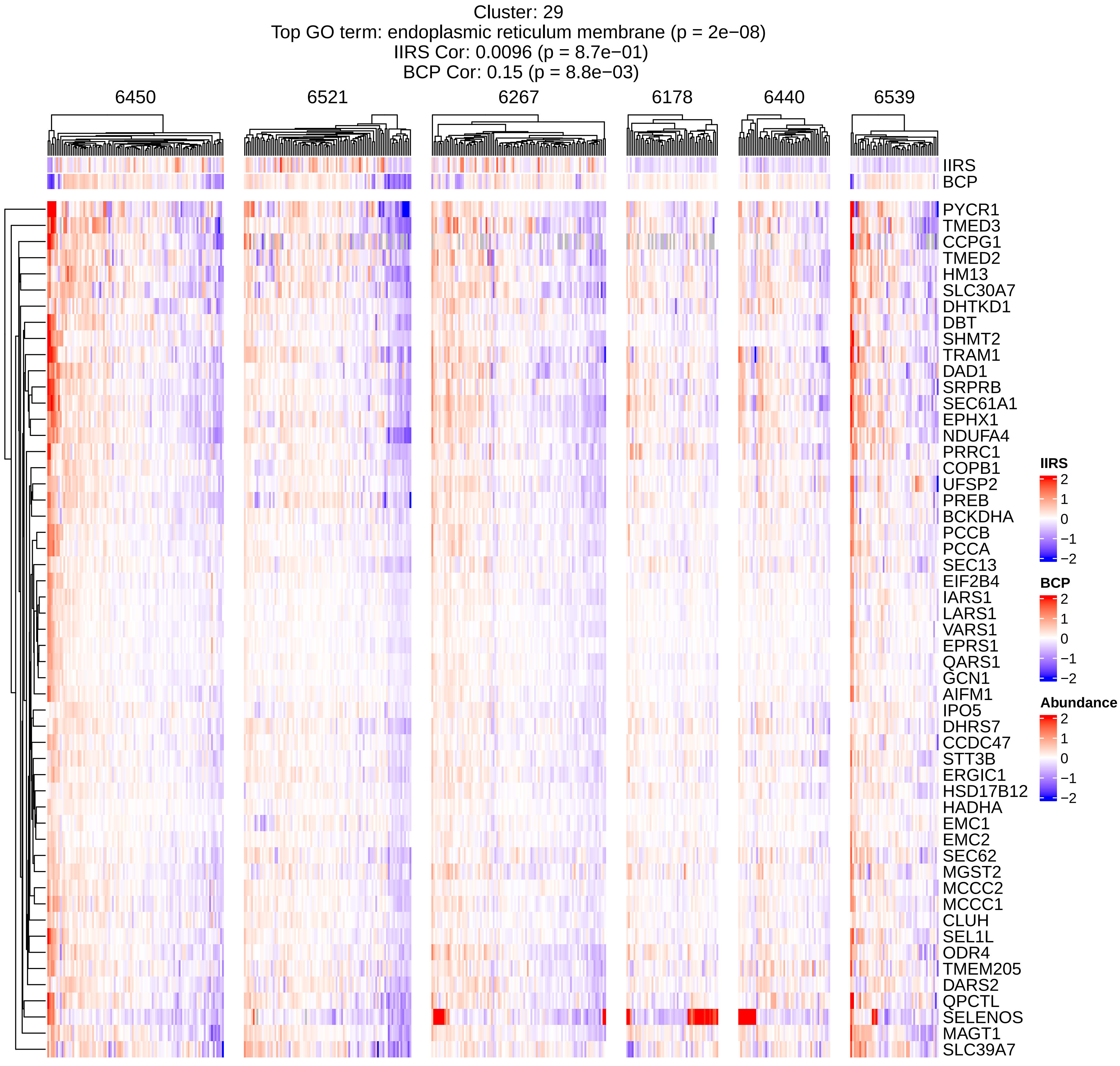
Cluster: 29
Top GO term: endoplasmic reticulum membrane (p = 2e−08)
IIRS Cor: 0.0096 (p = 8.7e−01)
BCP Cor: 0.15 (p = 8.8e−03)
6450
6521
6267
6178
6440
6539
IIRS
BCP
PYCR1
TMED3
CCPG1
TMED2
HM13
SLC30A7
DHTKD1
DBT
SHMT2
TRAM1
DAD1
SRPRB
SEC61A1
EPHX1
NDUFA4
PRRC1
COPB1
UFSP2
PREB
BCKDHA
PCCB
PCCA
SEC13
EIF2B4
IARS1
LARS1
VARS1
EPRS1
QARS1
GCN1
AIFM1
IPO5
DHRS7
CCDC47
STT3B
ERGIC1
HSD17B12
HADHA
EMC1
EMC2
SEC62
MGST2
MCCC2
MCCC1
CLUH
SEL1L
ODR4
TMEM205
DARS2
QPCTL
SELENOS
MAGT1
SLC39A7
IIRS
2
1
0
−1
−2
BCP
2
1
0
−1
−2
Abundance
2
1
0
−1
−2

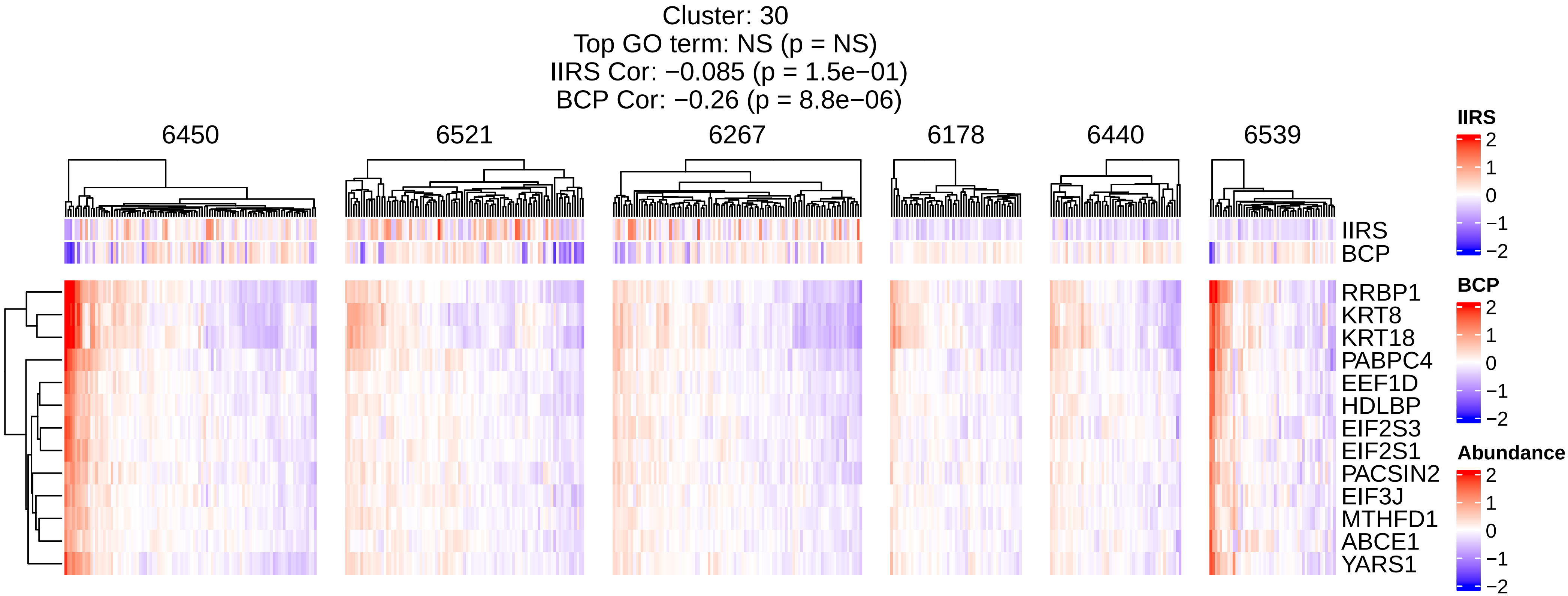
Cluster: 30
Top GO term: NS (p = NS)
IIRS Cor: −0.085 (p = 1.5e−01)
BCP Cor: −0.26 (p = 8.8e−06)
6450
6521
6267
6178
6440
6539
IIRS
BCP
RRBP1
KRT8
KRT18
PABPC4
EEF1D
HDLBP
EIF2S3
EIF2S1
PACSIN2
EIF3J
MTHFD1
ABCE1
YARS1
IIRS
2
1
0
−1
−2
BCP
2
1
0
−1
−2
Abundance
2
1
0
−1
−2

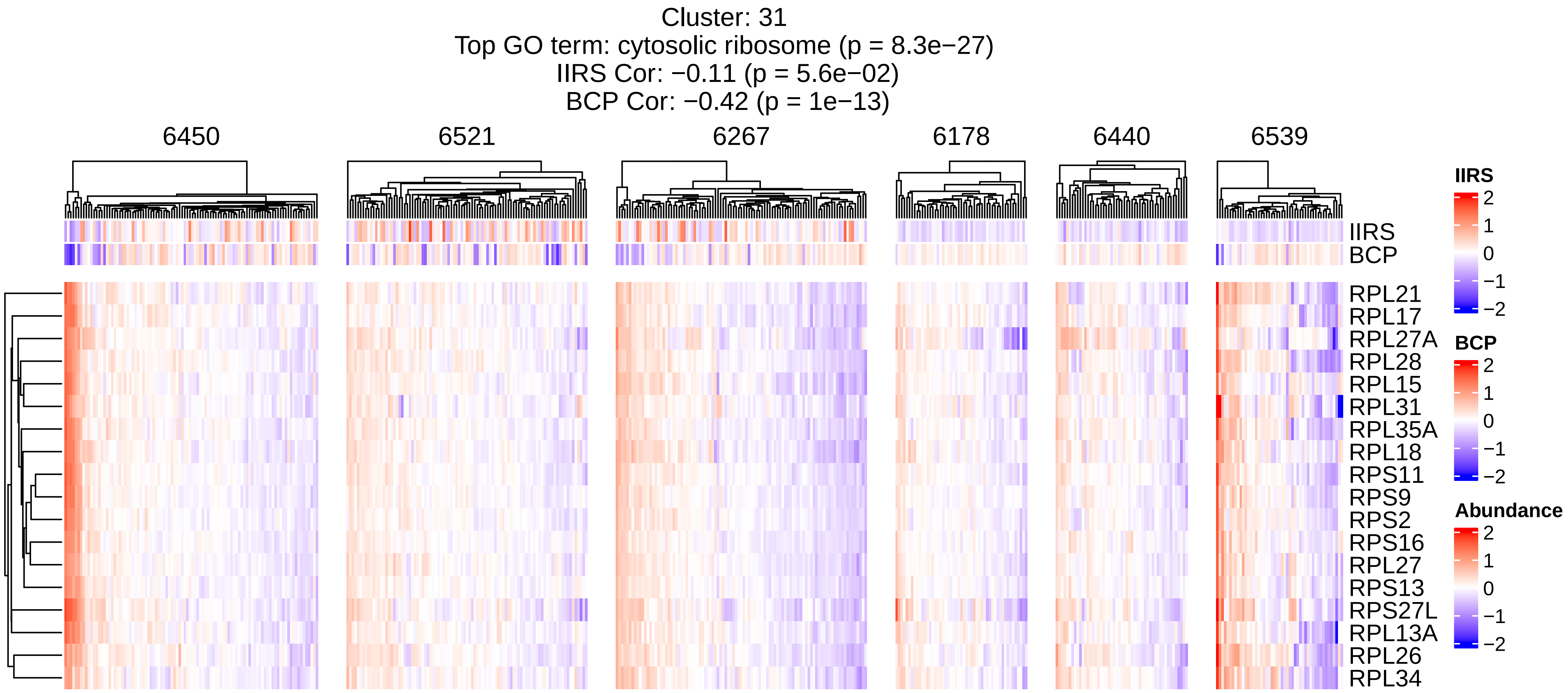
Cluster: 31
Top GO term: cytosolic ribosome (p = 8.3e−27)
IIRS Cor: −0.11 (p = 5.6e−02)
BCP Cor: −0.42 (p = 1e−13)
6450
6521
6267
6178
6440
6539
IIRS
BCP
RPL21
RPL17
RPL27A
RPL28
RPL15
RPL31
RPL35A
RPL18
RPS11
RPS9
RPS2
RPS16
RPL27
RPS13
RPS27L
RPL13A
RPL26
RPL34
IIRS
2
1
0
−1
−2
BCP
2
1
0
−1
−2
Abundance
2
1
0
−1
−2

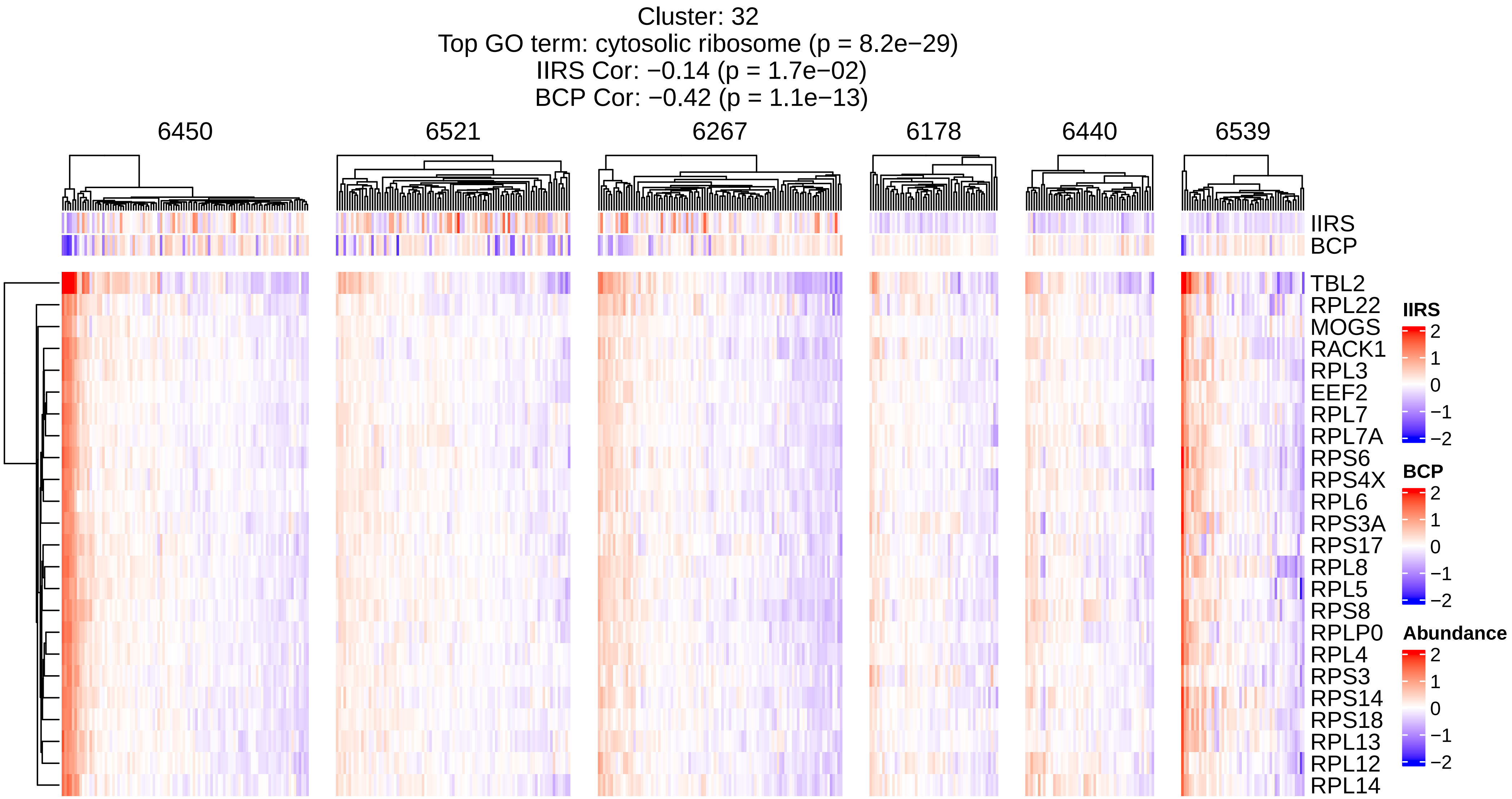

Cluster: 32
Top GO term: cytosolic ribosome (p = 8.2e−29)
IIRS Cor: −0.14 (p = 1.7e−02)
BCP Cor: −0.42 (p = 1.1e−13)
6450
6521
6267
6178
6440
6539
IIRS
BCP
TBL2
RPL22
MOGS
RACK1
RPL3
EEF2
RPL7
RPL7A
RPS6
RPS4X
RPL6
RPS3A
RPS17
RPL8
RPL5
RPS8
RPLP0
RPL4
RPS3
RPS14
RPS18
RPL13
RPL12
RPL14
IIRS
2
1
0
−1
−2
BCP
2
1
0
−1
−2
Abundance
2
1
0
−1
−2

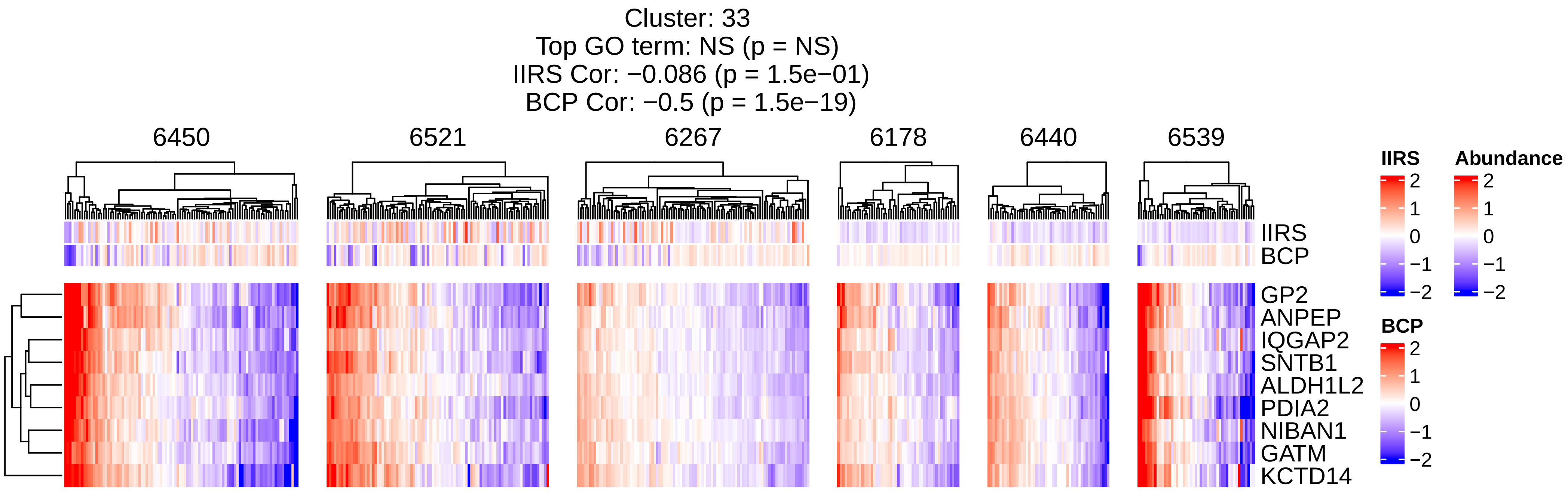
Cluster: 33
Top GO term: NS (p = NS)
IIRS Cor: −0.086 (p = 1.5e−01)
BCP Cor: −0.5 (p = 1.5e−19)
6450
6521
6267
6178
6440
6539
IIRS
BCP
GP2
ANPEP
IQGAP2
SNTB1
ALDH1L2
PDIA2
NIBAN1
GATM
KCTD14
IIRS
2
1
0
−1
−2
Abundance
2
1
0
−1
−2
BCP
2
1
0
−1
−2

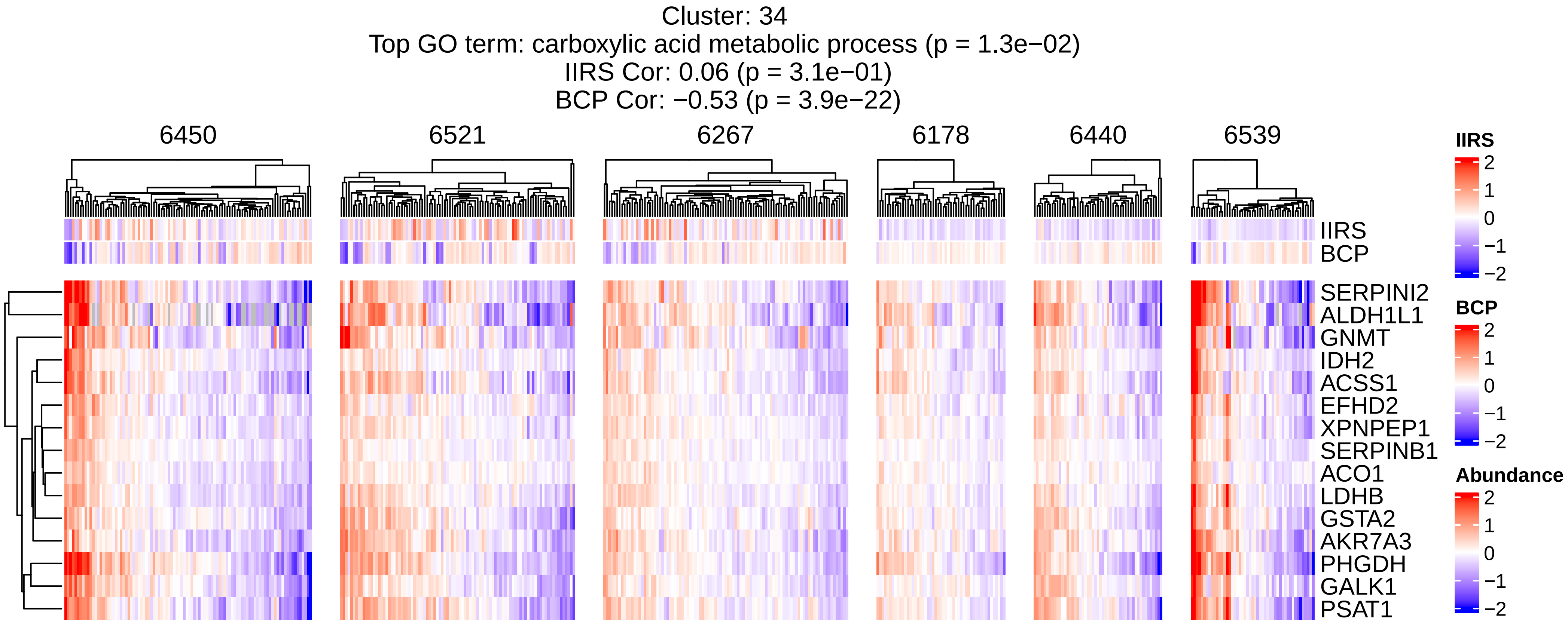
Cluster: 34
Top GO term: carboxylic acid metabolic process (p = 1.3e−02)
IIRS Cor: 0.06 (p = 3.1e−01)
BCP Cor: −0.53 (p = 3.9e−22)
6450
6521
6267
6178
6440
6539
IIRS
BCP
SERPINI2
ALDH1L1
GNMT
IDH2
ACSS1
EFHD2
XPNPEP1
SERPINB1
ACO1
LDHB
GSTA2
AKR7A3
PHGDH
GALK1
PSAT1
IIRS
2
1
0
−1
−2
BCP
2
1
0
−1
−2
Abundance
2
1
0
−1
−2

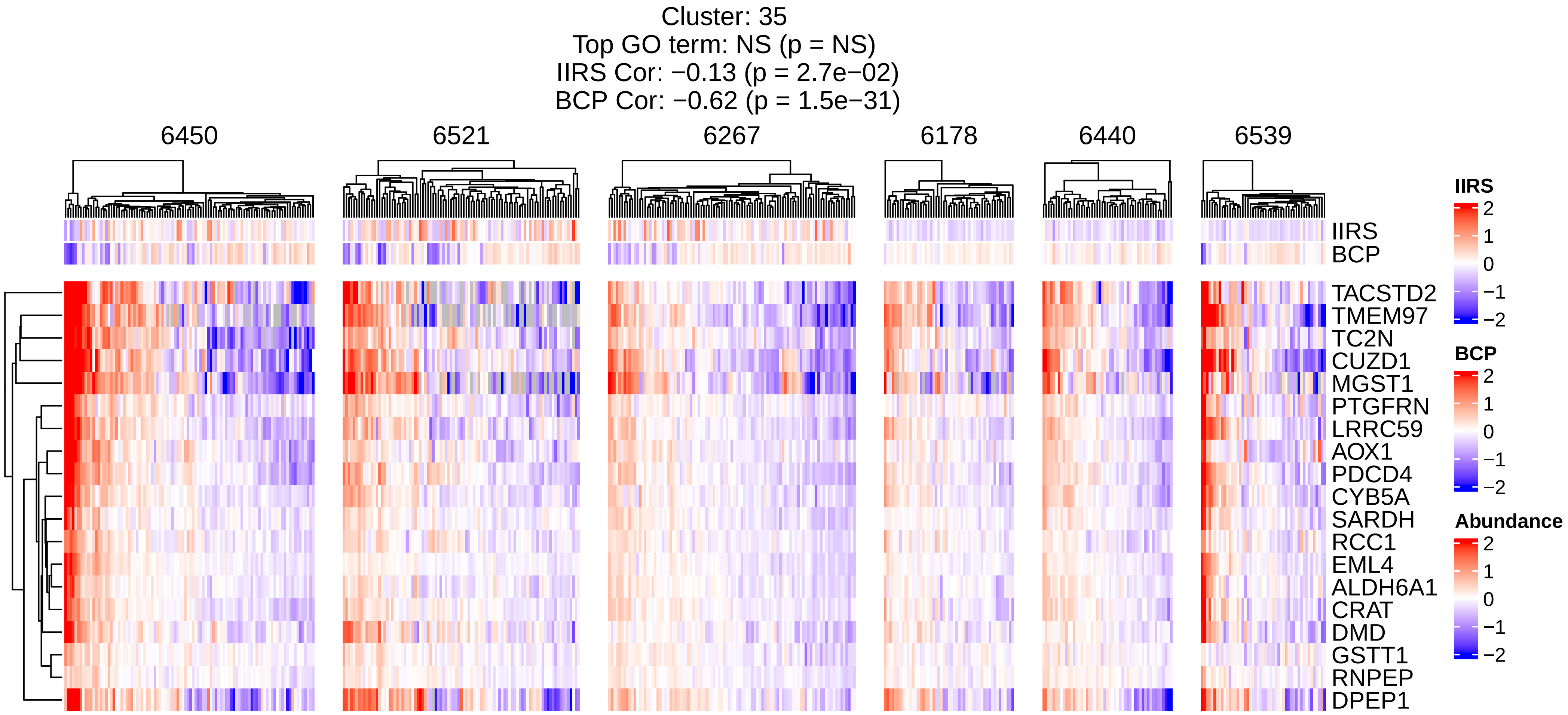
Cluster: 35
Top GO term: NS (p = NS)
IIRS Cor: −0.13 (p = 2.7e−02)
BCP Cor: −0.62 (p = 1.5e−31)
6450
6521
6267
6178
6440
6539
IIRS
BCP
TACSTD2
TMEM97
TC2N
CUZD1
MGST1
PTGFRN
LRRC59
AOX1
PDCD4
CYB5A
SARDH
RCC1
EML4
ALDH6A1
CRAT
DMD
GSTT1
RNPEP
DPEP1
IIRS
2
1
0
−1
−2
BCP
2
1
0
−1
−2
Abundance
2
1
0
−1
−2

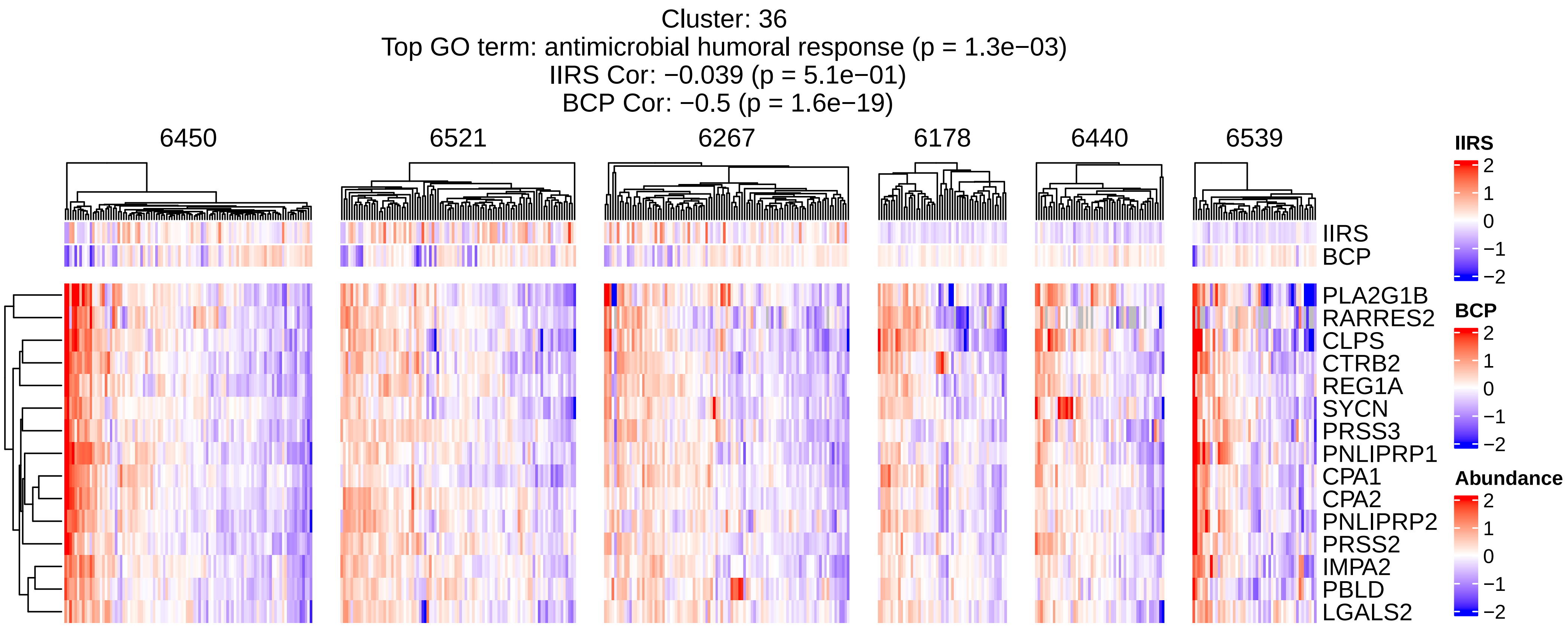

Cluster: 36
Top GO term: antimicrobial humoral response (p = 1.3e−03)
IIRS Cor: −0.039 (p = 5.1e−01)
BCP Cor: −0.5 (p = 1.6e−19)
6450
6521
6267
6178
6440
6539
IIRS
BCP
PLA2G1B
RARRES2
CLPS
CTRB2
REG1A
SYCN
PRSS3
PNLIPRP1
CPA1
CPA2
PNLIPRP2
PRSS2
IMPA2
PBLD
LGALS2
IIRS
2
1
0
−1
−2
BCP
2
1
0
−1
−2
Abundance
2
1
0
−1
−2

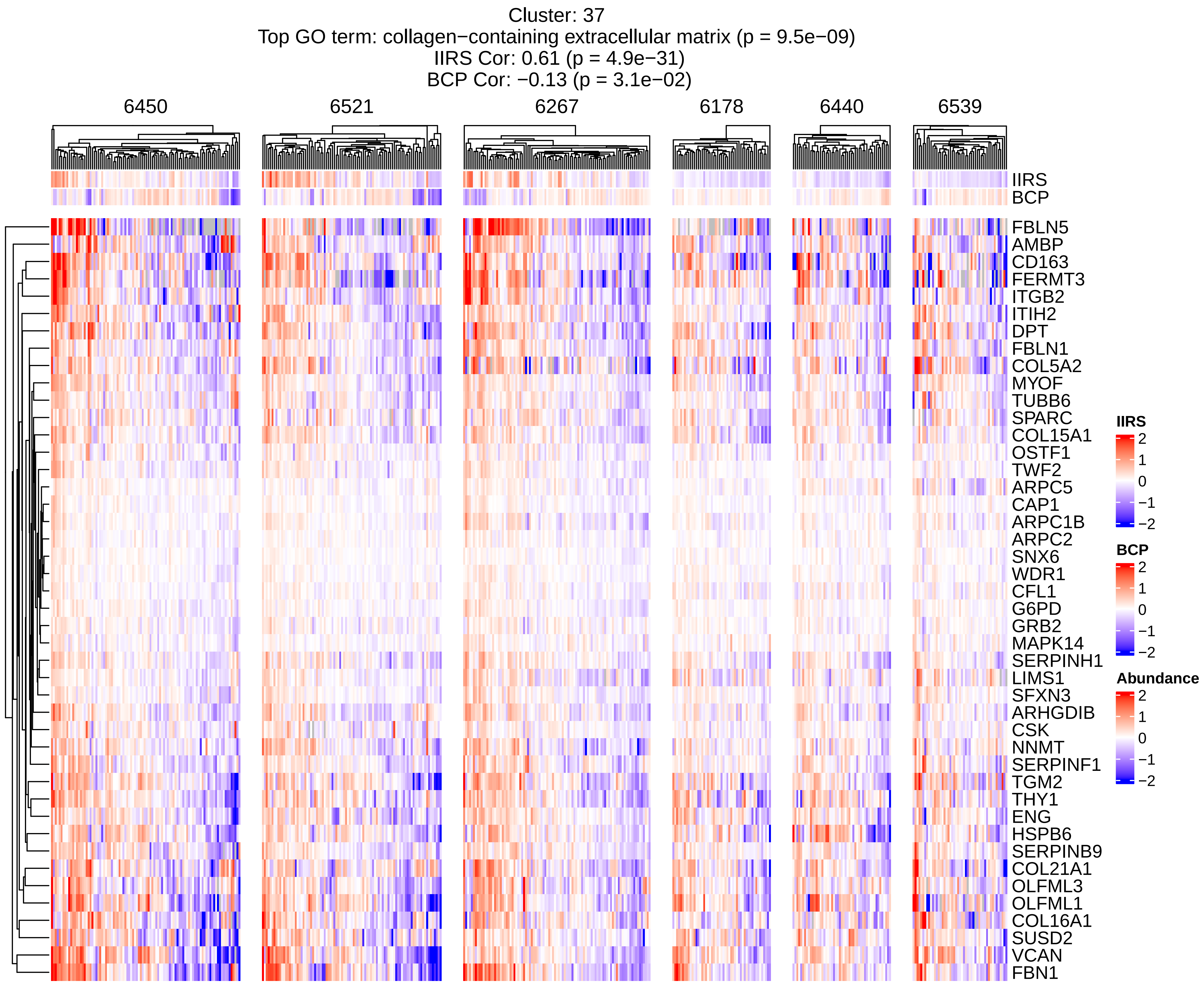
Cluster: 37
Top GO term: collagen−containing extracellular matrix (p = 9.5e−09)
IIRS Cor: 0.61 (p = 4.9e−31)
BCP Cor: −0.13 (p = 3.1e−02)
6450
6521
6267
6178
6440
6539
IIRS
BCP
FBLN5
AMBP
CD163
FERMT3
ITGB2
ITIH2
DPT
FBLN1
COL5A2
MYOF
TUBB6
SPARC
COL15A1
OSTF1
TWF2
ARPC5
CAP1
ARPC1B
ARPC2
SNX6
WDR1
CFL1
G6PD
GRB2
MAPK14
SERPINH1
LIMS1
SFXN3
ARHGDIB
CSK
NNMT
SERPINF1
TGM2
THY1
ENG
HSPB6
SERPINB9
COL21A1
OLFML3
OLFML1
COL16A1
SUSD2
VCAN
FBN1
IIRS
2
1
0
−1
−2
BCP
2
1
0
−1
−2
Abundance
2
1
0
−1
−2

Cluster: 38
Top GO term: neutrophil aggregation,autocrine signaling,calprotectin complex,Toll−like receptor 4 binding (p = 4.9e−03)
IIRS Cor: 0.034 (p = 5.6e−01)
BCP Cor: −0.34 (p = 2e−09)

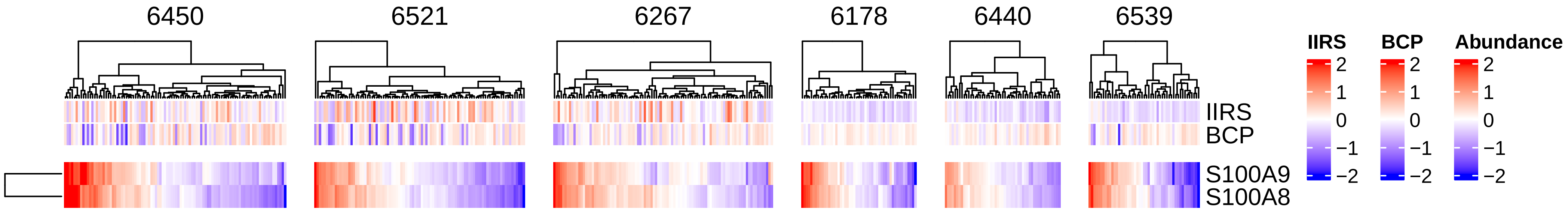

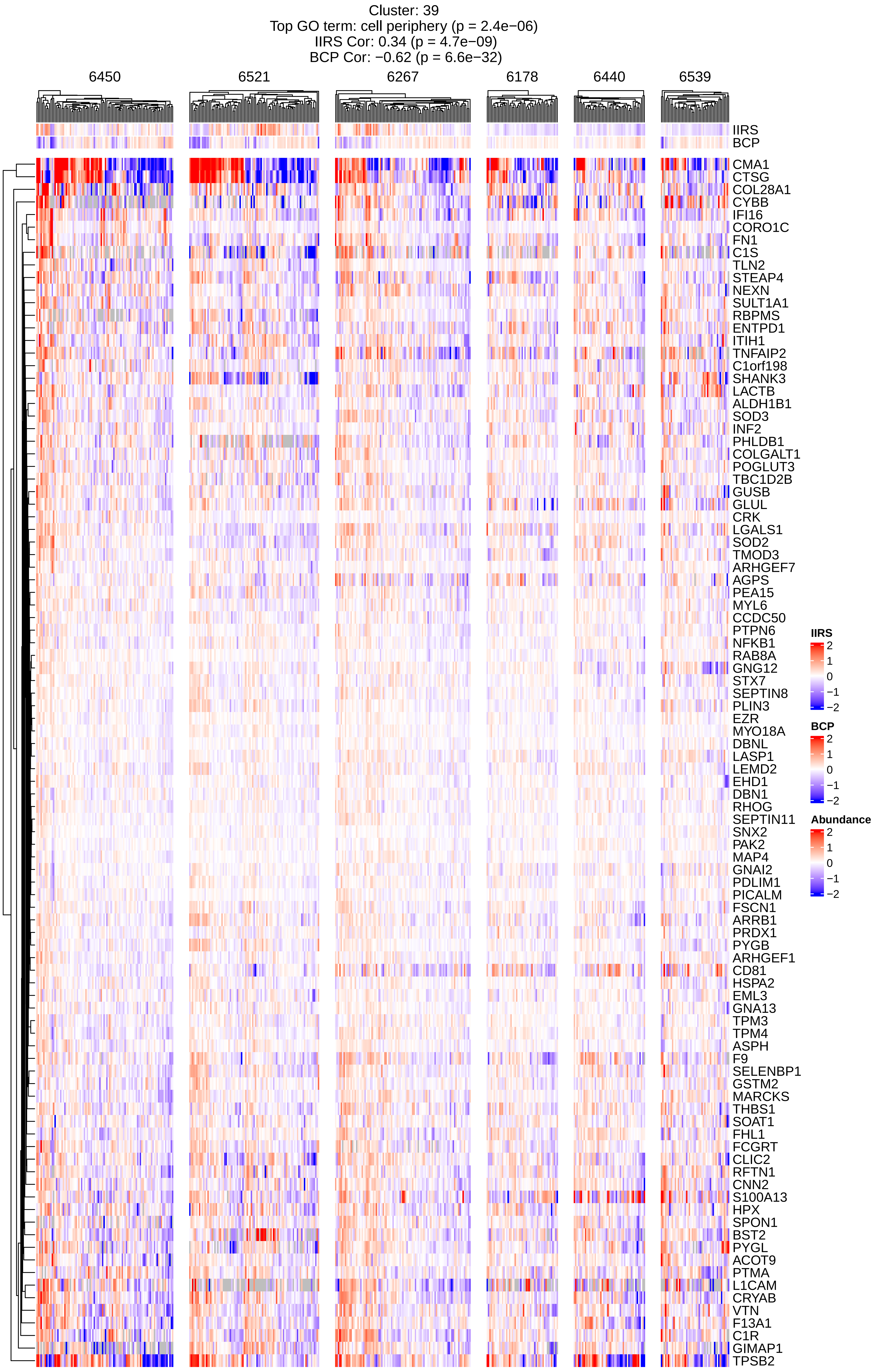
Cluster: 39
Top GO term: cell periphery (p = 2.4e−06)
IIRS Cor: 0.34 (p = 4.7e−09)
BCP Cor: −0.62 (p = 6.6e−32)
6450
6521
6267
6178
6440
6539
IIRS
BCP
CMA1
CTSG
COL28A1
CYBB
IFI16
CORO1C
FN1
C1S
TLN2
STEAP4
NEXN
SULT1A1
RBPMS
ENTPD1
ITIH1
TNFAIP2
C1orf198
SHANK3
LACTB
ALDH1B1
SOD3
INF2
PHLDB1
COLGALT1
POGLUT3
TBC1D2B
GUSB
GLUL
CRK
LGALS1
SOD2
TMOD3
ARHGEF7
AGPS
PEA15
MYL6
CCDC50
PTPN6
NFKB1
RAB8A
GNG12
STX7
SEPTIN8
PLIN3
EZR
MYO18A
DBNL
LASP1
LEMD2
EHD1
DBN1
RHOG
SEPTIN11
SNX2
PAK2
MAP4
GNAI2
PDLIM1
PICALM
FSCN1
ARRB1
PRDX1
PYGB
ARHGEF1
CD81
HSPA2
EML3
GNA13
TPM3
TPM4
ASPH
F9
SELENBP1
GSTM2
MARCKS
THBS1
SOAT1
FHL1
FCGRT
CLIC2
RFTN1
CNN2
S100A13
HPX
SPON1
BST2
PYGL
ACOT9
PTMA
L1CAM
CRYAB
VTN
F13A1
C1R
GIMAP1
TPSB2
IIRS
2
1
0
−1
−2
BCP
2
1
0
−1
−2
Abundance
2
1
0
−1
−2

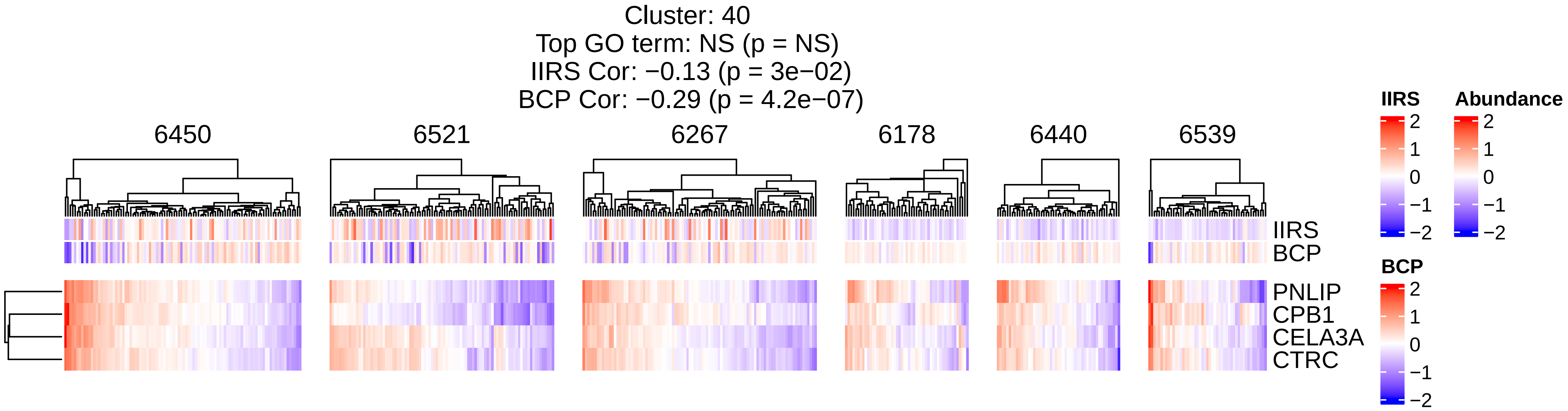

Cluster: 40
Top GO term: NS (p = NS)
IIRS Cor: −0.13 (p = 3e−02)
BCP Cor: −0.29 (p = 4.2e−07)
6450
6521
6267
6178
6440
6539
IIRS
BCP
PNLIP
CPB1
CELA3A
CTRC
IIRS
2
1
0
−1
−2
Abundance
2
1
0
−1
−2
BCP
2
1
0
−1
−2

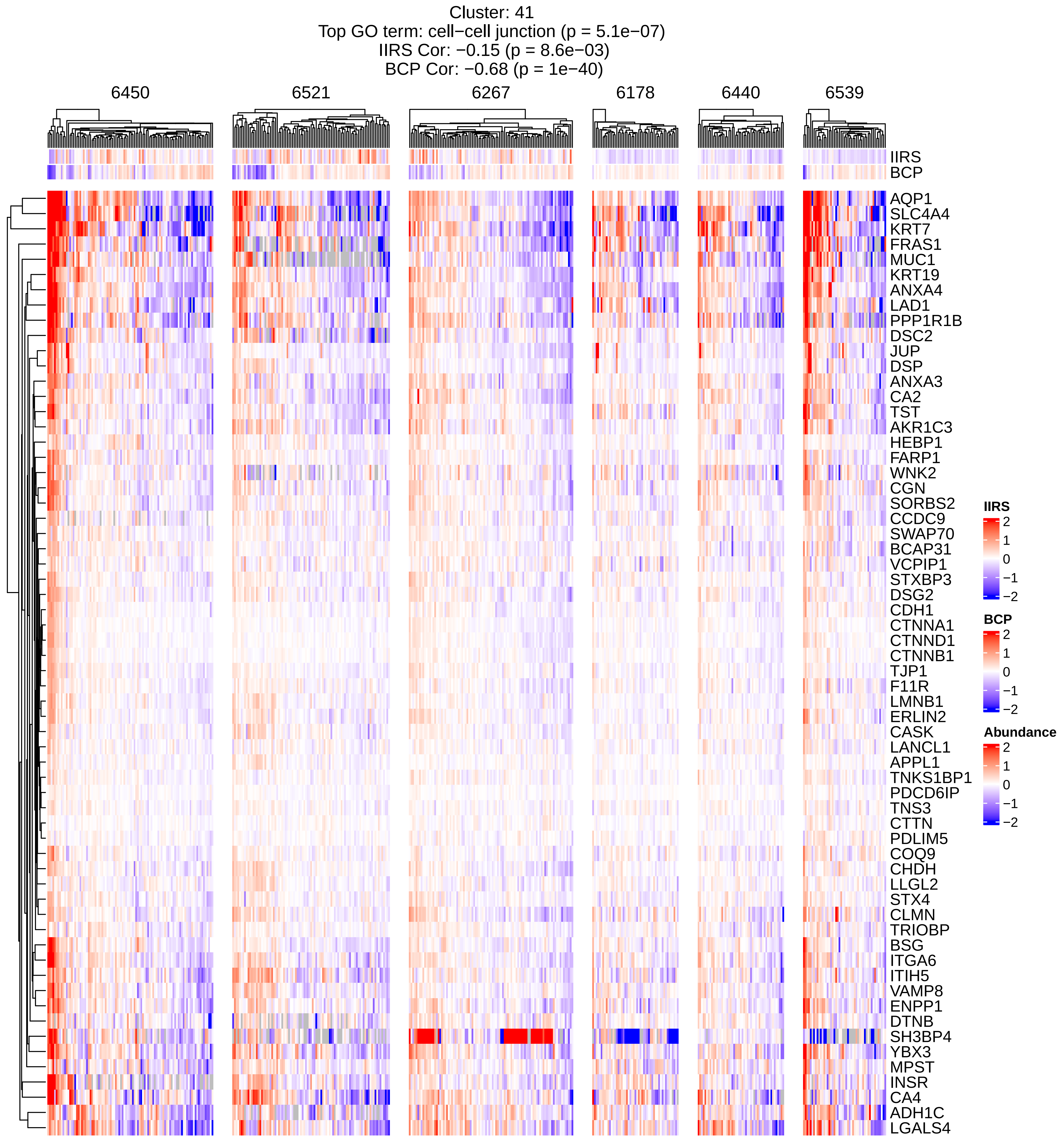
Cluster: 41
Top GO term: cell−cell junction (p = 5.1e−07)
IIRS Cor: −0.15 (p = 8.6e−03)
BCP Cor: −0.68 (p = 1e−40)
6450
6521
6267
6178
6440
6539
IIRS
BCP
AQP1
SLC4A4
KRT7
FRAS1
MUC1
KRT19
ANXA4
LAD1
PPP1R1B
DSC2
JUP
DSP
ANXA3
CA2
TST
AKR1C3
HEBP1
FARP1
WNK2
CGN
SORBS2
CCDC9
SWAP70
BCAP31
VCPIP1
STXBP3
DSG2
CDH1
CTNNA1
CTNND1
CTNNB1
TJP1
F11R
LMNB1
ERLIN2
CASK
LANCL1
APPL1
TNKS1BP1
PDCD6IP
TNS3
CTTN
PDLIM5
COQ9
CHDH
LLGL2
STX4
CLMN
TRIOBP
BSG
ITGA6
ITIH5
VAMP8
ENPP1
DTNB
SH3BP4
YBX3
MPST
INSR
CA4
ADH1C
LGALS4
IIRS
2
1
0
−1
−2
BCP
2
1
0
−1
−2
Abundance
2
1
0
−1
−2

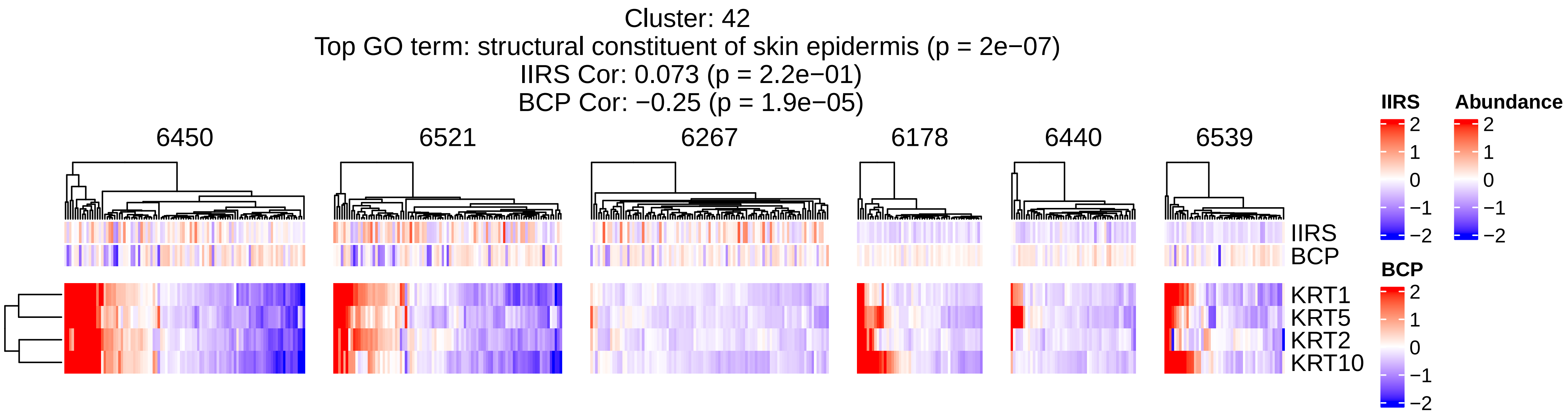

Cluster: 42
Top GO term: structural constituent of skin epidermis (p = 2e−07)
IIRS Cor: 0.073 (p = 2.2e−01)
BCP Cor: −0.25 (p = 1.9e−05)
6450
6521
6267
6178
6440
6539
IIRS
BCP
KRT1
KRT5
KRT2
KRT10
IIRS
2
1
0
−1
−2
Abundance
2
1
0
−1
−2
BCP
2
1
0
−1
−2

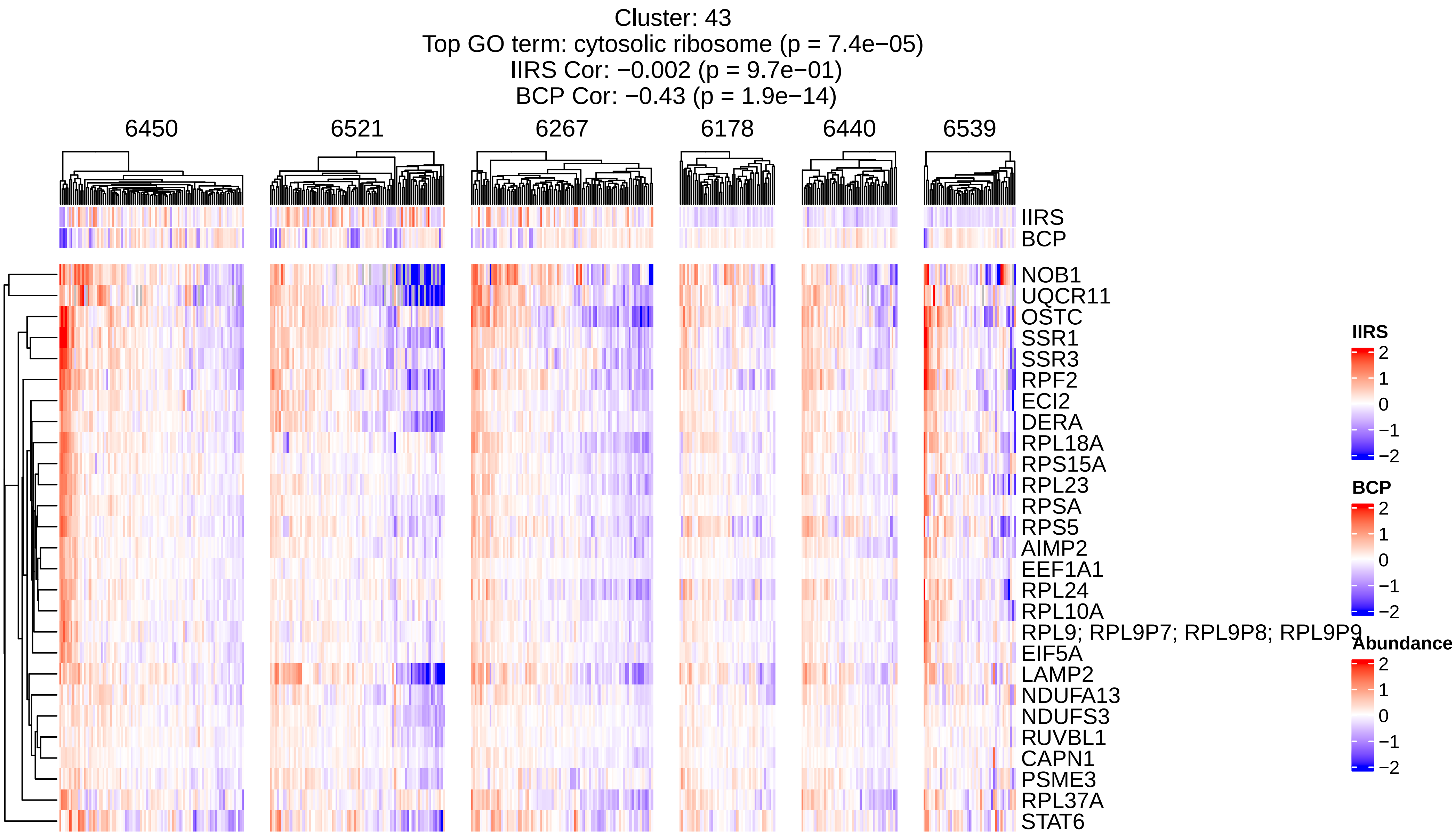
Cluster: 43
Top GO term: cytosolic ribosome (p = 7.4e−05)
IIRS Cor: −0.002 (p = 9.7e−01)
BCP Cor: −0.43 (p = 1.9e−14)
6450
6521
6267
6178
6440
6539
IIRS
BCP
NOB1
UQCR11
OSTC
SSR1
SSR3
RPF2
ECI2
DERA
RPL18A
RPS15A
RPL23
RPSA
RPS5
AIMP2
EEF1A1
RPL24
RPL10A
RPL9; RPL9P7; RPL9P8; RPL9P9
EIF5A
LAMP2
NDUFA13
NDUFS3
RUVBL1
CAPN1
PSME3
RPL37A
STAT6
IIRS
2
1
0
−1
−2
BCP
2
1
0
−1
−2
Abundance
2
1
0
−1
−2

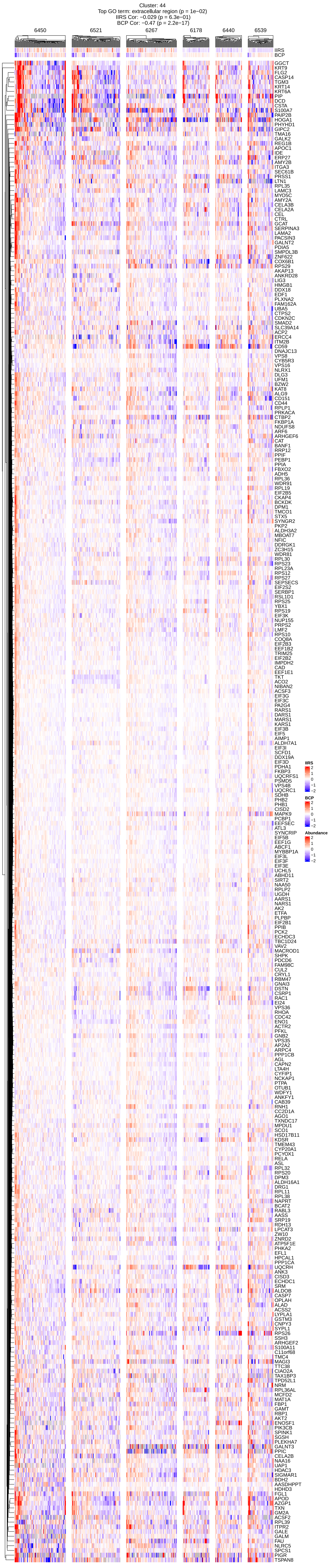
Cluster: 44
Top GO term: extracellular region (p = 1e−02)
IIRS Cor: −0.029 (p = 6.3e−01)
BCP Cor: −0.47 (p = 2.2e−17)
6450
6521
6267
6178
6440
6539
IIRS
BCP
GGCT
KRT9
FLG2
CASP14
TGM3
KRT14
KRT6A
PIP
DCD
CSTA
S100A7
PAIP2B
HOGA1
PHYHD1
GIPC2
TMA16
GALK2
REG1B
APOC1
IDE
ERP27
AMY2B
ITGA3
SEC61B
PRSS1
LTN1
RPL35
LAMC3
MYO5C
AMY2A
CELA3B
CELA2A
CEL
CTRL
GCAT
SERPINA3
LAMA2
PACSIN3
GALNT2
PDIA5
SMPDL3B
ZNF622
COX6B1
RPS29
AKAP13
ANKRD28
LIG3
HMGB1
DDX18
EDF1
PLXNA2
FAM162A
UBA5
CTPS2
CDKN2C
SMAD2
SLC39A14
ACP2
ERCC4
ITM2B
CD59
DNAJC13
VPS8
CYB5R3
VPS16
NLRX1
DLG3
UFM1
BZW2
KAT8
CD151
CD44
RPLP1
PRKACA
CTBP2
FKBP1A
NDUFS8
ARF6
ARHGEF6
CAT
BANF1
RRP12
PPIF
PEBP1
PPIA
FBXO2
ADH5
RPL36
WDR91
RPL19
EIF2B5
CKAP4
BCKDK
DPM1
TMCO1
STX5
SYNGR2
PKP2
ALDH3A2
MBOAT7
NFIC
DDRGK1
ZC3H15
WDR81
RPL30
RPS23
RPL23A
RPS12
RPS27
SEPSECS
EIF2S2
SERBP1
RSL1D1
RPS25
YBX1
RPS19
EIF3K
NUP155
PRPS2
LMF2
RPS10
COQ8A
EIF2B3
EEF1B2
TRIM25
EIF2B2
IMPDH2
CAD
EEF1E1
TKT
ACO2
NIBAN2
ACSF3
EIF3G
EIF3C
PA2G4
RARS1
DARS1
MARS1
KARS1
EIF3B
EIF5
AIMP1
ALDH7A1
EIF3I
SCFD1
DDX19A
EIF3D
PDHA1
FKBP3
UQCRFS1
PSMD5
VPS4B
UQCRC1
SDHB
PHB2
PHB1
CISD2
MAPK9
PCBP1
EEFSEC
ATL3
SYNCRIP
EIF5B
EEF1G
ABCF1
MYBBP1A
EIF3L
EIF3F
EIF3E
UCHL5
ABHD11
SIRT2
NAA50
RPLP2
UGDH
AARS1
NARS1
AK2
ETFA
PLPBP
EIF2B1
PPIB
PCK2
ECHDC3
TBC1D24
VAV2
MACROD1
SHPK
PDCD6
FAM98C
CUL2
CRYL1
RBM47
GNAI3
DSTN
CSRP1
RAC1
EI24
VPS36
RHOA
CDC42
ENO1
ACTR2
PFKL
GNB2
VPS35
AP2A2
ARPC4
PPP1CB
AGL
CAPN2
LTA4H
CYFIP1
NCKAP1
PTPA
OTUB1
WDFY1
ANKFY1
CAB39
RNH1
CC2D1A
AGO1
TXNDC17
MPDU1
SCO1
HSD17B11
KDSR
TMEM43
CYP20A1
PCYOX1
RELA
ASL
RPL32
RPS20
DPM3
ALDH16A1
DRG1
RPL11
RPL38
NAPRT
BCAT2
RABL3
AASS
SRP19
RDH13
LPCAT3
ZW10
ZNRD2
ATP5F1E
PHKA2
EFL1
HPCAL1
PPP1CA
UQCRH
ANK3
CISD3
ECHDC1
SRM
ALDOB
CASP7
OPLAH
ALAD
ACSS2
LYPLA1
GSTM3
CNPY3
SYPL1
RPS26
SSH3
ARHGEF2
S100A11
C11orf68
TMC4
MAGI3
TTC38
CIAO2A
TAX1BP3
TPD52L1
NRM
RPL36AL
MCFD2
MAT1A
FBP1
GAMT
RBP1
AKT2
ENOSF1
PIK3CB
SPINK1
SGSH
PLEKHA7
GALNT3
PPIC
CELA2B
NAA16
UAP1
HDAC3
SIGMAR1
BDH2
AASDHPPT
HDHD3
FGL1
APOD
AZGP1
TXN
GM2A
ACSF2
RPL39
ITPR2
GALE
GALM
FAU
NLRC5
SPCS2
PIGR
TSPAN8
IIRS
2
1
0
−1
−2
BCP
2
1
0
−1
−2
Abundance
2
1
0
−1
−2

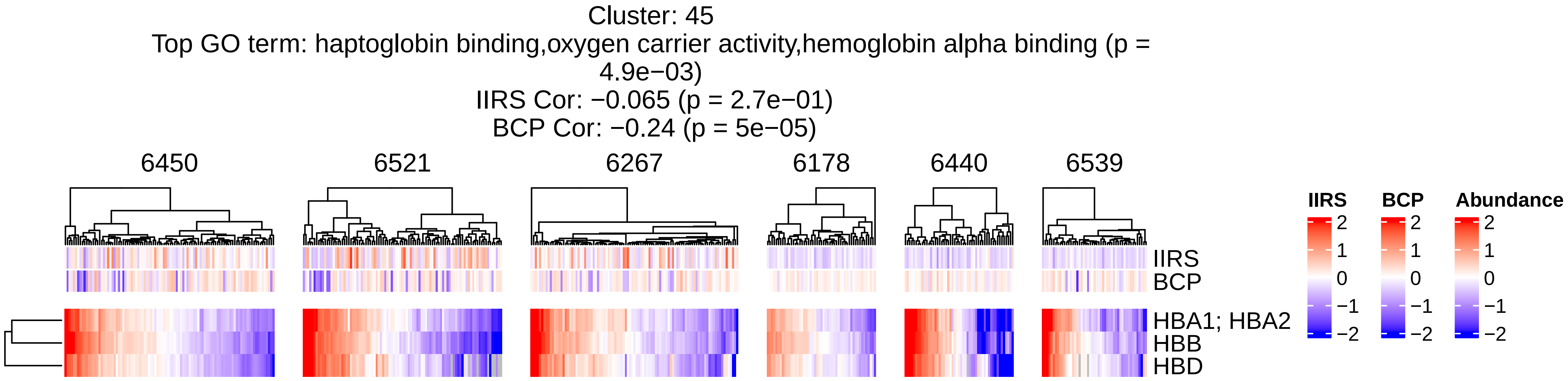
Cluster: 45
Top GO term: haptoglobin binding,oxygen carrier activity,hemoglobin alpha binding (p = 4.9e−03)
IIRS Cor: −0.065 (p = 2.7e−01)
BCP Cor: −0.24 (p = 5e−05)
6450
6521
6267
6178
6440
6539
IIRS
BCP
HBA1; HBA2
HBB
HBD
IIRS
2
1
0
−1
−2
BCP
2
1
0
−1
−2
Abundance
2
1
0
−1
−2

Cluster: 46
Top GO term: nucleoplasm (p = 1.3e−10)
IIRS Cor: −0.21 (p = 4.2e−04)
BCP Cor: −0.35 (p = 1.1e−09)

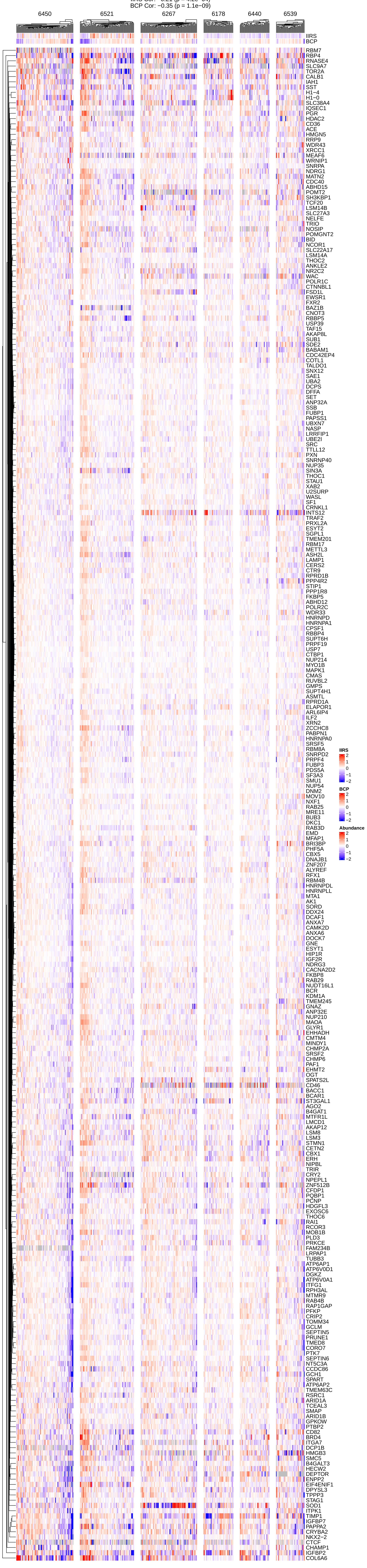

Cluster: 47
Top GO term: organelle membrane (p = 9.5e−03)
IIRS Cor: 0.12 (p = 5e−02)
BCP Cor: 0.25 (p = 2.1e−05)

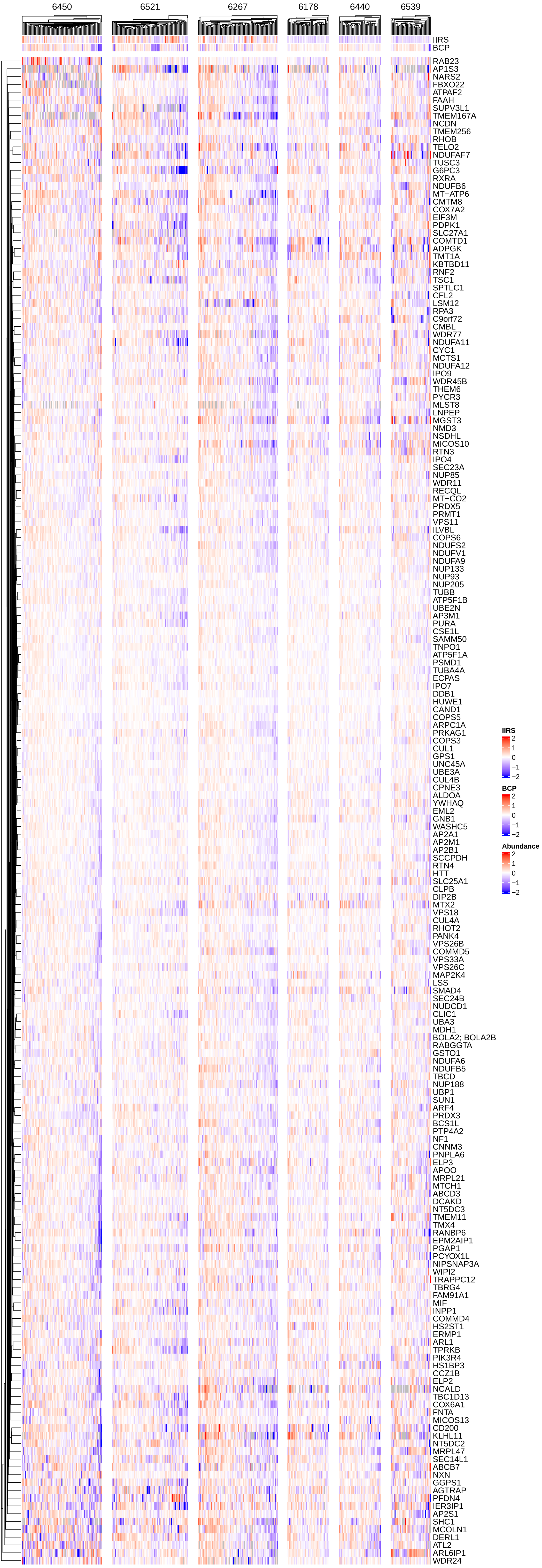

Cluster: 48A
Top GO term: nucleus (p = 7.2e−03)
IIRS Cor: 0.0096 (p = 8.7e−01)
BCP Cor: 0.067 (p = 2.6e−01)

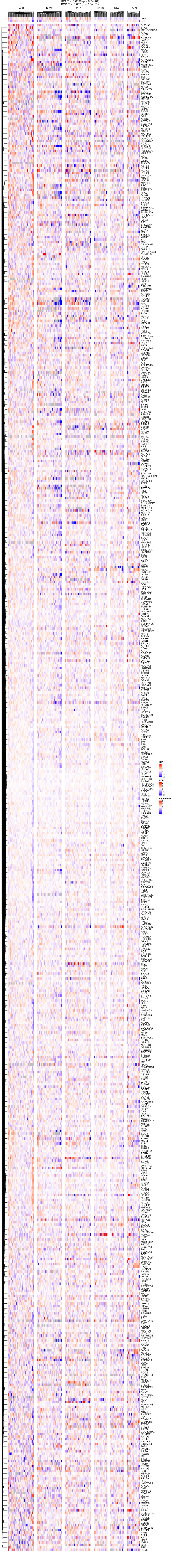

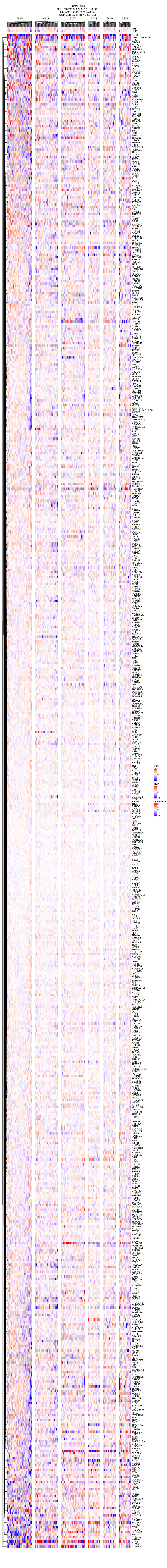
Cluster: 48B
Top GO term: nucleus (p = 7.2e−03)
IIRS Cor: 0.0096 (p = 8.7e−01)
BCP Cor: 0.067 (p = 2.6e−01)
6450
6521
6267
6178
6440
6539
IIRS
BCP

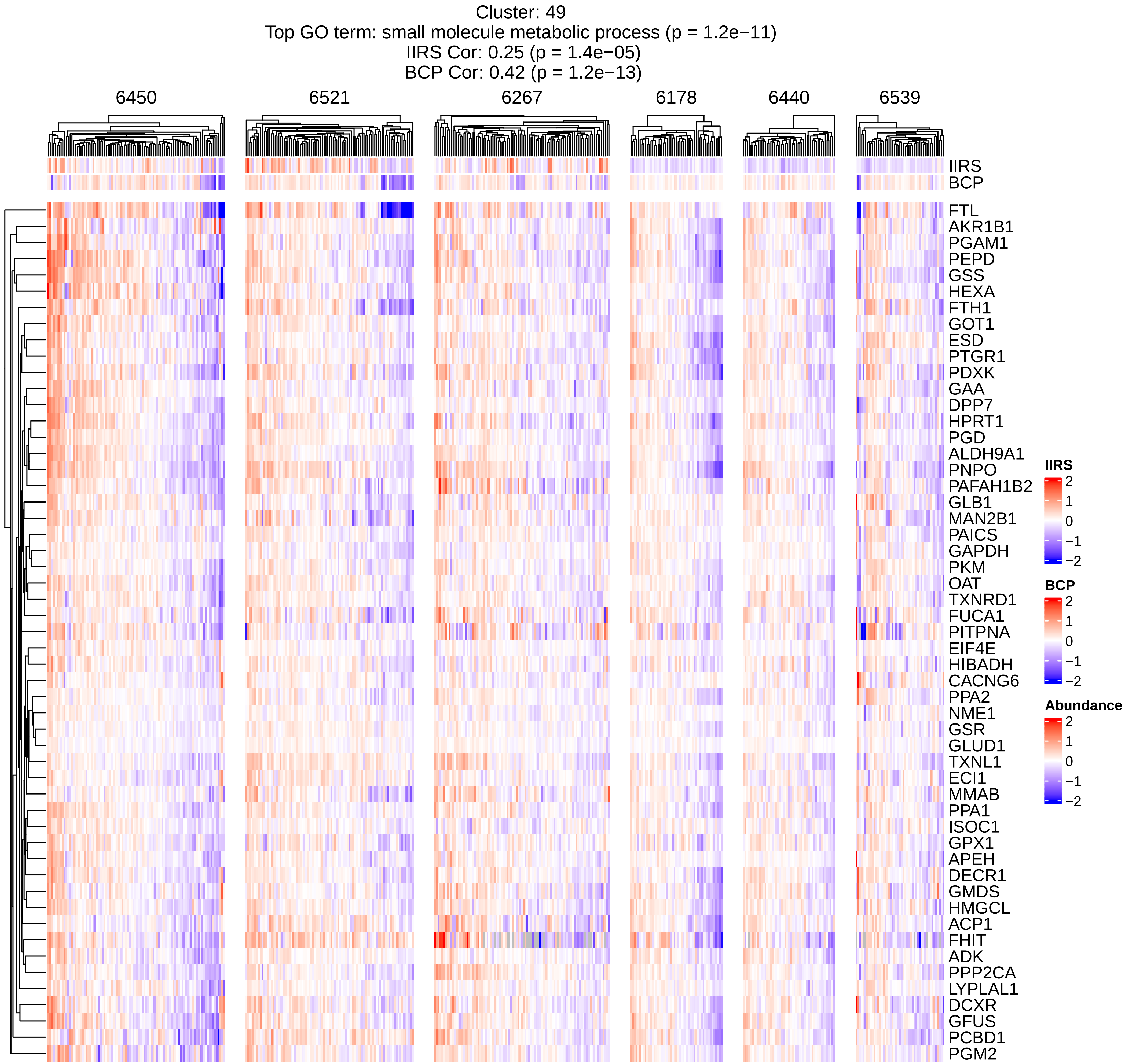
Cluster: 49
Top GO term: small molecule metabolic process (p = 1.2e−11)
IIRS Cor: 0.25 (p = 1.4e−05)
BCP Cor: 0.42 (p = 1.2e−13)
6450
6521
6267
6178
6440
6539
IIRS
BCP
FTL
AKR1B1
PGAM1
PEPD
GSS
HEXA
FTH1
GOT1
ESD
PTGR1
PDXK
GAA
DPP7
HPRT1
PGD
ALDH9A1
PNPO
PAFAH1B2
GLB1
MAN2B1
PAICS
GAPDH
PKM
OAT
TXNRD1
FUCA1
PITPNA
EIF4E
HIBADH
CACNG6
PPA2
NME1
GSR
GLUD1
TXNL1
ECI1
MMAB
PPA1
ISOC1
GPX1
APEH
DECR1
GMDS
HMGCL
ACP1
FHIT
ADK
PPP2CA
LYPLAL1
DCXR
GFUS
PCBD1
PGM2
IIRS
2
1
0
−1
−2
BCP
2
1
0
−1
−2
Abundance
2
1
0
−1
−2

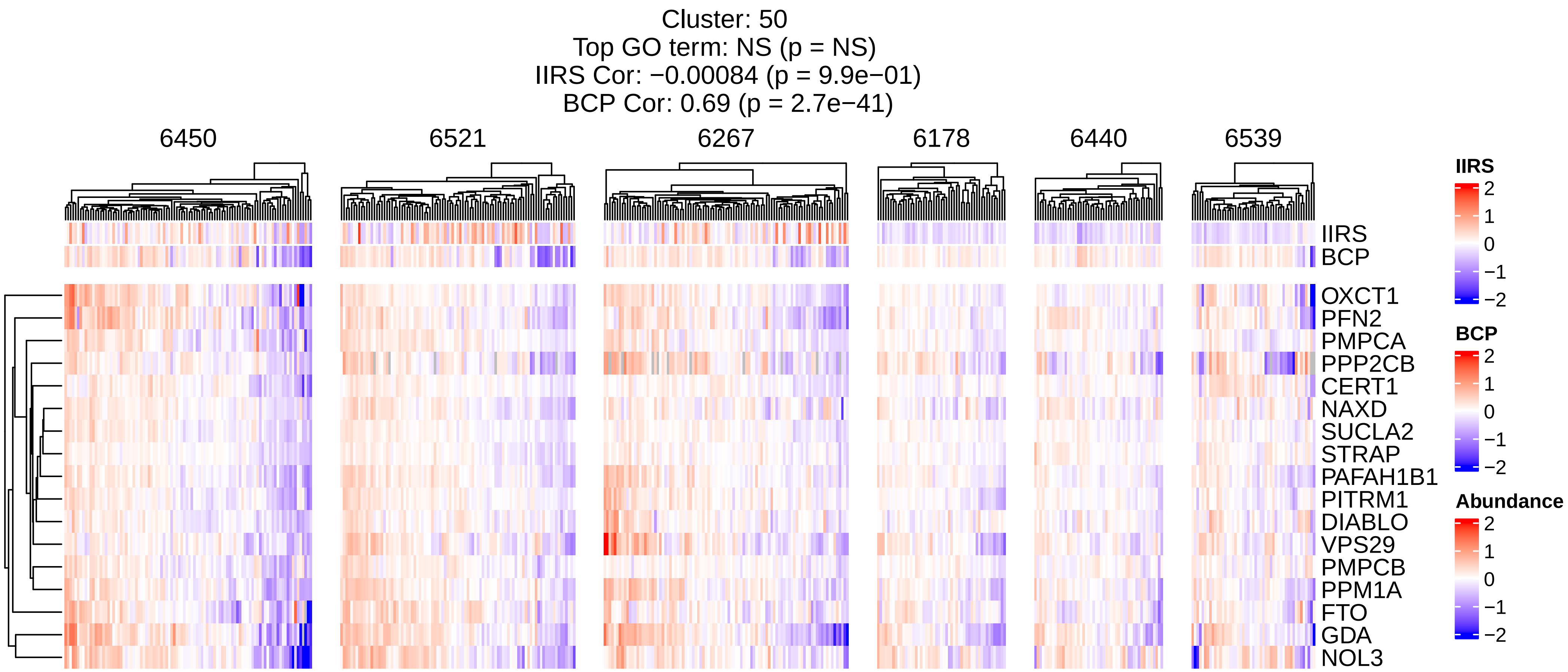
Cluster: 50
Top GO term: NS (p = NS)
IIRS Cor: −0.00084 (p = 9.9e−01)
BCP Cor: 0.69 (p = 2.7e−41)
6450
6521
6267
6178
6440
6539
IIRS
BCP
OXCT1
PFN2
PMPCA
PPP2CB
CERT1
NAXD
SUCLA2
STRAP
PAFAH1B1
PITRM1
DIABLO
VPS29
PMPCB
PPM1A
FTO
GDA
NOL3
IIRS
2
1
0
−1
−2
BCP
2
1
0
−1
−2
Abundance
2
1
0
−1
−2

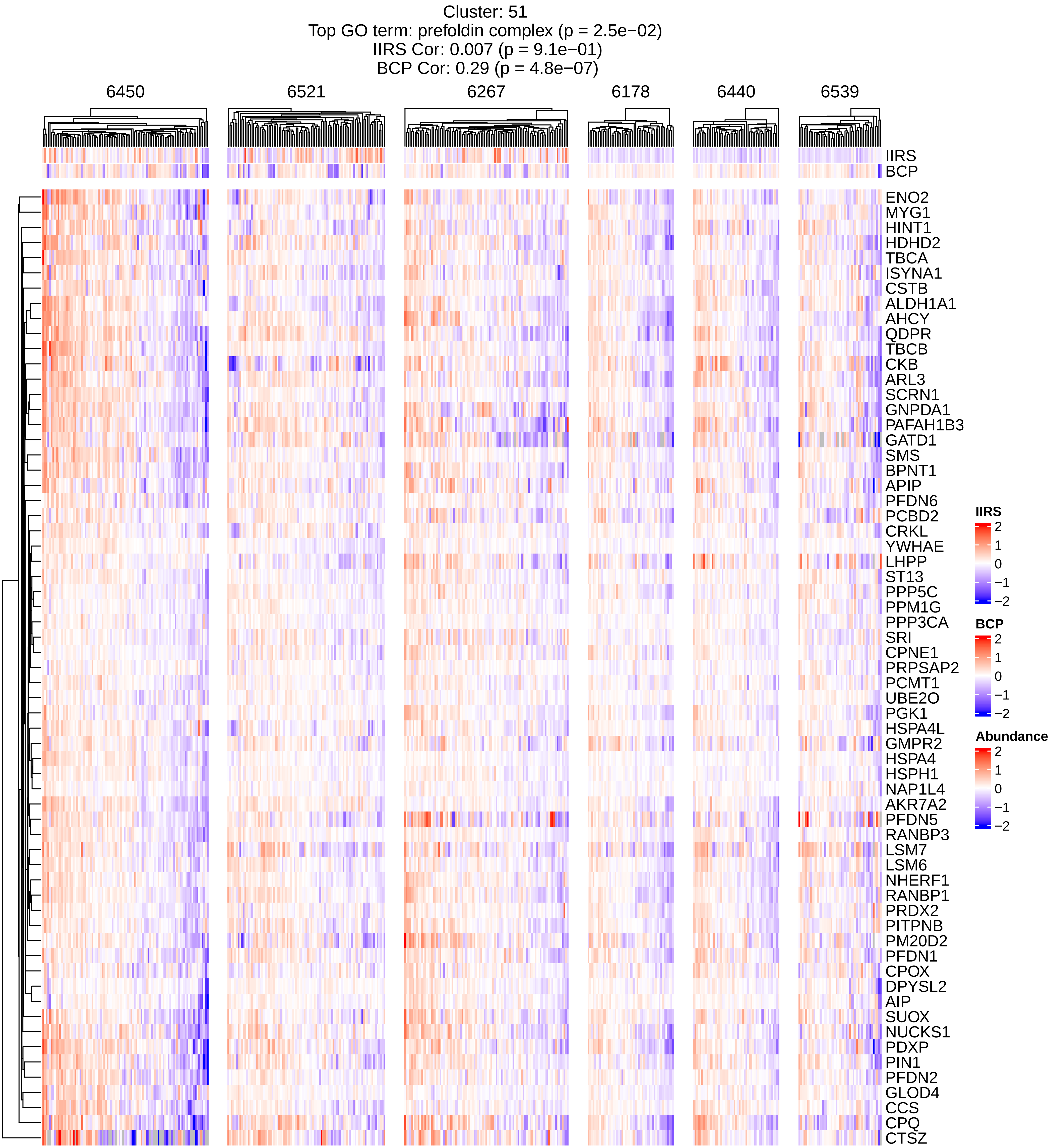

Cluster: 51
Top GO term: prefoldin complex (p = 2.5e−02)
IIRS Cor: 0.007 (p = 9.1e−01)
BCP Cor: 0.29 (p = 4.8e−07)
6450
6521
6267
6178
6440
6539
IIRS
BCP
ENO2
MYG1
HINT1
HDHD2
TBCA
ISYNA1
CSTB
ALDH1A1
AHCY
QDPR
TBCB
CKB
ARL3
SCRN1
GNPDA1
PAFAH1B3
GATD1
SMS
BPNT1
APIP
PFDN6
PCBD2
CRKL
YWHAE
LHPP
ST13
PPP5C
PPM1G
PPP3CA
SRI
CPNE1
PRPSAP2
PCMT1
UBE2O
PGK1
HSPA4L
GMPR2
HSPA4
HSPH1
NAP1L4
AKR7A2
PFDN5
RANBP3
LSM7
LSM6
NHERF1
RANBP1
PRDX2
PITPNB
PM20D2
PFDN1
CPOX
DPYSL2
AIP
SUOX
NUCKS1
PDXP
PIN1
PFDN2
GLOD4
CCS
CPQ
CTSZ
IIRS
2
1
0
−1
−2
BCP
2
1
0
−1
−2
Abundance
2
1
0
−1
−2

Cluster: 52
Top GO term: acyl−CoA dehydrogenase activity,oxidoreductase activity, acting on the CH−CH group of donors, with a flavin as acceptor (p = 3.2e−02)
IIRS Cor: 0.15 (p = 1.1e−02)
BCP Cor: 0.28 (p = 1.5e−06)

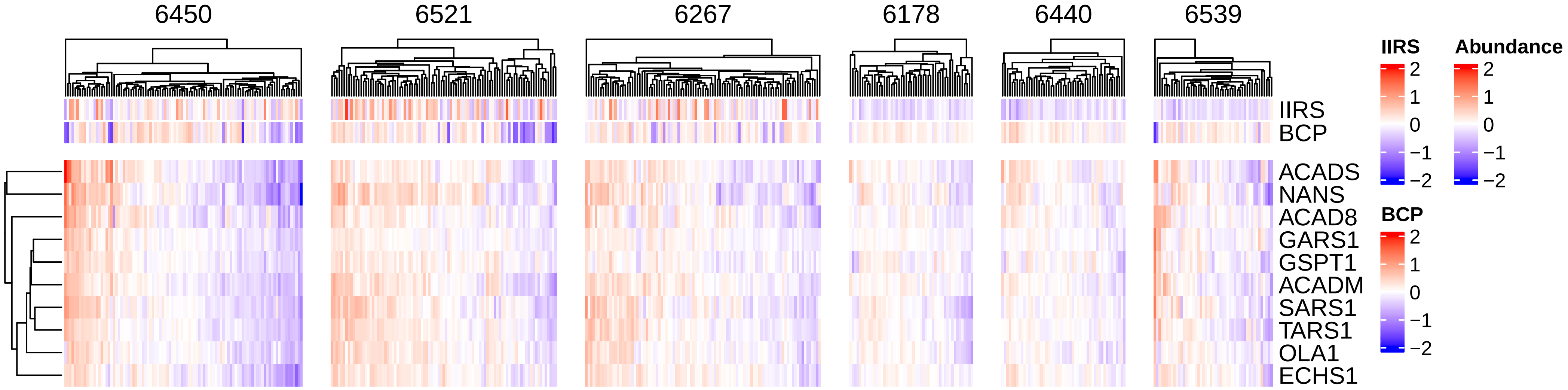

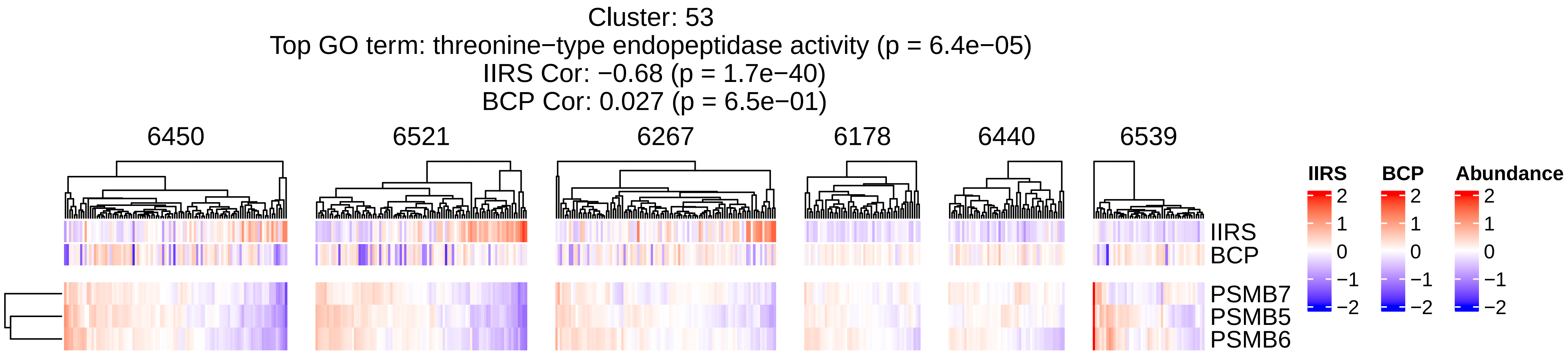
Cluster: 53
Top GO term: threonine−type endopeptidase activity (p = 6.4e−05)
IIRS Cor: −0.68 (p = 1.7e−40)
BCP Cor: 0.027 (p = 6.5e−01)
6450
6521
6267
6178
6440
6539
IIRS
BCP
PSMB7
PSMB5
PSMB6
IIRS
BCP
Abundance
2
1
0
−1
−2

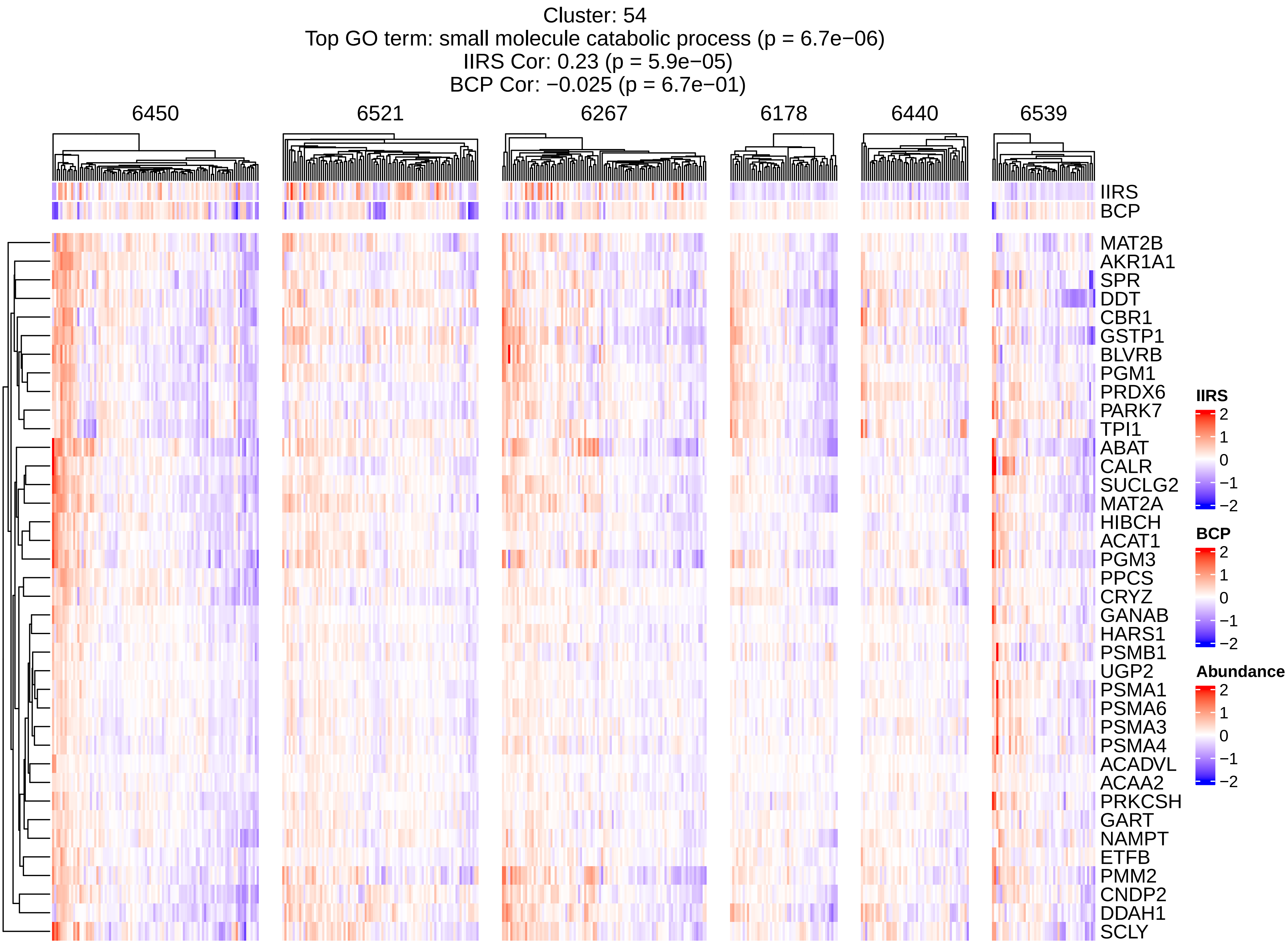
Cluster: 54
Top GO term: small molecule catabolic process (p = 6.7e−06)
IIRS Cor: 0.23 (p = 5.9e−05)
BCP Cor: −0.025 (p = 6.7e−01)
6450
6521
6267
6178
6440
6539
IIRS
BCP
MAT2B
AKR1A1
SPR
DDT
CBR1
GSTP1
BLVRB
PGM1
PRDX6
PARK7
TPI1
ABAT
CALR
SUCLG2
MAT2A
HIBCH
ACAT1
PGM3
PPCS
CRYZ
GANAB
HARS1
PSMB1
UGP2
PSMA1
PSMA6
PSMA3
PSMA4
ACADVL
ACAA2
PRKCSH
GART
NAMPT
ETFB
PMM2
CNDP2
DDAH1
SCLY
IIRS
2
1
0
−1
−2
BCP
2
1
0
−1
−2
Abundance
2
1
0
−1
−2

Cluster: 55
Top GO term: mitochondrial matrix (p = 1.4e−02)
IIRS Cor: 0.16 (p = 8.3e−03)
BCP Cor: 0.12 (p = 4.2e−02)

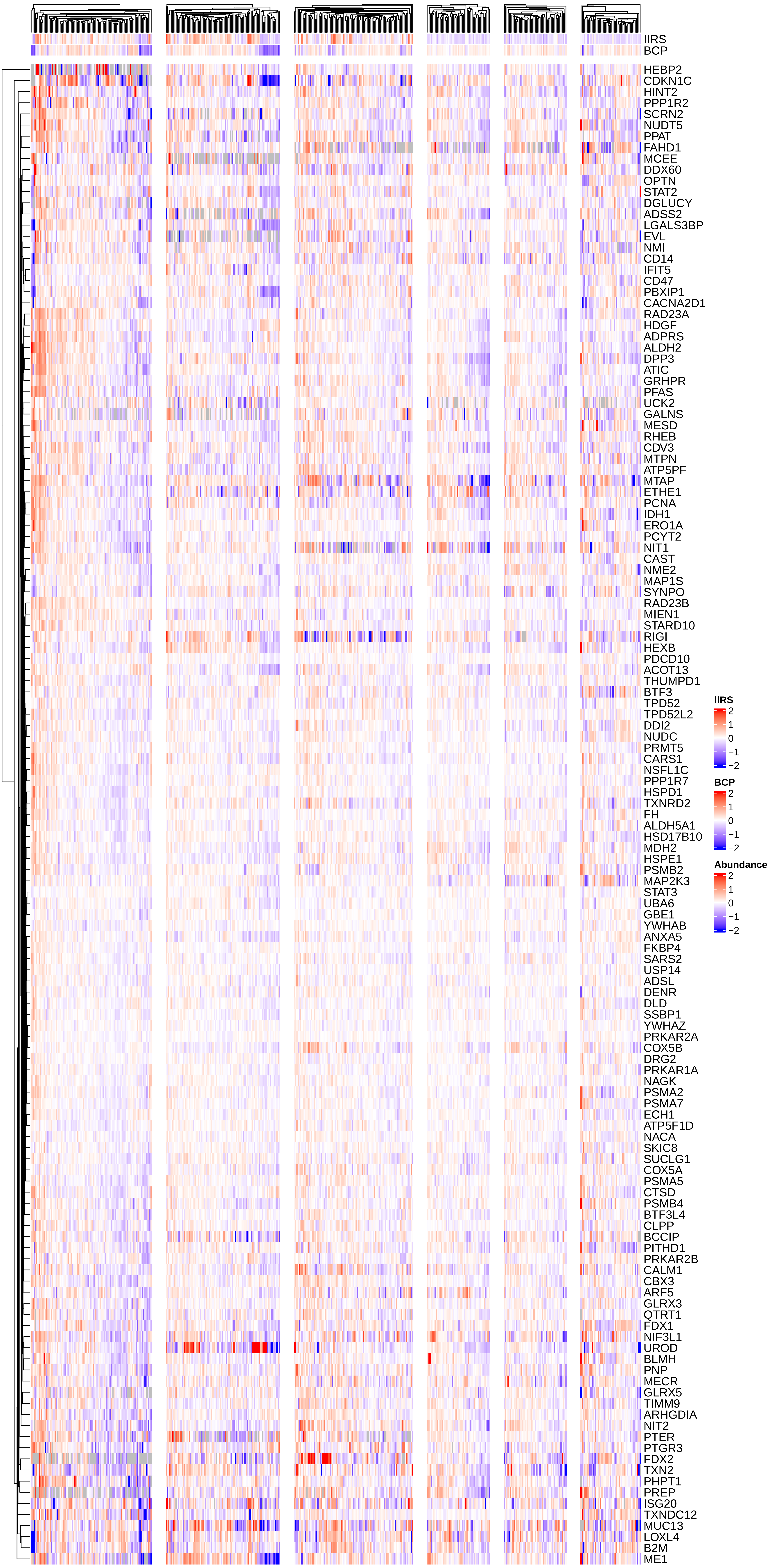

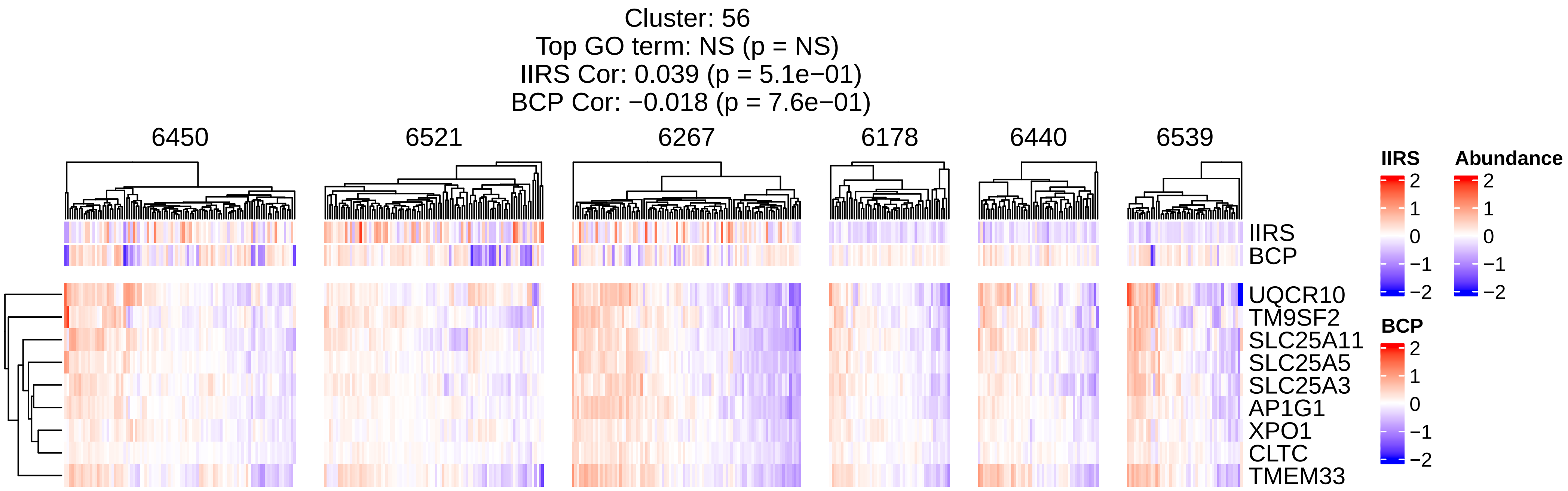
Cluster: 56
Top GO term: NS (p = NS)
IIRS Cor: 0.039 (p = 5.1e−01)
BCP Cor: −0.018 (p = 7.6e−01)
6450
6521
6267
6178
6440
6539
IIRS
BCP
UQCR10
TM9SF2
SLC25A11
SLC25A5
SLC25A3
AP1G1
XPO1
CLTC
TMEM33
IIRS
Abundance
BCP
2
1
0
−1
−2

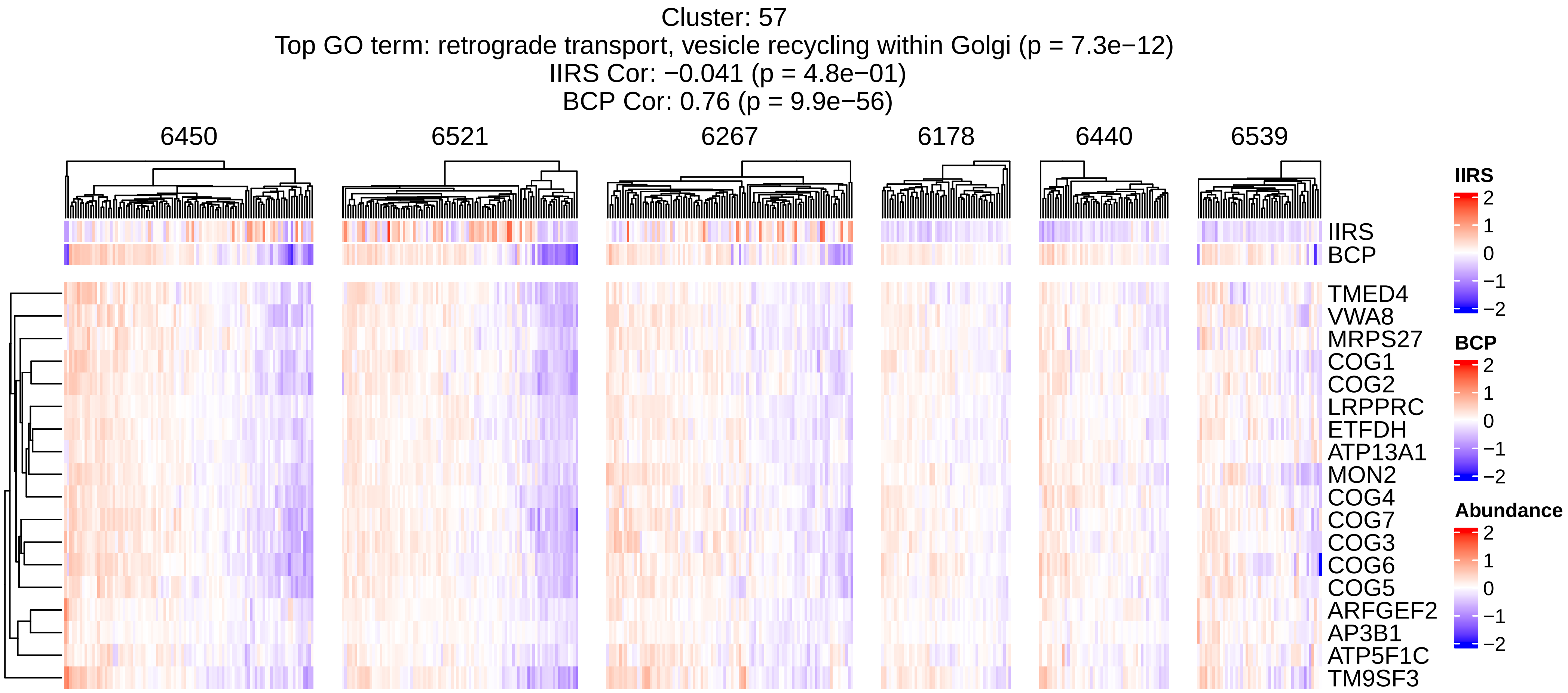
Cluster: 57
Top GO term: retrograde transport, vesicle recycling within Golgi (p = 7.3e−12)
IIRS Cor: −0.041 (p = 4.8e−01)
BCP Cor: 0.76 (p = 9.9e−56)
6450
6521
6267
6178
6440
6539
IIRS
BCP
TMED4
VWA8
MRPS27
COG1
COG2
LRPPRC
ETFDH
ATP13A1
MON2
COG4
COG7
COG3
COG6
COG5
ARFGEF2
AP3B1
ATP5F1C
TM9SF3
IIRS
2
1
0
−1
−2
BCP
2
1
0
−1
−2
Abundance
2
1
0
−1
−2

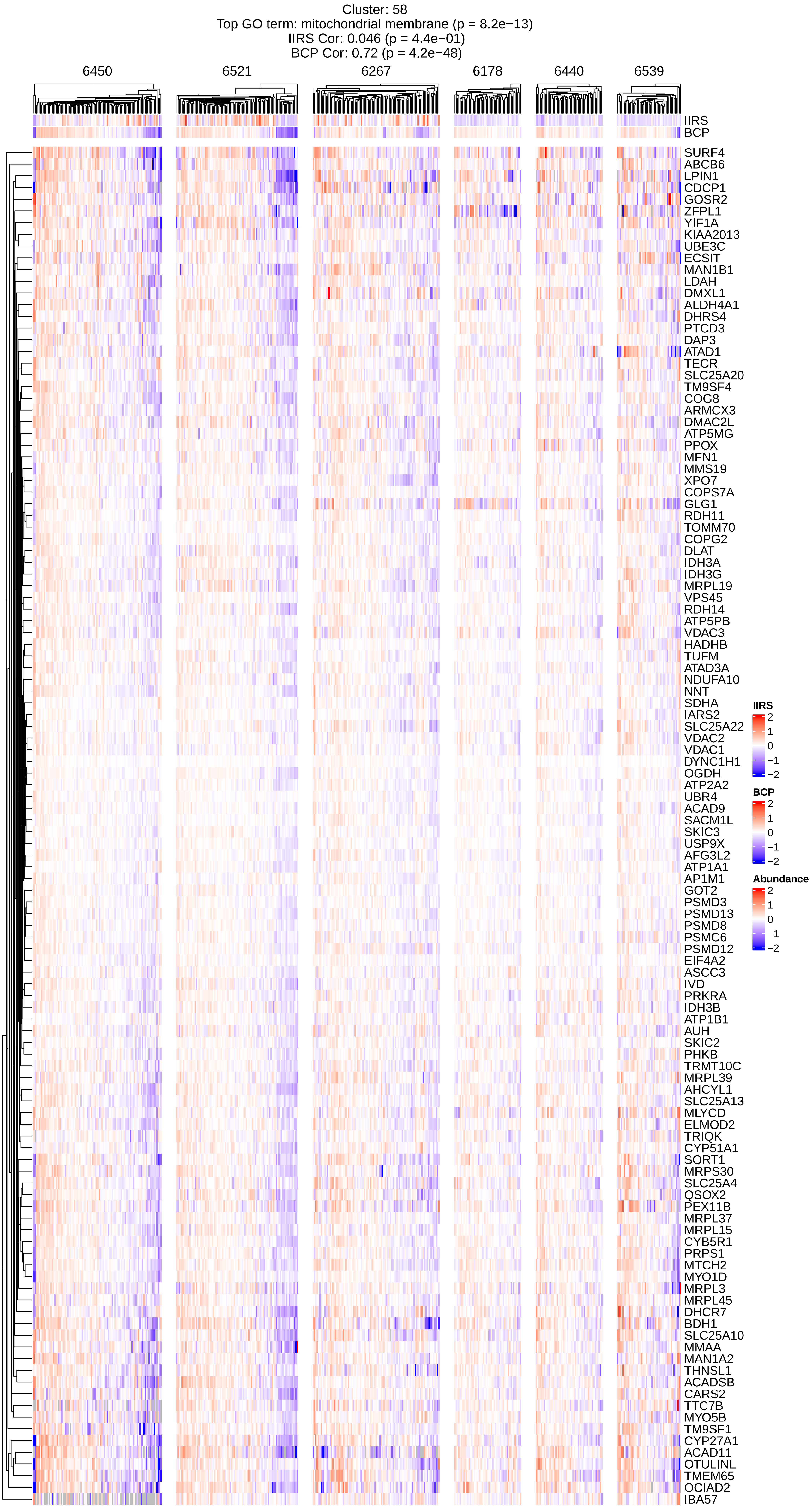
Cluster: 58
Top GO term: mitochondrial membrane (p = 8.2e−13)
IIRS Cor: 0.046 (p = 4.4e−01)
BCP Cor: 0.72 (p = 4.2e−48)
6450
6521
6267
6178
6440
6539
IIRS
BCP
SURF4
ABCB6
LPIN1
CDCP1
GOSR2
ZFPL1
YIF1A
KIAA2013
UBE3C
ECSIT
MAN1B1
LDAH
DMXL1
ALDH4A1
DHRS4
PTCD3
DAP3
ATAD1
TECR
SLC25A20
TM9SF4
COG8
ARMCX3
DMAC2L
ATP5MG
PPOX
MFN1
MMS19
XPO7
COPS7A
GLG1
RDH11
TOMM70
COPG2
DLAT
IDH3A
IDH3G
MRPL19
VPS45
RDH14
ATP5PB
VDAC3
HADHB
TUFM
ATAD3A
NDUFA10
NNT
SDHA
IARS2
SLC25A22
VDAC2
VDAC1
DYNC1H1
OGDH
ATP2A2
UBR4
ACAD9
SACM1L
SKIC3
USP9X
AFG3L2
ATP1A1
AP1M1
GOT2
PSMD3
PSMD13
PSMD8
PSMC6
PSMD12
EIF4A2
ASCC3
IVD
PRKRA
IDH3B
ATP1B1
AUH
SKIC2
PHKB
TRMT10C
MRPL39
AHCYL1
SLC25A13
MLYCD
ELMOD2
TRIQK
CYP51A1
SORT1
MRPS30
SLC25A4
QSOX2
PEX11B
MRPL37
MRPL15
CYB5R1
PRPS1
MTCH2
MYO1D
MRPL3
MRPL45
DHCR7
BDH1
SLC25A10
MMAA
MAN1A2
THNSL1
ACADSB
CARS2
TTC7B
MYO5B
TM9SF1
CYP27A1
ACAD11
OTULINL
TMEM65
OCIAD2
IBA57
IIRS
2
1
0
−1
−2
BCP
2
1
0
−1
−2
Abundance
2
1
0
−1
−2

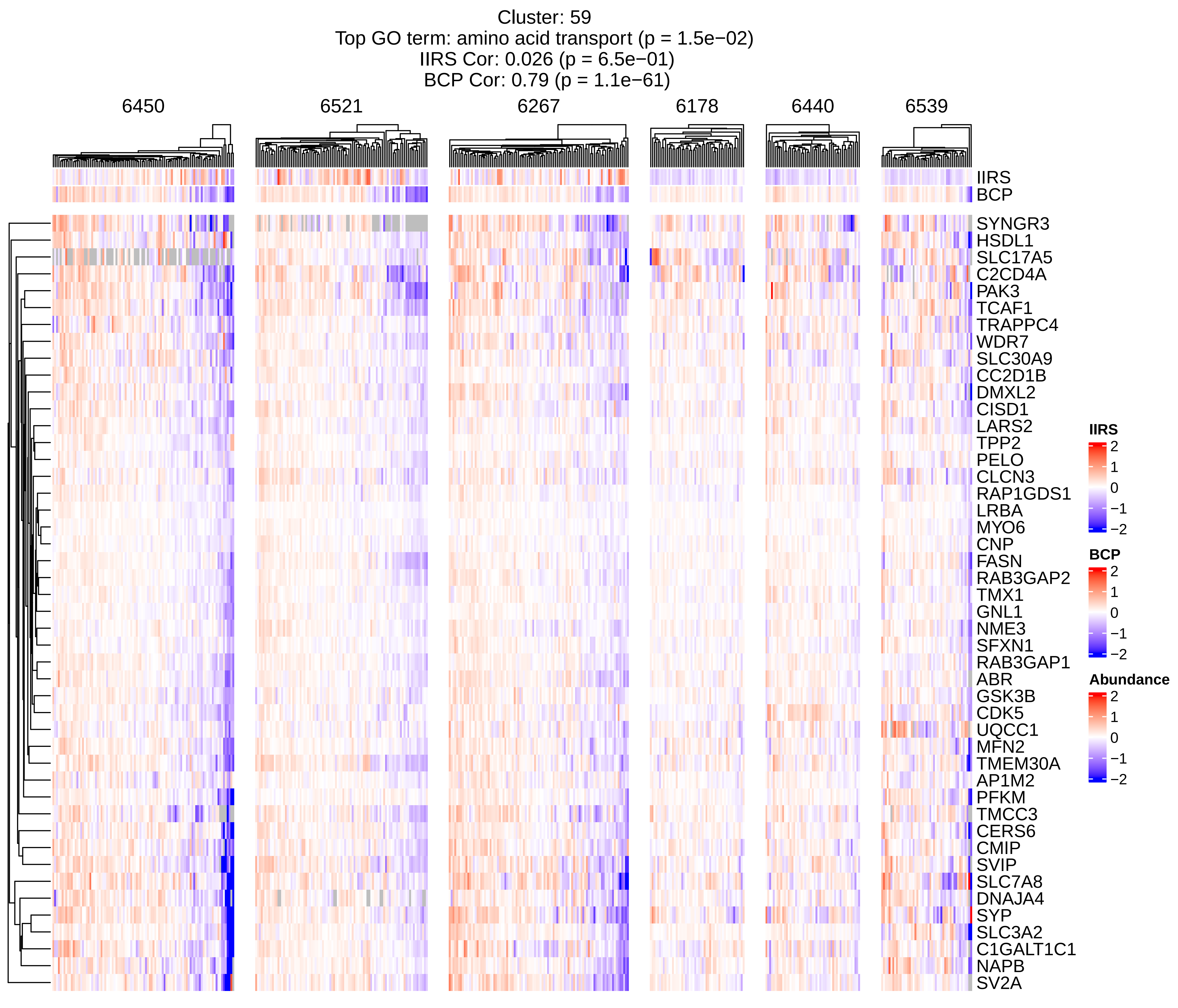

Cluster: 59
Top GO term: amino acid transport (p = 1.5e−02)
IIRS Cor: 0.026 (p = 6.5e−01)
BCP Cor: 0.79 (p = 1.1e−61)
6450
6521
6267
6178
6440
6539
IIRS
BCP
SYNGR3
HSDL1
SLC17A5
C2CD4A
PAK3
TCAF1
TRAPPC4
WDR7
SLC30A9
CC2D1B
DMXL2
CISD1
LARS2
TPP2
PELO
CLCN3
RAP1GDS1
LRBA
MYO6
CNP
FASN
RAB3GAP2
TMX1
GNL1
NME3
SFXN1
RAB3GAP1
ABR
GSK3B
CDK5
UQCC1
MFN2
TMEM30A
AP1M2
PFKM
TMCC3
CERS6
CMIP
SVIP
SLC7A8
DNAJA4
SYP
SLC3A2
C1GALT1C1
NAPB
SV2A
IIRS
2
1
0
−1
−2
BCP
2
1
0
−1
−2
Abundance
2
1
0
−1
−2

Cluster: 60
Top GO term: organelle membrane (p = 1.5e−09)
IIRS Cor: 0.04 (p = 5e−01)
BCP Cor: 0.95 (p = 3.9e−141)

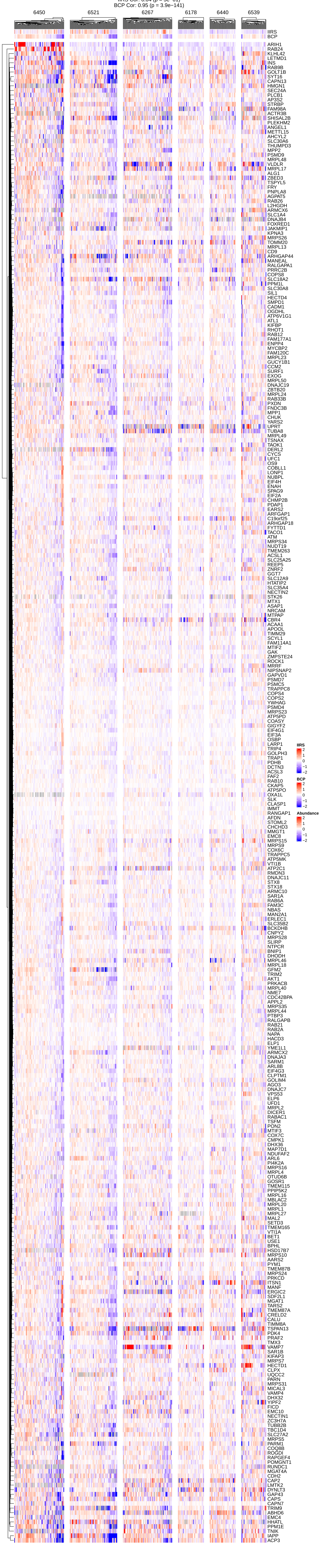

Cluster: 61
Top GO term: NS (p = NS)
IIRS Cor: −0.076 (p = 2e−01)
BCP Cor: 0.58 (p = 3e−27)

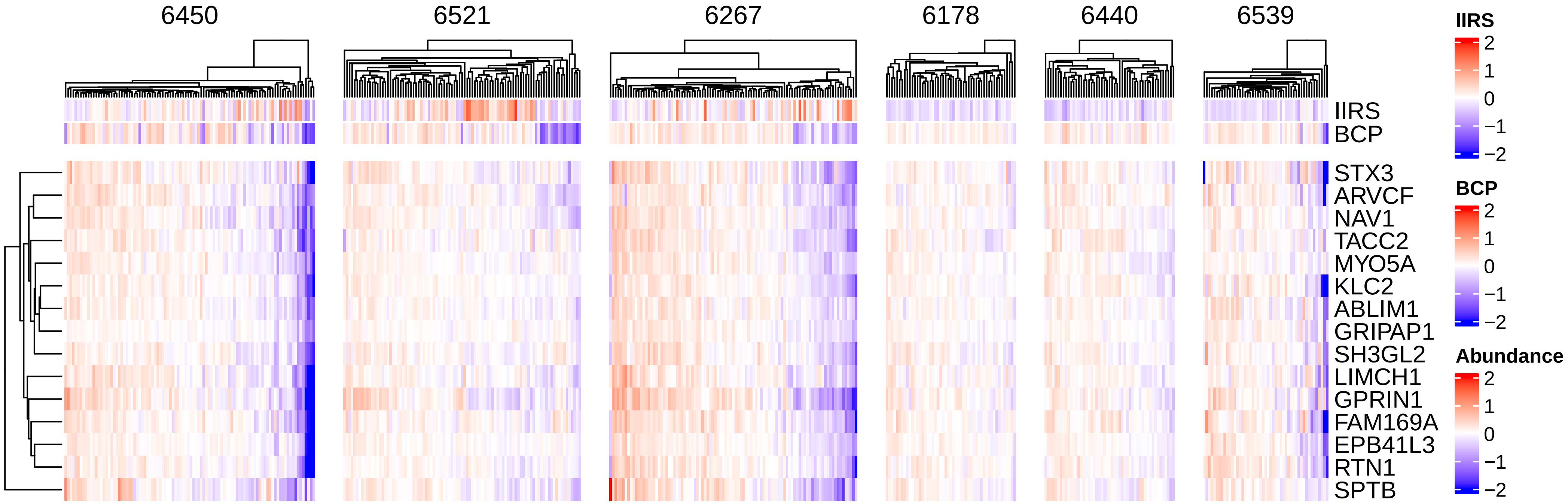

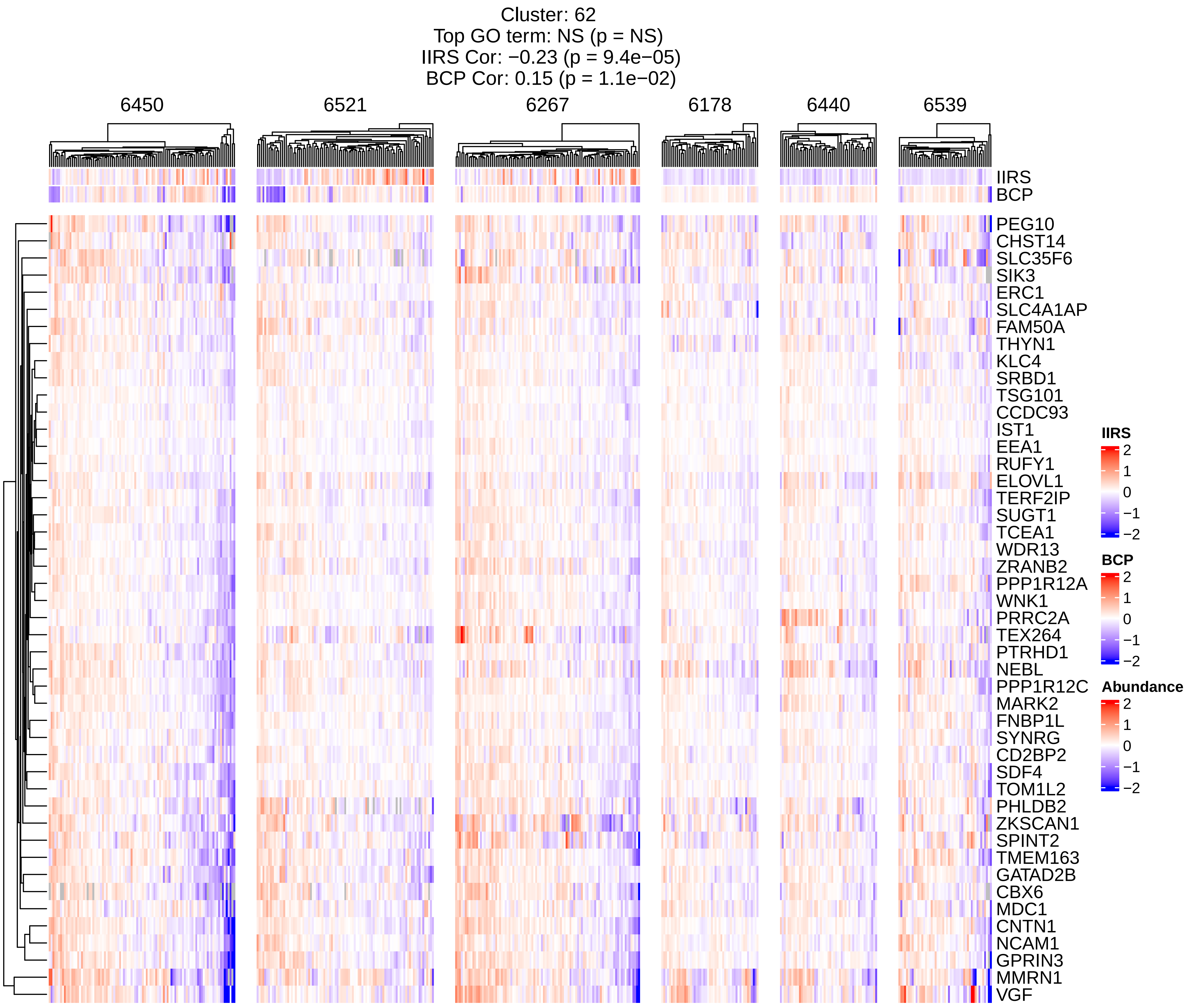

Cluster: 62
Top GO term: NS (p = NS)
IIRS Cor: −0.23 (p = 9.4e−05)
BCP Cor: 0.15 (p = 1.1e−02)
6450
6521
6267
6178
6440
6539
IIRS
BCP
PEG10
CHST14
SLC35F6
SIK3
ERC1
SLC4A1AP
FAM50A
THYN1
KLC4
SRBD1
TSG101
CCDC93
IST1
EEA1
RUFY1
ELOVL1
TERF2IP
SUGT1
TCEA1
WDR13
ZRANB2
PPP1R12A
WNK1
PRRC2A
TEX264
PTRHD1
NEBL
PPP1R12C
MARK2
FNBP1L
SYNRG
CD2BP2
SDF4
TOM1L2
PHLDB2
ZKSCAN1
SPINT2
TMEM163
GATAD2B
CBX6
MDC1
CNTN1
NCAM1
GPRIN3
MMRN1
VGF
IIRS
2
1
0
−1
−2
BCP
2
1
0
−1
−2
Abundance
2
1
0
−1
−2

Cluster: 63
Top GO term: vesicle localization (p = 3.7e−05)
IIRS Cor: −0.073 (p = 2.2e−01)
BCP Cor: 0.64 (p = 2.5e−34)

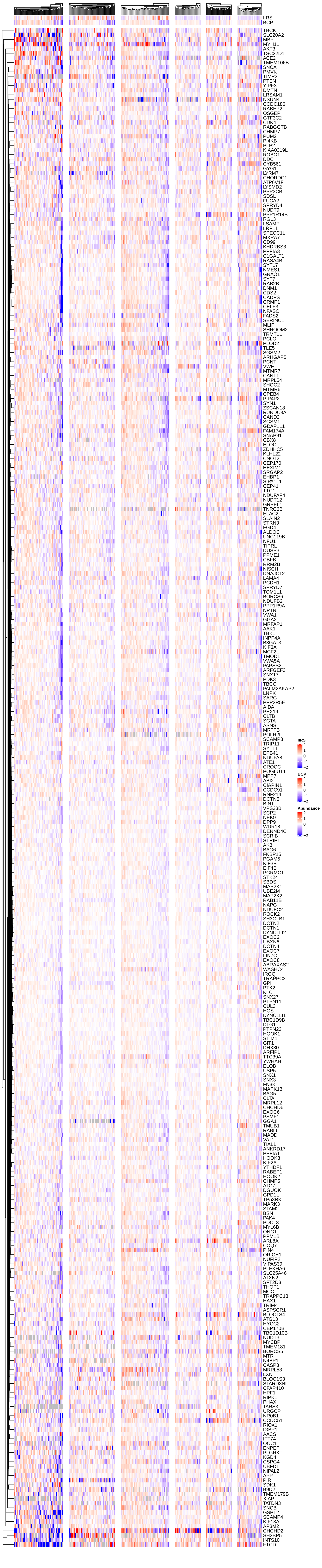

Cluster: 64
Top GO term: extracellular matrix structural constituent conferring compression resistance
(p = 1.1e−03)
IIRS Cor: 0.3 (p = 2.9e−07)
BCP Cor: 0.34 (p = 3.7e−09)

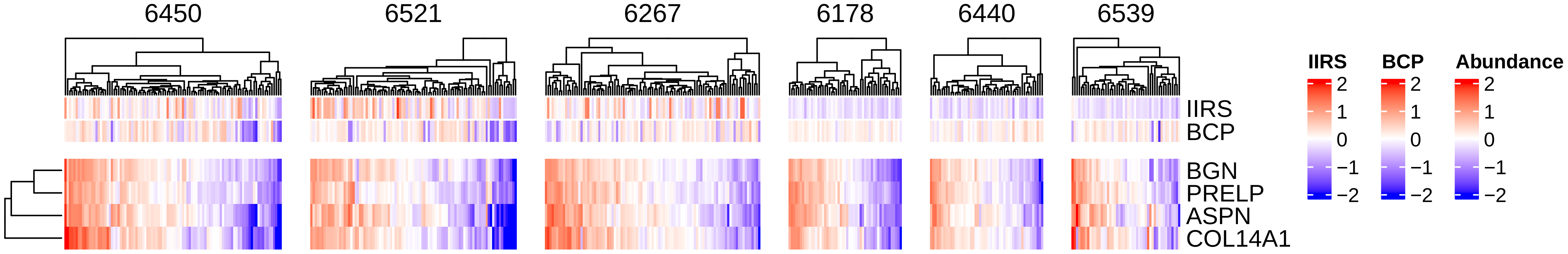

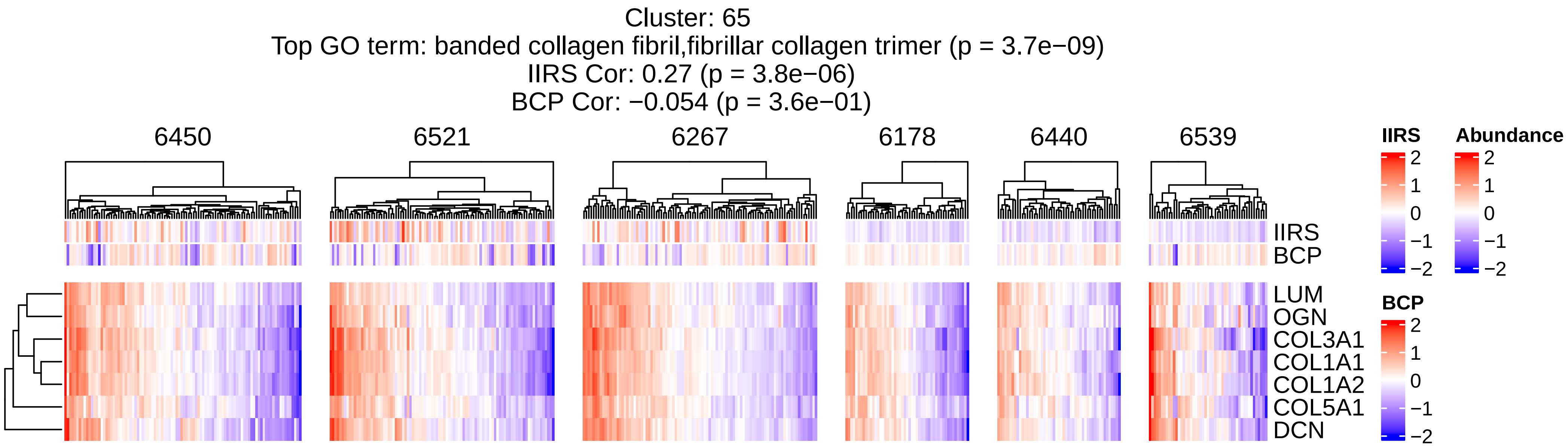

Cluster: 65
Top GO term: banded collagen fibril,fibrillar collagen trimer (p = 3.7e−09)
IIRS Cor: 0.27 (p = 3.8e−06)
BCP Cor: −0.054 (p = 3.6e−01)
6450
6521
6267
6178
6440
6539
IIRS
BCP
LUM
OGN
COL3A1
COL1A1
COL1A2
COL5A1
DCN
IIRS
2
1
0
−1
−2
Abundance
2
1
0
−1
−2
BCP
2
1
0
−1
−2

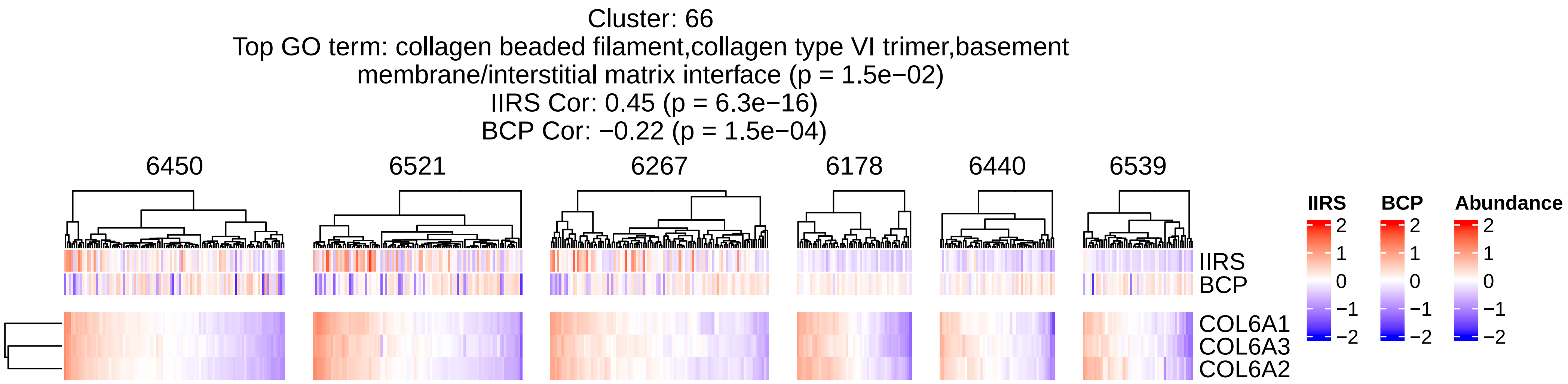

Cluster: 66
Top GO term: collagen beaded filament,collagen type VI trimer,basement membrane/interstitial matrix interface (p = 1.5e−02)
IIRS Cor: 0.45 (p = 6.3e−16)
BCP Cor: −0.22 (p = 1.5e−04)
6450
6521
6267
6178
6440
6539
IIRS
BCP
COL6A1
COL6A3
COL6A2
IIRS
BCP
Abundance
2
1
0
−1
−2

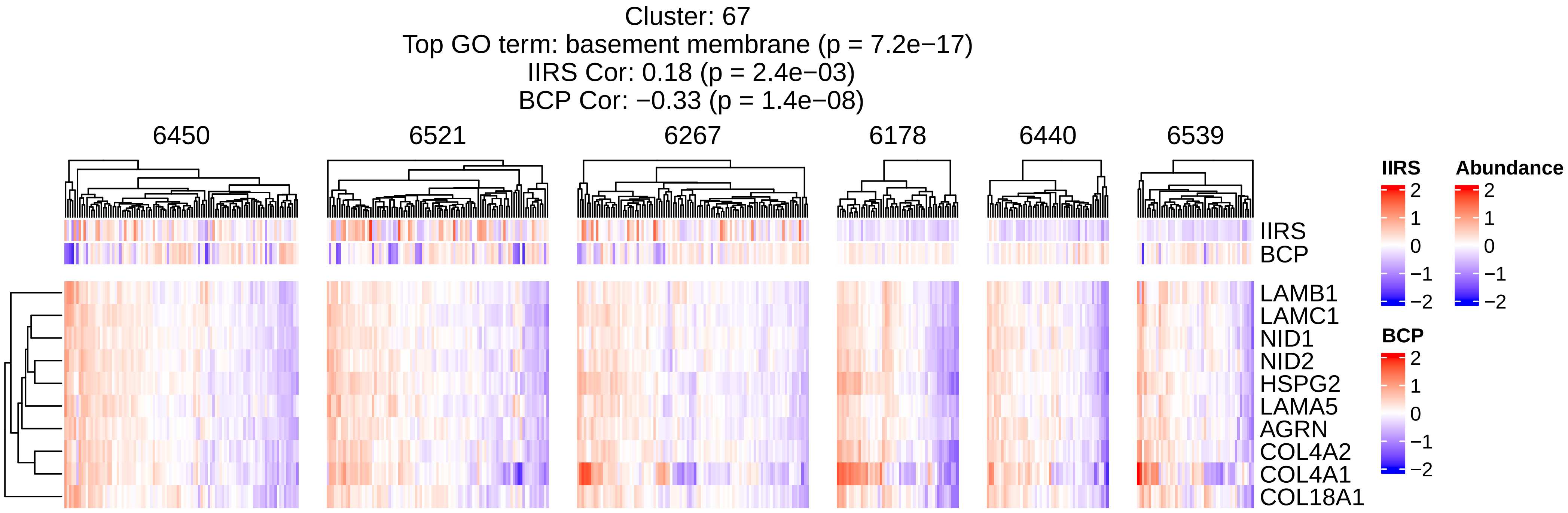
Cluster: 67
Top GO term: basement membrane (p = 7.2e−17)
IIRS Cor: 0.18 (p = 2.4e−03)
BCP Cor: −0.33 (p = 1.4e−08)
6450
6521
6267
6178
6440
6539
IIRS
BCP
LAMB1
LAMC1
NID1
NID2
HSPG2
LAMA5
AGRN
COL4A2
COL4A1
COL18A1
IIRS
2
1
0
−1
−2
Abundance
2
1
0
−1
−2
BCP
2
1
0
−1
−2

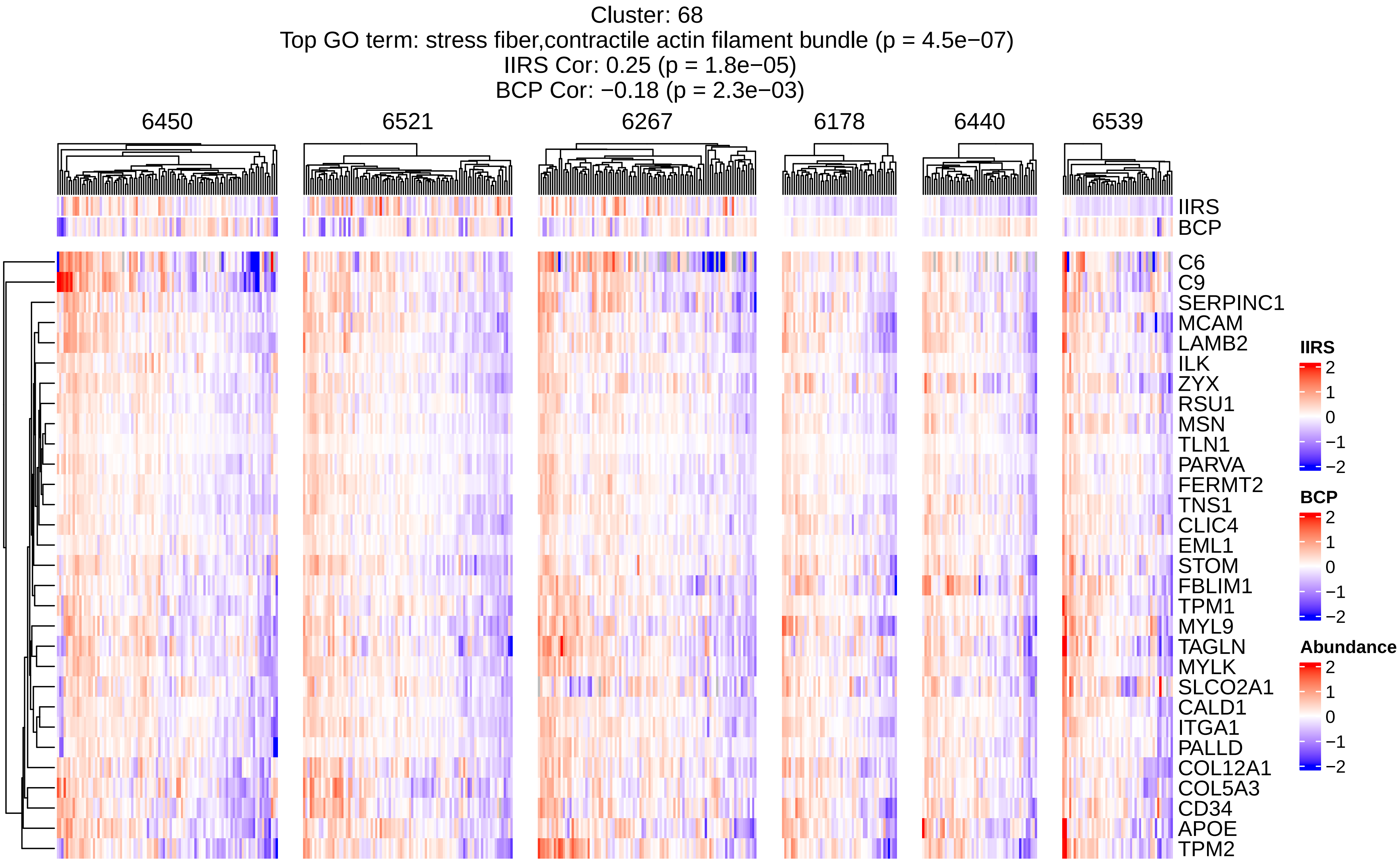
Cluster: 68
Top GO term: stress fiber,contractile actin filament bundle (p = 4.5e−07)
IIRS Cor: 0.25 (p = 1.8e−05)
BCP Cor: −0.18 (p = 2.3e−03)
6450
6521
6267
6178
6440
6539
IIRS
BCP
C6
C9
SERPINC1
MCAM
LAMB2
ILK
ZYX
RSU1
MSN
TLN1
PARVA
FERMT2
TNS1
CLIC4
EML1
STOM
FBLIM1
TPM1
MYL9
TAGLN
MYLK
SLCO2A1
CALD1
ITGA1
PALLD
COL12A1
COL5A3
CD34
APOE
TPM2
IIRS
2
1
0
−1
−2
BCP
2
1
0
−1
−2
Abundance
2
1
0
−1
−2

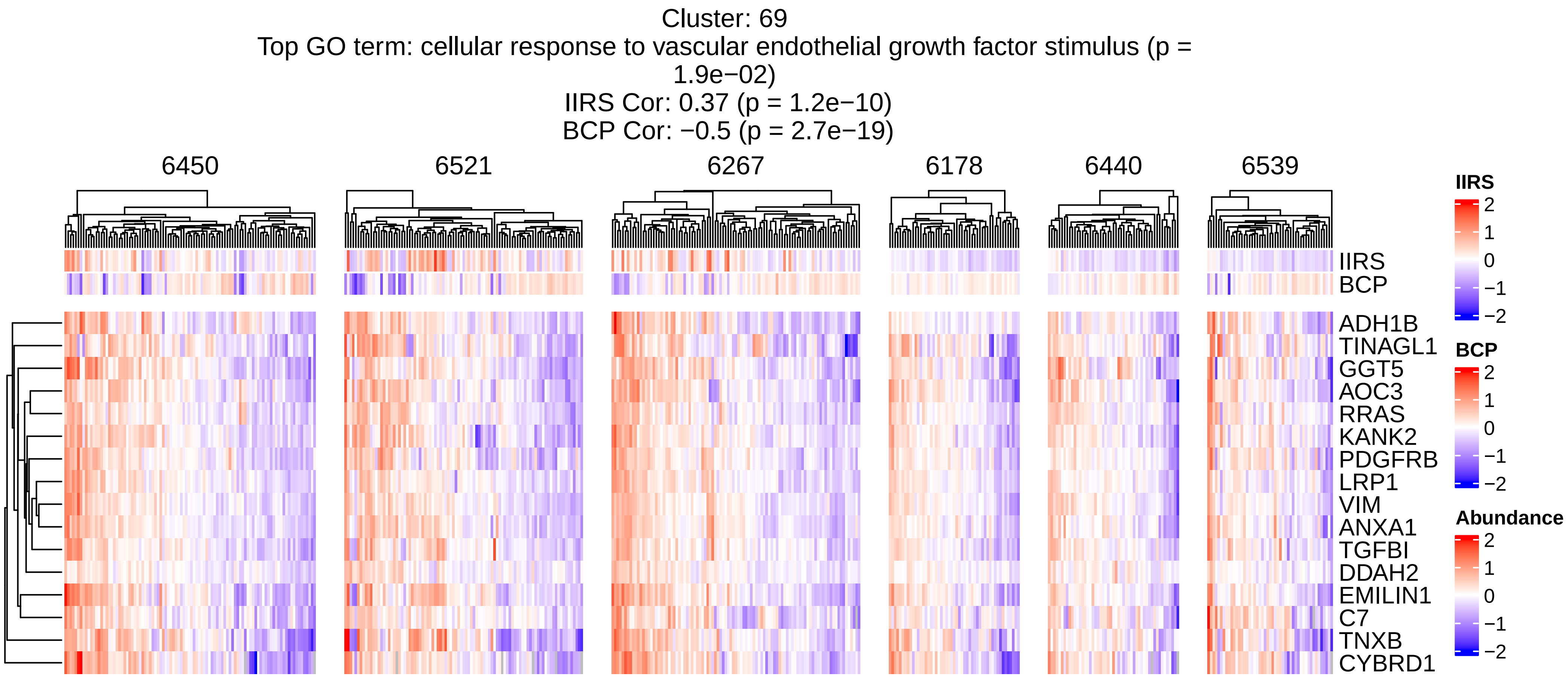
Cluster: 69
Top GO term: cellular response to vascular endothelial growth factor stimulus (p = 1.9e−02)
IIRS Cor: 0.37 (p = 1.2e−10)
BCP Cor: −0.5 (p = 2.7e−19)
6450
6521
6267
6178
6440
6539
IIRS
BCP
ADH1B
TINAGL1
GGT5
AOC3
RRAS
KANK2
PDGFRB
LRP1
VIM
ANXA1
TGFBI
DDAH2
EMILIN1
C7
TNXB
CYBRD1
IIRS
2
1
0
−1
−2
BCP
2
1
0
−1
−2
Abundance
2
1
0
−1
−2

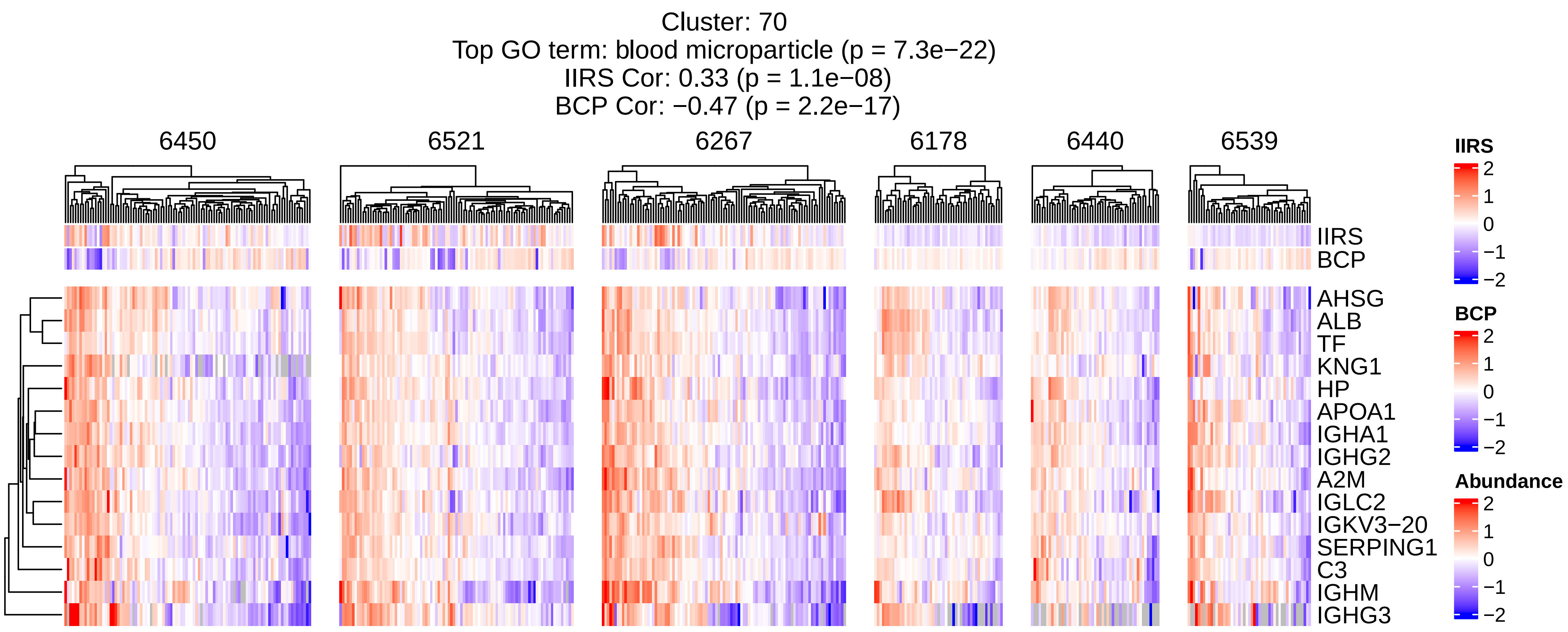

Cluster: 70
Top GO term: blood microparticle (p = 7.3e−22)
IIRS Cor: 0.33 (p = 1.1e−08)
BCP Cor: −0.47 (p = 2.2e−17)
6450
6521
6267
6178
6440
6539
IIRS
BCP
AHSG
ALB
TF
KNG1
HP
APOA1
IGHA1
IGHG2
A2M
IGLC2
IGKV3−20
SERPING1
C3
IGHM
IGHG3
IIRS
2
1
0
−1
−2
BCP
2
1
0
−1
−2
Abundance
2
1
0
−1
−2

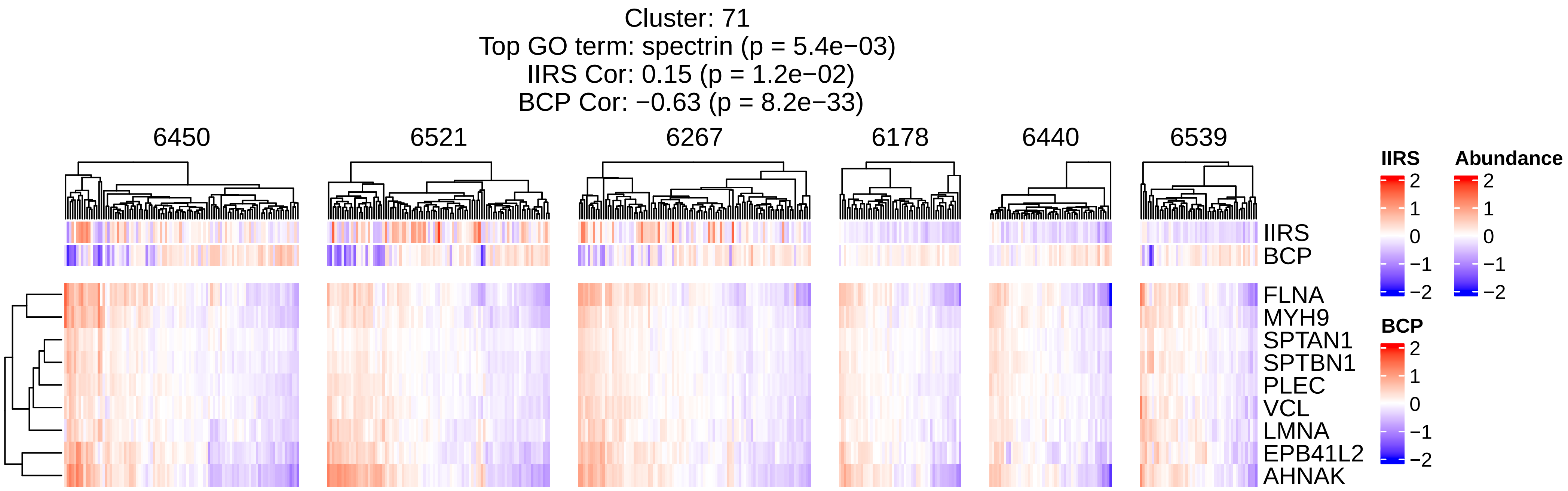
Cluster: 71
Top GO term: spectrin (p = 5.4e−03)
IIRS Cor: 0.15 (p = 1.2e−02)
BCP Cor: −0.63 (p = 8.2e−33)
6450
6521
6267
6178
6440
6539
IIRS
BCP
FLNA
MYH9
SPTAN1
SPTBN1
PLEC
VCL
LMNA
EPB41L2
AHNAK
IIRS
2
1
0
−1
−2
Abundance
2
1
0
−1
−2
BCP
2
1
0
−1
−2

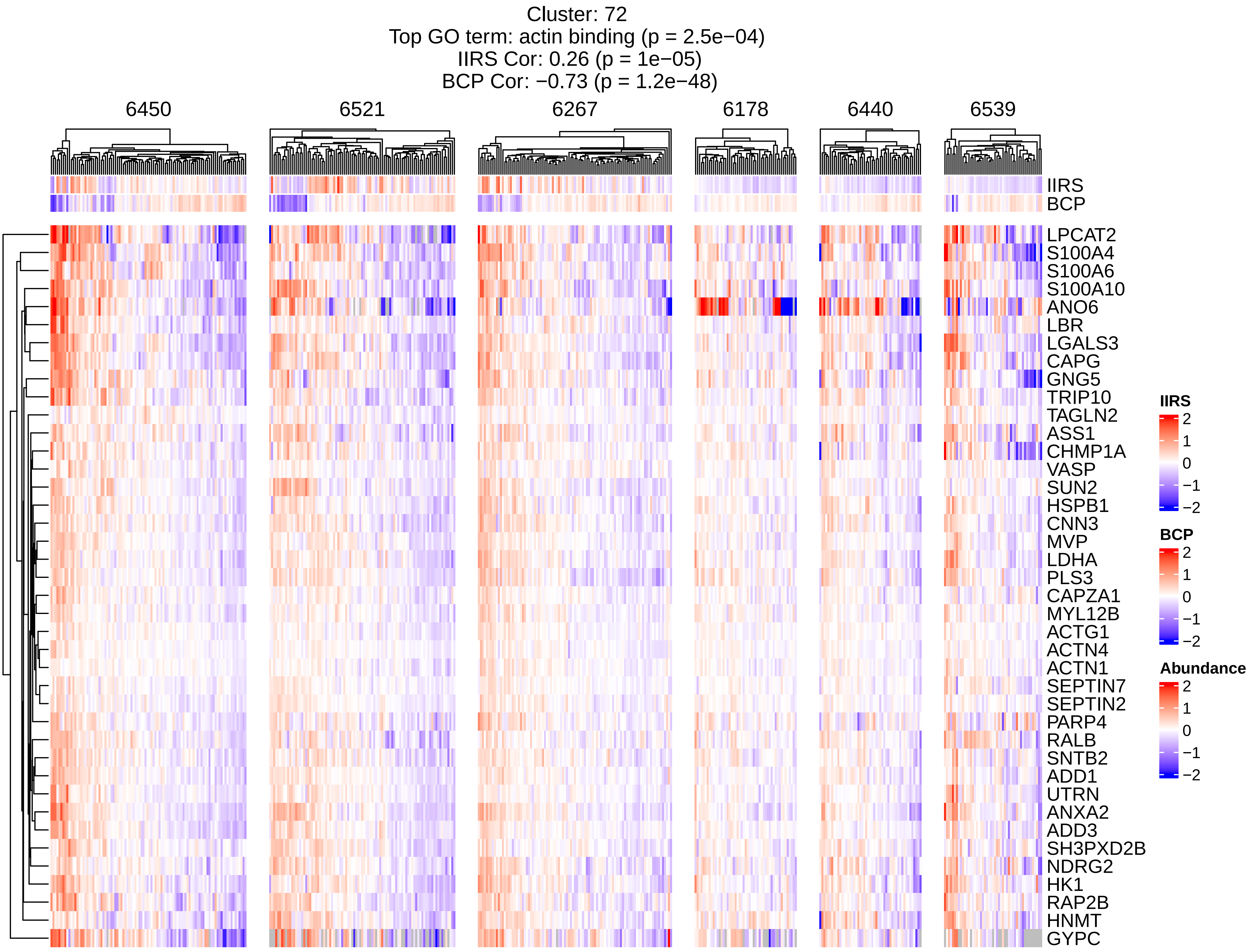
Cluster: 72
Top GO term: actin binding (p = 2.5e−04)
IIRS Cor: 0.26 (p = 1e−05)
BCP Cor: −0.73 (p = 1.2e−48)
6450
6521
6267
6178
6440
6539
IIRS
BCP
LPCAT2
S100A4
S100A6
S100A10
ANO6
LBR
LGALS3
CAPG
GNG5
TRIP10
TAGLN2
ASS1
CHMP1A
VASP
SUN2
HSPB1
CNN3
MVP
LDHA
PLS3
CAPZA1
MYL12B
ACTG1
ACTN4
ACTN1
SEPTIN7
SEPTIN2
PARP4
RALB
SNTB2
ADD1
UTRN
ANXA2
ADD3
SH3PXD2B
NDRG2
HK1
RAP2B
HNMT
GYPC
IIRS
2
1
0
−1
−2
BCP
2
1
0
−1
−2
Abundance
2
1
0
−1
−2

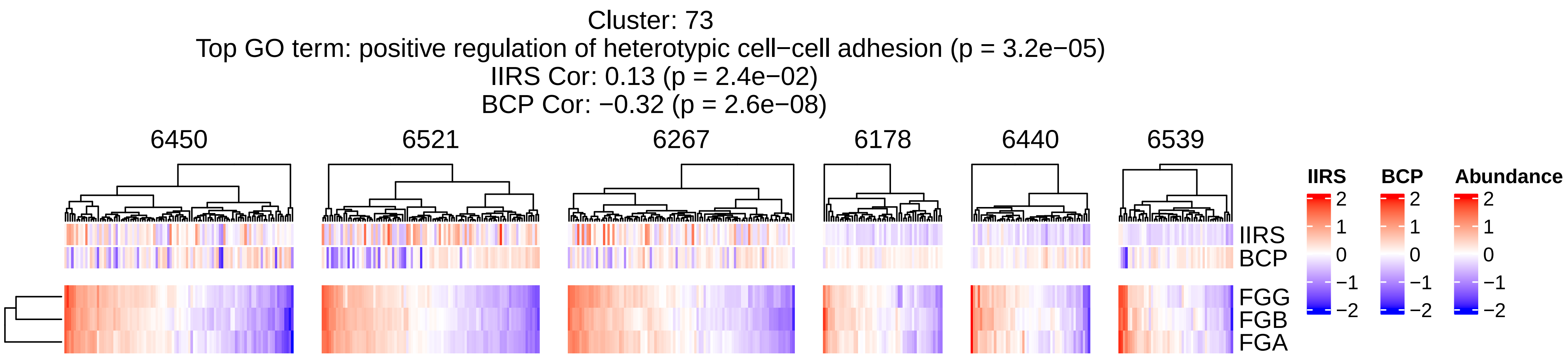
Cluster: 73
Top GO term: positive regulation of heterotypic cell−cell adhesion (p = 3.2e−05)
IIRS Cor: 0.13 (p = 2.4e−02)
BCP Cor: −0.32 (p = 2.6e−08)
6450
6521
6267
6178
6440
6539
IIRS
BCP
FGG
FGB
FGA
IIRS
BCP
Abundance
2
1
0
−1
−2

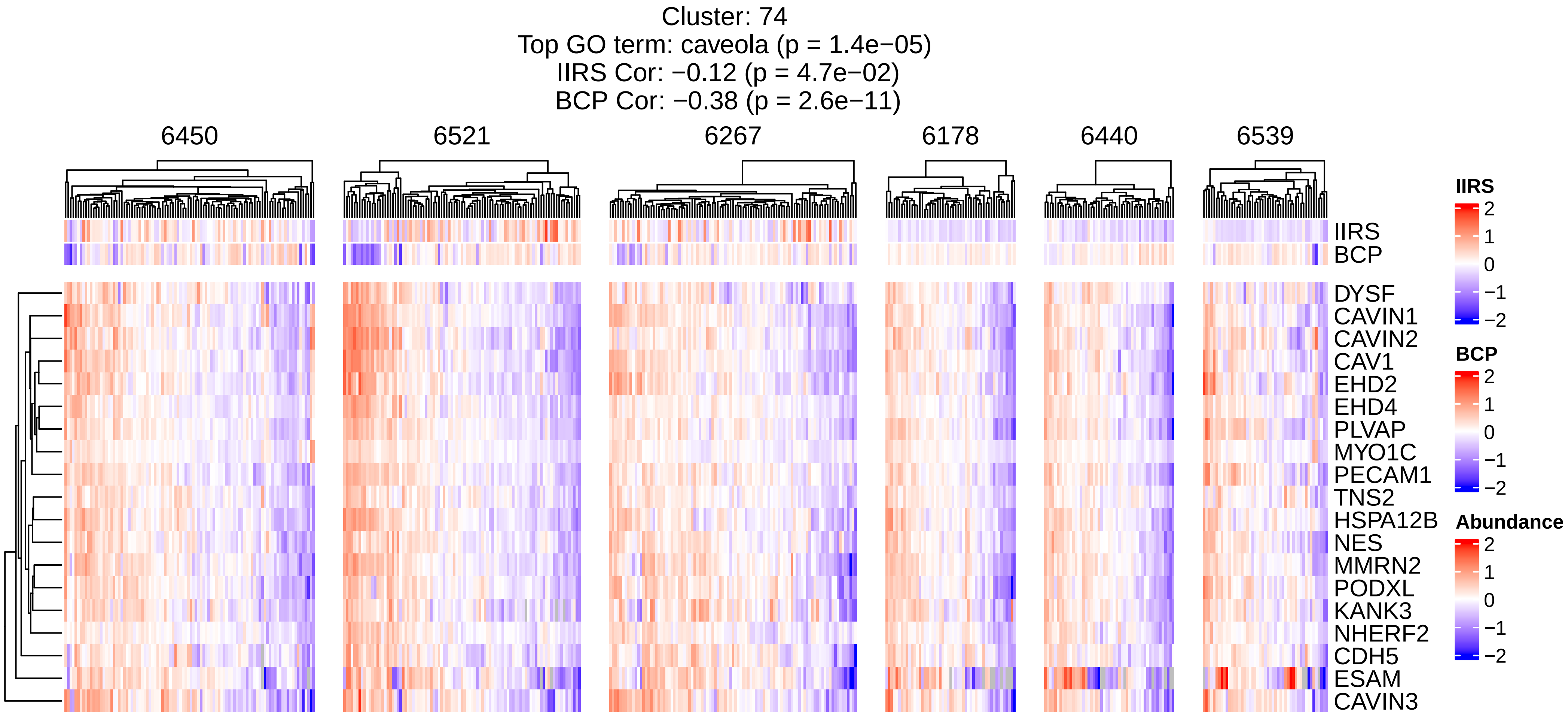

Cluster: 74
Top GO term: caveola (p = 1.4e−05)
IIRS Cor: −0.12 (p = 4.7e−02)
BCP Cor: −0.38 (p = 2.6e−11)
6450
6521
6267
6178
6440
6539
IIRS
BCP
DYSF
CAVIN1
CAVIN2
CAV1
EHD2
EHD4
PLVAP
MYO1C
PECAM1
TNS2
HSPA12B
NES
MMRN2
PODXL
KANK3
NHERF2
CDH5
ESAM
CAVIN3
IIRS
2
1
0
−1
−2
BCP
2
1
0
−1
−2
Abundance
2
1
0
−1
−2

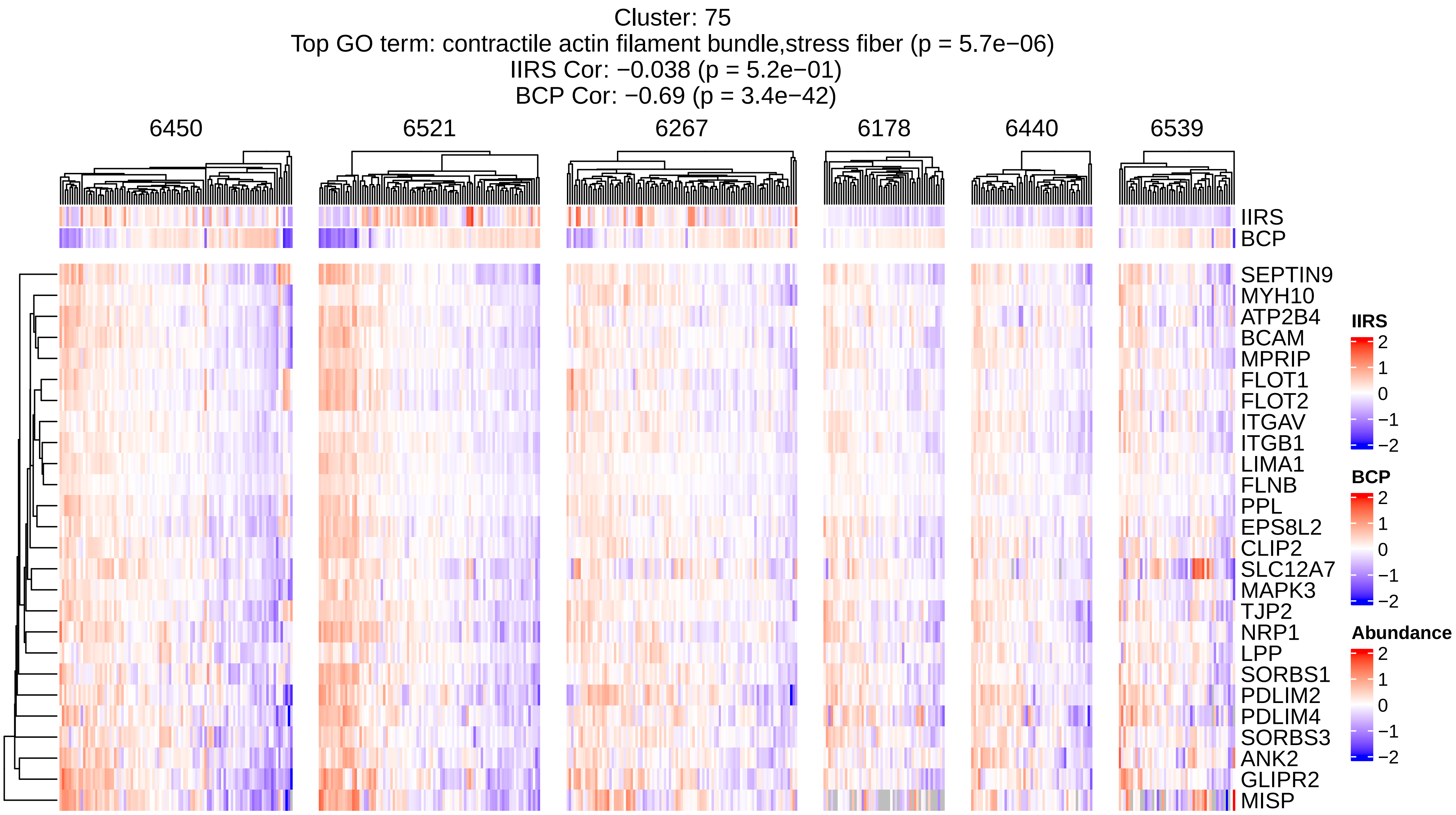
Cluster: 75
Top GO term: contractile actin filament bundle,stress fiber (p = 5.7e−06)
IIRS Cor: −0.038 (p = 5.2e−01)
BCP Cor: −0.69 (p = 3.4e−42)
6450
6521
6267
6178
6440
6539
IIRS
BCP
SEPTIN9
MYH10
ATP2B4
BCAM
MPRIP
FLOT1
FLOT2
ITGAV
ITGB1
LIMA1
FLNB
PPL
EPS8L2
CLIP2
SLC12A7
MAPK3
TJP2
NRP1
LPP
SORBS1
PDLIM2
PDLIM4
SORBS3
ANK2
GLIPR2
MISP
IIRS
2
1
0
−1
−2
BCP
2
1
0
−1
−2
Abundance
2
1
0
−1
−2
